# Neuronal IL-17 controls *C. elegans* developmental diapause through CEP-1/p53

**DOI:** 10.1101/2022.11.22.517560

**Authors:** Abhishiktha Godthi, Sehee Min, Srijit Das, Johnny Cruz-Corchado, Andrew Deonarine, Kara Misel-Wuchter, Priya D. Issuree, Veena Prahlad

## Abstract

During metazoan development, how cell division and metabolic programs are coordinated with nutrient availability remains unclear. Here, we show that nutrient availability signaled by the neuronal cytokine, ILC-17.1 switches *C. elegans* development between reproductive growth and dormancy by controlling the activity of the tumor suppressor p53 ortholog, CEP-1. Specifically, upon food availability, ILC-17.1 signaling by amphid neurons promotes glucose utilization and suppresses CEP-1/p53 to allow growth. In the absence of ILC-17.1, CEP-1/p53 is activated, upregulates cell-cycle inhibitors, decreases phosphofructokinase and cytochrome C expression, and causes larvae to arrest as stress-resistant, quiescent dauers. We propose a model whereby ILC-17.1 signaling links nutrient availability and energy metabolism to cell cycle progression through CEP-1/p53. These studies describe ancestral functions of IL-17s and the p53-family of proteins and are relevant to our understanding of neuroimmune mechanisms in cancer. They also reveal a DNA damage-independent function of CEP-1/p53 in invertebrate development and support the existence of a previously undescribed *C. elegans* dauer pathway.

During metazoan development, nutrient availability is coordinated with the division, growth and metabolic activity of individual cells through cell-cell communication. This is also the case in the invertebrate *C. elegans*, a free-living bacterivore, which displays a dramatic developmental plasticity to ensure that its growth and reproduction match available resources(1–10). When *C. elegans* larvae hatch under optimal conditions (at 20°C, low population densities, on abundant food) they develop continuously into reproducing adults. However, if they hatch under suboptimal conditions, such as in the paucity of food, at high population densities, or high ambient temperatures, larvae implement an alternative developmental program and arrest as quiescent, stress-resistant larvae called ‘dauer‘ larvae. Dauer larvae display metabolic and organismal phenotypes specialized for dispersal and survival, and can remain arrested in this state for months to resume development into reproductive adults when favorable conditions return(1–10). Previous studies have identified molecular pathways that mediate the dauer decision, showing that growth promoting molecules like insulins, transforming β growth factor (TGFβ/DAF-7) and lipid based dafachronic acid hormones are released by sensory neurons and other cells to license continued development; adverse environments inhibit these growth promoting signals and trigger dauer arrest(1–11). A number of quantitative trait loci (QTL) also modulate dauer (12). Yet, how the dauer entry decision results in a coordinated change in cell fates across different tissues and is linked with the systemic shut-down of anabolic pathways remains poorly understood.

An important group of proteins that mediate cell-cell communication and metabolism in metazoa are secreted proteins called cytokines(13, 14). The IL-17 cytokines constitute a family of proinflammatory cytokines, highly conserved across animal phyla. In mammals, these cytokines are released by specialized immune cells to activate immune surveillance, enhance barrier function, promote wound healing, and play crucial immunometabolic roles in maintaining energy homeostasis(15). In humans, IL-17s also promote cancers and autoimmune disease such as psoriasis(16, 17). Here, we show that the *C. elegans* IL-17 ortholog, ILC-17.1, signals food availability, and coordinates cell division with metabolism by controlling the activity of the *C. elegans* tumor suppressor p53 ortholog, CEP-1. Specifically, neuronal ILC-17.1 suppresses CEP-1/p53 activity in the presence of food to license growth. Upon the loss of ILC-17.1 signaling, CEP-1/p53 is activated, and remarkably, this switches whole organism development from continuous growth to dormancy. The p53-like tumor suppressor genes are found in all multicellular animals where they prevent the transmission of damaged DNA by activating a multifaceted program that controls cell cycle checkpoints, mediates reversible growth arrest or apoptosis, and controls metabolic flux (18–22). Our studies show that these functions of CEP-1/p53 also act, in the absence of DNA damage, to control developmental quiescence of *C. elegans*, suggesting that the developmental function of the p53-gene family could have shaped their evolution(23–25).

**Significance:** Development in a metazoan requires that the division and differentiation of diverse cells be coordinated with nutrient availability. We show that one mechanism by which this occurs in *C. elegans* is through signaling by the neuronal IL-17 cytokine, ILC-17.1, and its control over p53/CEP-1. In the presence of food, ILC-17.1 release suppresses p53/CEP-1 and allows reproductive growth; decreased ILC-17.1 signaling activates p53/CEP-1-dependent transcription and metabolic programs, leading to the reversible arrest of larvae as quiescent dauers. Our studies suggest an ancestral function of IL-17 is linking nutrient availability to energy metabolism and growth. They reveal a DNA damage-independent function of p53/CEP-1 in invertebrate development. Finally, our studies support the existence of a previously undescribed dauer pathway in *C. elegans*.

## Results

### ILC-17.1 is released by *C. elegans* amphid neurons in response to food and prevents dauer arrest

*C. elegans* express three IL-17 orthologs(26, 27). Of these, ILC-17.1 is a neuromodulator that is expressed by a subset of specialized sensory neurons with epithelial properties called amphid neurons, that are exposed to the environment(26–28). We discovered that a deletion in *ilc-17.1, syb5296,* that removes almost all the coding sequence (2173bp out of 2980 bp; Supplementary Fig. S1a) and abolishes mRNA expression (Supplementary Fig. S1b) caused larvae to constitutively enter the dauer state (Figure 1a, b; Supplementary Fig. S1c, d). Dauer larvae can be identified by their growth arrest and distinct morphology characterized by specialized cuticular structures called alae, a constricted pharynx, arrested germ line, decreased pharyngeal pumping rates, and detergent (1% SDS) resistance(3–9). *ilc-17.1*(*syb5296*)X dauer larvae displayed all these features (Supplementary Fig. S1e-j). Even at 20°C, *ilc-17.1* deleted larvae entered the dauer state transiently: approximately 30% (31.2 ±5%) were SDS-resistant 48 hours post-hatching on OP50 bacteria, and most larvae exited dauer and become reproductive adults by 72 hrs. (Supplementary Fig. S1c, d). Under the same conditions, none of the wild-type larvae were detergent resistant, nor did they arrest as dauers. However, as with other mutations that promote dauer entry(3, 4, 29), the dauer arrest of *ilc-17.1* deletion mutants was dramatically accentuated at the slightly higher ambient temperature of 25°C, which still supported the growth of all wild-type larvae into reproductive adults, but caused practically 100% of larvae lacking *ilc-17.1* to enter and persist in the arrested dauer state for days (Fig.1a-d).

**Figure 1:**
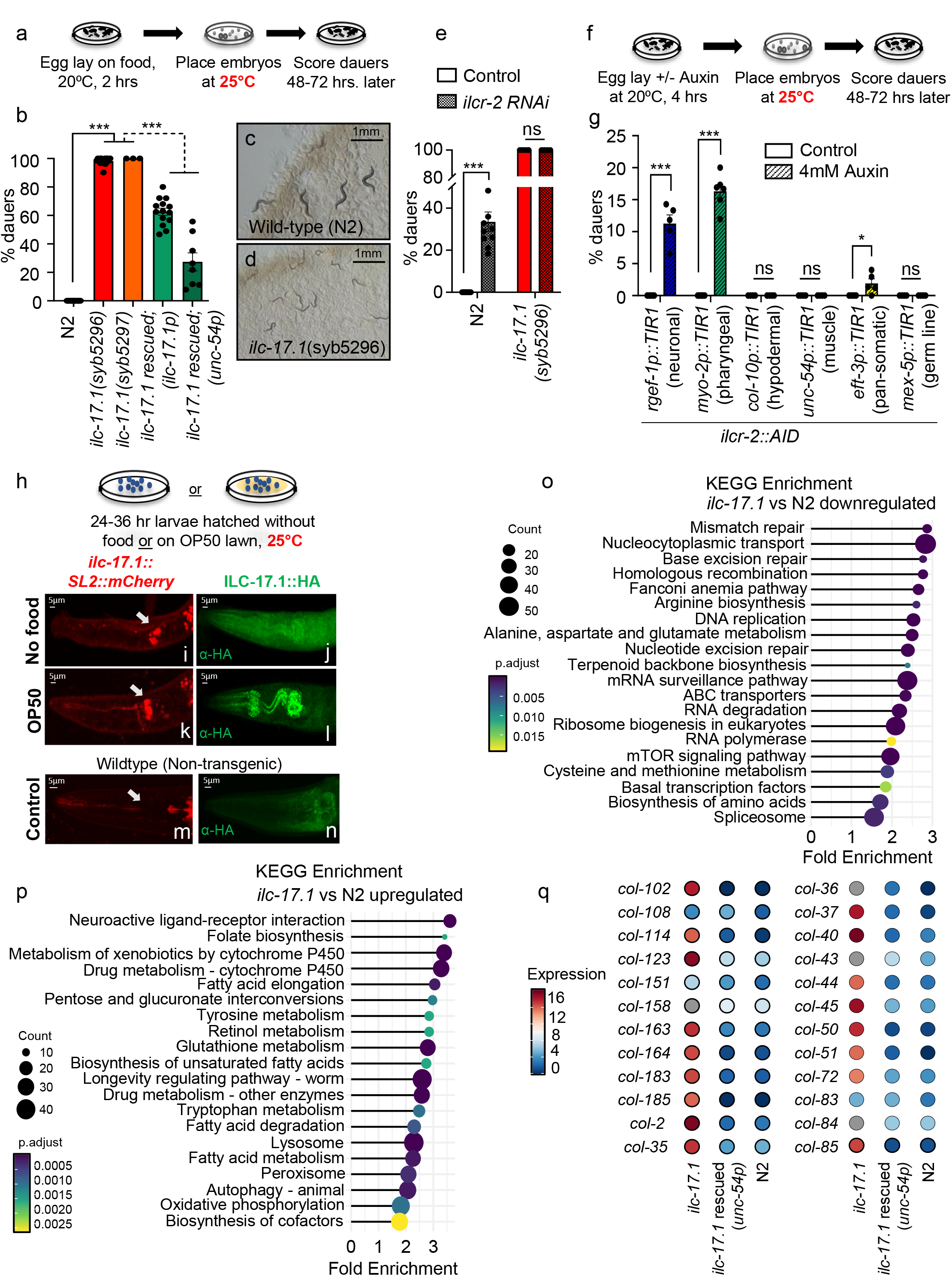
ILC-17.1 signaling of food availability by larval amphid neurons modulates dauer arrest. **(a),** Schematic of experiment. For each experiment, independent samples of 10 day-one adults of each strain laid eggs for two-four hours at 20°C on OP50 lawns. Adults were removed, embryos transferred to 25°C., larval phenotype scored after approximately 48 hr., and categorized as dauers or stage 4 larvae (L4) and/or young adults. SDS-resistance was used to confirm dauers. Extent of crowding on plates was comparable (Representative average numbers of embryos/plates are 64, 68, 56, 46 and 65 for wild type (N2), *ilc-17.1*(*syb5296*) X, *ilc-17.1*(*syb5297*) X, *ilc-17.1*(*syb5296*) rescued with *ilc-17.1p::ilc-17.1*, and *ilc-17.1*(*syb5296*) rescued with *unc-54p::ilc-17.1* respectively). *ilc-17.1 rescued (ilc-17.1p)* are *ilc-17.1* deletion mutants expressing extrachromosomal *ilc-17.1* cDNA driven by *ilc-17.1* promoter*. ilc-17.1* rescued *(unc-54p)* are *ilc-17.1* deletion mutants expressing extrachromosomal *ilc-17.1* cDNA driven by *unc-54* promoter. Pearson’s Chi-squared test with Yates’ continuity correction was used to compare samples in manuscript, unless otherwise stated. Chi-squared, df and p values for the whole experiment and p values for pair-wise comparisons are reported. ***p < 0.001, **p < 0.01, *p < 0.05 and ns, not significant. **(b),** Percentage larvae that arrest as dauers, (*n*=3-12 experiments; Chi-squared = 2359.6, df = 4, p-value < 2.2e-16.). (**c**, **d)**. Representative micrographs 48 hours post-hatching at 25°C: **(c),** wild type (N2) are L4s, **(d),** *ilc-17.1*(*syb5296*) are dauer. Scale bar, 1 mm. **(e)**. Percentage dauers amongst wild type (N2), and *ilc-17.1* mutants on control (L4440) and *ilcr-2* RNAi, (n=3-10 experiments. Chi-squared = 1815.1, df = 7, p-value < 2.2e-16.). **(f),** Schematic of AID experiment to degrade ILCR-2. **(g),** Percentage dauers following tissue specific degradation of ILCR-2. X-axis: tissue specific TIR1 expression, (n=4-6 experiments; Chi-squared = 335.56, df =9, p-value <2.2e-16). (**h).** Schematic of experiment to localize *ilc-17.1* mRNA and ILC-17.1 protein in the presence and absence of food (OP50). Embryos hatched at 20°C and imaged after 24-36 hours (L1 stage larvae): **(i, k),** Representative micrographs of z-section of pharyngeal region of larvae showing mCherry as proxy for *ilc-17.1* mRNA expression in amphid neurons**. (j, l),** z-section projection showing immunolocalization of ILC-17.1::HA protein. **(i, j)** larvae in absence of food, and **(k, l),** presence of food. **(m, n),** Controls (N2) that do not express HA or mCherry to indicate specificity of the fluorescence signal. (n>3 experiments, 4-5 larvae each). Scale bar, 5µm. (**o),** KEGG enrichments associated with differentially downregulated genes, and (**p)**, differentially upregulated genes in *ilc-17.1* mutant larvae. p.adjust <0.05. RNA-seq on 32-36 hr. larvae. (**q),** Heatmap depicting expression levels (log_2_ normalized counts) of the major dauer-specific collagens in L2 wild-type (N2) larvae *en route* to adult development, *ilc-17.1* deletion mutants *en route* to dauer entry and *ilc-17.1* deletion mutants rescued from dauer with *unc-54p::ilc-17.1.* Bars show mean ± S.E.M. Individual data points in the bar graphs in **b** and **e, g** represent the % dauers/experiment, and the bar graph depicts mean of these percentages. ***p < 0.001, **p < 0.01, *p < 0.05 and ns, not significant

*Ilc-17.1* has not been identified in previous genetic screens for dauer pathway genes(3, 5, 30–32). Therefore, although no off-target effects using CRISPR have been observed in *C. elegans* even after whole genome sequencing (33–35), we conducted additional experiments to confirm the role of *ilc-17.1* in dauer entry, by (i) examining the developmental phenotype of another independently generated CRISPR deletion in *ilc-17.1, syb5297* (loss of 2188 bp of 2980 bp; Supplementary Fig. S1a), (ii) backcrossing *ilc-17.1 (syb5296)* mutants, and (iii) downregulating *ilc-17.1* and its receptor(26, 27) in a wild-type (N2) background using RNA interference (RNAi) and auxin-induced degradation (AID). Like *ilc-17.1* (*syb5296*), *ilc-17.1* (*syb5297*) created using a different guide RNA also displayed a completely penetrant dauer arrest at 25°C (Fig. 1a, b; Material and Methods). Not surprisingly, CRISPR alone did not trigger dauer arrest, and a similarly large CRISPR-induced deletion in a related gene *F25D1.3, syb7367* (loss of 1475 bp of 1642 bp) did not trigger dauer arrest (Supplementary Fig. S1k). *ilc-17.1* (*syb5296*) larvae continued to arrest as dauers after they were backcrossed twice into a wild-type background (Supplementary Fig. S1k). In addition, RNAi-mediated downregulation of *ilc-17.1* in wild-type (N2) animals, but not control RNAi treatment, caused a small but significant number of larvae to arrest as SDS-resistant dauers (Supplementary Fig. S1l). ILC-17.1 signals though cytokine receptors which in *C. elegans* consists of ILCR-1/ILCR-2(26, 27). The ILCR-2 subunit is widely expressed and readily visible in the pharynx, hypodermis, intestine, neurons and germ line, as determined using a bicistronic SL2 cassette to tag the *ilcr-2* receptor at its endogenous locus with GFP (Supplementary Fig. S2a-g). Downregulation of *ilcr-2* through RNAi in wild-type larvae also caused over one-third, 39.6% ± 4%, to arrest as dauers and did not alter the dauer entry of *ilc-17.1* deleted larvae (Fig. 1e). RNAi is generally insufficient to induce dauer arrest even with known dauer pathway genes(3). Therefore, the RNAi-induced dauer entry of larvae exposed to *ilc-17.1* or *ilcr-2* RNAi was not only statistically significant, but also strongly supportive of the role of the *ilc-17.1* pathway in dauer.

*ilcr-2* RNAi downregulated *ilcr-2* expression across all somatic and germline tissues, albeit to variable extents (Supplementary Fig.S2h-m). Therefore, as additional confirmation, and to identify the tissues where ILCR-2 signaling was required to promote continuous development, we used the AID system to degrade the endogenous ILCR-2 receptor in individual tissues (Fig. 1f). We targeted the nervous system (using the *rgef-1p* promoter), pharynx (*myo-2p*), hypodermis (*col-10p)*, muscle (*unc-54p*), all somatic cells (*eft-3p*) and the germ line (*mex-5p*) individually, using animals that expressed Transport Inhibitor Response 1 (TIR1) in these tissues and endogenous *ilcr-2* fused to an AID degron. Degradation of ILCR-2 in the pharynx or in the nervous system alone caused some larvae to arrest as dauers (Fig. 1g). Degradation of ILCR-2 in the muscle, hypodermis or germ line alone had no effect (Fig. 1g). Blue Fluorescent Protein (BFP), which served as a proxy for AID, was reduced to undetectable levels upon auxin exposure in a similar percentage of larvae in all cases (Supplementary Fig. S2n), and thus did not explain the differences in dauer entry seen upon ILCR-2 degradation in the different tissues. Control larvae of the same genotype, not exposed to auxin, did not arrest as dauers. Thus, the loss of ILCR-2 receptors in the nervous system or pharynx was sufficient to induce the dauer arrest in a subset of animals.

Surprisingly, degrading ILCR-2 using TIR1 expressed under the pan-somatic *eft-3* promoter, which is expressed in somatic cells such as neurons, pharyngeal cells, intestine, hypodermis, and muscle, but not in the germ line, was less effective in inducing dauer arrest than *ilcr-2* RNAi, or degradation in the pharynx or nervous system alone: only 1.9 ± 0.8 % arrested as dauers compared to 39.6% ± 4% with *ilcr-2* RNAi, and 16.3 ± 1.1 % and 11.3 ± 1.5 % upon neuronal or pharyngeal AID (Fig. 1g). A logical hypothesis which requires future testing is that downregulation of ILCR-2 elicits complex tissue cross-talk, whereby germline-specific ILCR-2 degradation alone, while insufficient to trigger dauer arrest (e.g. *mex-5p::TIR1*), might synergize with knock-down in somatic tissue to promote dauer arrest mirroring the effects observed with *ilcr-2* RNAi. Likewise, decreased ILC-17.1 signaling in some tissues upon pan-somatic ILCR-2 degradation using the *eft-3* promoter, might antagonize the dauer-inducing effects of degradation in the pharynx or nervous system. Although more in-depth studies are required to understand the tissue-requirements, taken together, these observations support the role of ILC-17.1 signaling through the ubiquitously expressed ILCR-2 receptors in the dauer pathway. They also suggest that during normal development ILC-17.1 signaling may elicit complex inter-tissue crosstalk to promote the growth of larvae into reproducing adults or trigger their dauer arrest.

*ilc-17.1* mRNA expression is restricted to a small subset of neurons: only a pair of amphid sensory neurons (ASE) and two pairs of interneurons (AUA and RMG) expressed transgenic *ilc-17.1* reporter constructs in a previous study (26), and a few dye-filling amphid neurons expressed mCherry as a bicistronic SL2 cassette along with *ilc-17.1* mRNA from the endogenous *ilc-17.1* locus, in our studies (Fig. 1h; Supplementary Fig. S3a-g). The dauer phenotype of *ilc-17.1* deletion mutants could be rescued by re-expressing *ilc-17.1* under the control of its own promoter (Fig.1b and Supplementary Fig. S3h; see Supplementary Fig. S3i for expression levels of rescue construct in comparison to the endogenous gene). The extent of rescue was significant, although modest on OP50 (Fig.1b), but was higher on HT115 bacteria where ILC-17.1 re-expression rescued approximately 63.6% of the *ilc-17.1* deleted larvae from dauer arrest (Supplementary Fig. S3h). Re-expressing *ilc-17.1* ectopically in *ilc-17.1* deleted larvae, remote from its normal site of expression, using the muscle-specific *unc-54* promoter (Supplementary Fig. S3j-t), also rescued dauer arrest of *ilc-17.1* deleted larvae (Fig.1b and Supplementary Fig. S3h), and RNAi mediated downregulation of *ilcr-2* reduced the percentage of *ilc-17.1* larvae that were rescued by ectopically re-expressing ILC-17.1 (Supplementary Fig. S3h). The diet-dependence of rescue was not observed with ectopically expressed *ilc-17.1* (Fig.1b and Supplementary Fig. S3h). The effects of AID-induced knockdown of ILCR-2 in different tissues were also similar when larvae were fed HT115 bacteria instead of OP50 (Supplementary Fig. S3u and Fig.1g). This suggested that the role of ILC-17.1 signaling in dauer might be related to food, a prominent modulator of the dauer decision(3–9), and that the ILC-17.1 ligand, and perhaps not the *ILCR-2* receptor *per se*, could be responsive to the food signals.

To directly examine whether ILC-17.1 responded to food, we immunolocalized ILC-17.1 protein tagged at its endogenous locus using a HA-tag, in larvae that were hatched in the absence or presence of food (OP50; Fig 1h-l). While the site of *ilc-17.1* expression was not apparently altered by the presence of food, as detected by mCherry expression that remained restricted to amphid neurons (Fig. 1i, k), ILC-17.1 protein was present outside the amphid neurons, concentrated in the pharynx when larvae were exposed to food (Fig 1l), but not when larvae hatched in the absence of food (Fig.1j, quantification in Supplementary Fig. S3v; negative controls Fig. 1m,n). This suggested that ILC-17.1 protein was released from its site of expression in amphid neurons, to be either selectively translated or retained by the pharynx in response to food signals [we could not reliably detect ILC-17.1 protein in distal tissues such as the intestine or coelomocytes]. The secretion of ILC-17.1 protein was also supported by immunolocalization studies on the muscle-expressed ILC-17.1 protein which, although expressed under a well-characterized muscle-specific promoter (i.e. *unc-54*), could be constitutively detected outside muscle cells, throughout the animal (Supplementary Fig. S3m-t). This release of the ectopically expressed ILC-17.1 was not obviously affected by the absence of food (Supplementary Fig.S3q-t). These studies together showed that ILC-17.1 was released from its locus of expression in response to food, consistent with its being secreted, and that the ILC-17.1 signal, transduced through the ILCR-2 receptors in the neurons and/or pharyngeal muscle, acted cell non-autonomously to control the growth of *C. elegans*.

Sequencing total RNA (RNA-seq) extracted from bleach synchronized larvae grown for 32-36 hours at 25°C when *ilc-17.1* mutant larvae were *en route* to dauer arrest, and wild-type larvae were late L2s *en route* to developing into reproducing adults, showed that *ilc-17.1* mutant larvae expressed the molecular signatures of dauer larvae(6, 36–38) that were distinct from wild-type larvae and from *ilc-17.1* deletion mutants that re-expressed ILC-17.1 (Supplementary Figure S4a; Datasets S1-S3). This signature consisted of the downregulation of anabolic processes such as DNA replication and ribosome biogenesis (Fig. 1o), upregulation of catabolic processes and stress responses such as autophagy, fatty acid metabolism, glutathione metabolism, and xenobiotic defense pathways (Fig. 1p) and the expression of dauer-specific collagen genes that differ from the collagens expressed during continuous development(39) (Fig. 1q; Dataset S4). We evaluated whether the gene expression profile of the *ilc-17.1* deleted dauers was similar to previously described *C. elegans* dauers that accumulate in the population upon starvation or crowding by comparing our RNA-seq data with published datasets (38). Consistent with a role for ILC-17.1 in signaling nutrient availability, the expression profile of *ilc-17.1* deleted larvae was more closely correlated to dauer larvae generated by starvation(38) (Supplementary Fig. S4b), compared to ascaroside induced dauers(37) (Supplementary Fig. S4c; Datasets S5, S6). These differences were surprising and suggest that the trigger and route of dauer entry could determine their gene expression profiles (40, 41).

### ILC-17.1 loss activates CEP-1/p53 to trigger dauer entry through the activation of DAF-16/FOXO, DAF-3/SMAD-DAF-5/Ski complex and steroid hormone pathways

To understand how the loss of ILC-17.1 was triggering dauer entry we first considered the possibility that, because of its role as a neuromodulator(26), the lack of *ilc-17.1* prevented larvae from finding or ingesting food. However, this did not seem to be the case (Supplementary Fig. S5; see Extended Text 1 for details). Therefore, to answer how the loss of ILC-17.1 triggered dauer entry, we adopted a genetic approach (2–11, 29). In *C. elegans*, insulins released in the presence of food act through the sole insulin-like receptor, DAF-2, antagonize the activation of the Forkhead transcription factor DAF-16, and promote continuous growth: loss of DAF-2 signaling leads to DAF-16 activation and dauer entry(2–11, 29). Likewise, TGFβ/DAF-7 ligands antagonize the DAF-3/SMAD-DAF-5/Ski transcription factor complex to permit continuous growth: inhibition of DAF-7 activates DAF-3/SMAD-DAF-5/Ski and leads to dauer arrest(2–11, 29). Both insulins and TGF-β promote the synthesis of sterol based dafachronic acid (DA) hormones that act through the nuclear hormone receptor, DAF-12, to support adult development(2–11, 29). Loss of DA pathway components also triggers dauer arrest, and providing exogenous DA to, or decreasing DAF-12 activity in, animals deficient in insulin or TGFβ signaling, can bypass their dauer arrest(3, 42). Finally, a mutation in a transmembrane guanylyl cyclase *daf-11* that causes chemosensory defects also triggers dauer arrest(3, 43). *ilc-17.1* deletion mutants lacked obvious chemosensory defects (Supplementary Fig. S5; Extended Text 1) and continued to arrest as dauers on 5mM 8-Bromo-cGMP, which modestly rescues *daf-11*-associated dauer formation(3, 43) (Supplementary Fig. 6a). Therefore, to identify signal transduction pathways through which the loss of ILC-17.1 signaling controlled the dauer decision, we examined its genetic interactions with *daf-16, daf-5*, and DA signaling.

A *daf-16* loss of function mutation*, mu86*(44), completely suppressed dauer formation in animals lacking ILC-17.1, indicating that activation of DAF-16 was one mechanism responsible for their dauer arrest (Fig. 2a). The activation of DAF-16 in animals lacking ILC-17.1 could be quantified by an increase in the proportion of larvae with DAF-16::GFP(45) positive intestinal nuclei even at 20°C (Fig. 2b-d), and the upregulation of DAF-16 target genes(46, 47) both at 20°C (Supplementary Fig. S6b) and 25°C (Fig.2e). DAF-16 activation could be suppressed by re-expressing ILC-17.1 in *ilc-17.1* deletion mutants under its endogenous promoter, or ectopically, under the muscle-specific promoter (Fig.2e). Surprisingly, a loss of function mutation in *daf-5*, *e1386*(48) also completely suppressed the dauer arrest of *ilc-17.1* mutant larvae (Fig. 2f), and exposing *ilc-17.1* deficient larvae to 50 nM exogenous Δ7-dafachronic acid (42), partially but significantly suppressed their dauer arrest (Fig 2g). The activation of DAF-16/FOXO and DAF-3/SMAD-DAF-5/Ski by the loss of ILC-17.1 was not simply because *ilc-17.1* deleted larvae were deficient in the expression of insulin signaling and TGFβ/ DAF-7 signaling pathway components, as determined by expression levels of the signaling pathway components and transcriptional reporters [Supplementary Fig S7; Datasets S7, S8, see Extended Text 2 for details]. The dauer arrest of *daf-2* (*e1370*)(49), could not be rescued by ILC-17.1 overexpression, suggesting that ILC-17.1 did not act genetically downstream of *daf-2* (Supplementary Fig. S8a). Thus, it appeared that ILC-17.1 acted upstream of both DAF-16/FOXO and DAF-3/SMAD-DAF-5/Ski (and likely, DAF-9/DAF-12), but not downstream of DAF-2, to modulate dauer entry.

**Figure 2:**
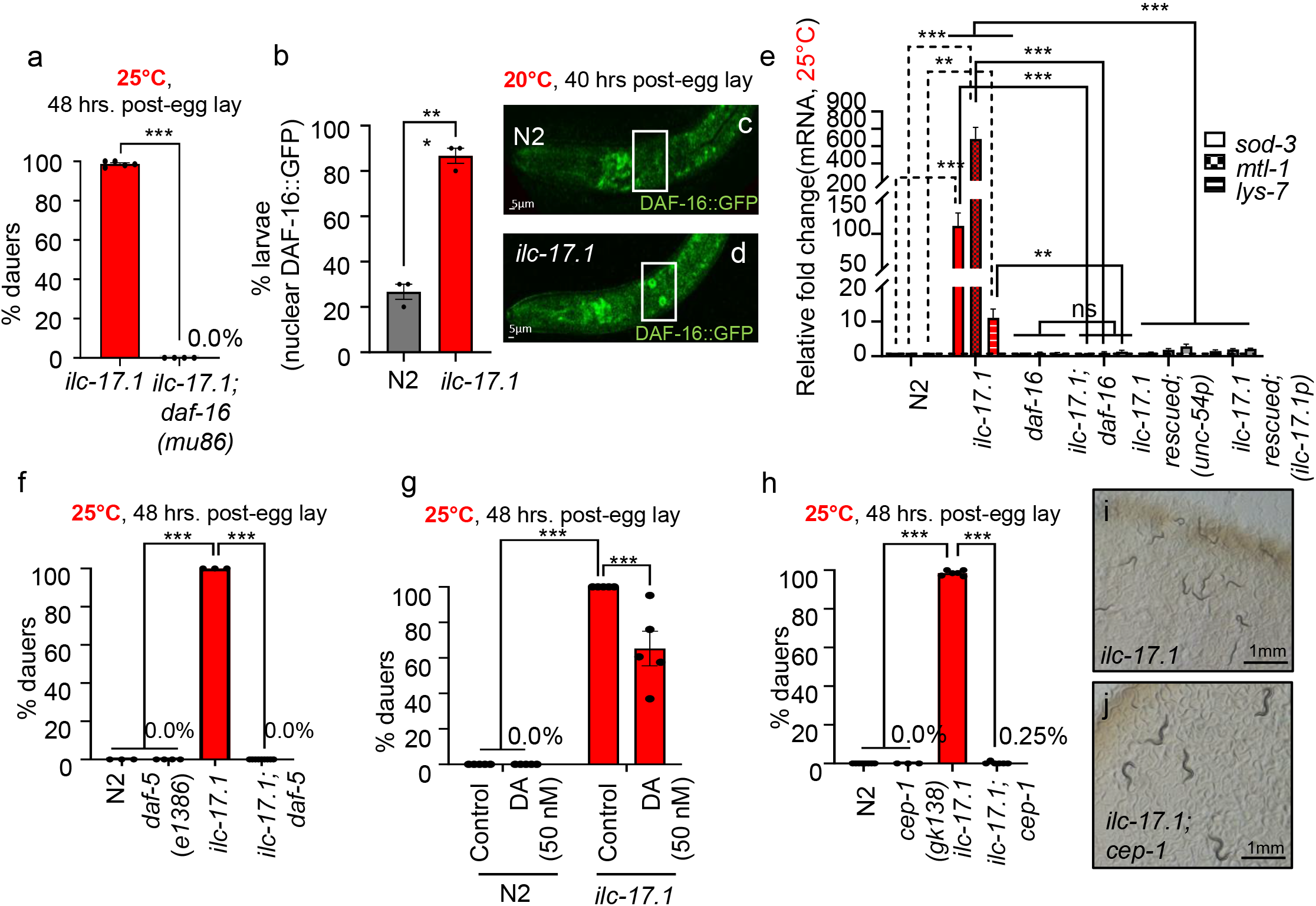
Reduced ILC-17.1 signaling acts through DAF-16/FOXO, DAF-3/SMAD-DAF-5/Ski complex, steroid hormone signaling and CEP-1/p53 to control dauer arrest. **(a),** Percent dauers. X-axis: genotype. (*n*=4-8 experiments, Chi-squared = 497.65, df = 1, p-value < 2.2e-16). **(b),** Percent 36 hrs larvae, grown at 20°C showing nuclear DAF-16::GFP. *n*=3 experiments with 10-15 larvae each. Each data point represents average no. of larvae with nuclear DAF-16::GFP/experiment, (unpaired t-test). Note: experiments were conducted at 20°C, since DAF-16::GFP localized constitutively in intestinal nuclei even in wild-type larvae at 25°C. **(c, d),** representative micrographs of DAF-16::GFP (boxed) in wild type and *ilc-17.1* deletion mutant larvae. Scale bar, 5µm. **(e),** Average *sod-3, mtl-1, and lys-7* mRNA levels in larvae 32-36 hr. post-hatching at 25°C. mRNA levels are relative to *pmp-3* and normalized to wild type (N2) values. *n*=4-10 experiments, (unpaired t-test). (**f),** Percent dauers. X-axis: genotype. (*n*=3-5 experiments, Chi-squared = 787.3, df = 3, p-value < 2.2e-16). (**g),** Percent dauers in *ilc-17.1* mutants on control (H_2_O) and exogenous 50nM Δ7-dafachronic acid (DA), (*n*=3-5 experiments, Chi-squared = 784.32, df = 3, p-value < 2.2e-16). (**h),** Percent dauers, X-axis: genotype (*n*=3-5 experiments, Chi-squared = 1029.2, df = 3, p-value < 2.2e-16). **(i, j),** Representative micrographs of **i,** *ilc-17.1* deleted larvae arrested as dauers, and j, *ilc-17.1;cep-1* that grew to L4s, 48 hr. post-hatching at 25°C. Scale bar, 1 mm. Bars show the mean ± S.E.M. Individual data points in bar graphs in **a, f-h,** represent the average % dauers/experiment. ***p < 0.001, **p < 0.01, *p < 0.05 and ns, not significant

This was curious, since DAF-16/FOXO and DAF-3/SMAD-DAF-5/Ski are thought to act independently, and in parallel to modulate the *C. elegans* dauer decision(3, 5). Therefore, to identify mechanism(s) that could activate both pathways in *ilc-17.1* deleted larvae, we conducted RNAi knockdown of candidate genes known to interact with these dauer effectors (50–57), and asked which, if any, also rescued the dauer entry of *ilc-17.1* larvae (Supplementary Fig. S8b). We considered only the positive hits, due to the variable penetrance of RNAi. Amongst the positive candidates, we found most interesting, *cep-1*, the *C. elegans* ortholog of p53(58, 59), since p53-family of proteins restrain cell cycle progression and alter metabolism, both cardinal features of dauer entry. Indeed, a deletion in *cep-1*, *gk138*, rescued the dauer arrest of *ilc-17.1* deletion mutants and nearly all (98%) of *cep-1;ilc-17.1* double mutant larvae grew into reproductive adults (Fig 2h-j), suggesting that CEP-1/p53 was also responsible for the dauer phenotype of *ilc-17.1* mutants, and was activated.

In mammalian cells, p53 activation occurs through an increase in p53 protein levels due to its escape from constitutive degradation by the E3 ubiquitin ligase MDM2(18, 20, 60). Although the *C. elegans* genome lacks MDM2 orthologs, CEP-1/p53 activation through translational and posttranslational modifications has been reported (61, 62). We therefore used Western analysis of FLAG-tagged endogenous *C.* elegans CEP-1/p53 to test whether CEP-1/p53 was activated in *ilc-17.1* mutant larvae, as implied by the rescue. This was the case: in *ilc-17.1* deleted larvae, CEP-1::FLAG levels increased by 2-fold, levels similar to that seen upon gamma irradiation, which also activates CEP-1/p53, and was used as a positive control (Fig 3a) (63, 64). Accordingly, overexpressing CEP-1 alone, even to a modest 1.5-fold level over wild type (see Supplementary Fig. S9a for *cep-1 o/e* mRNA levels), was sufficient to arrest growth and promote an almost completely penetrant dauer phenotype (Fig. 3b-d). CEP-1 overexpression was achieved by crossing out the *cep-1 lg12501* deletion mutation from a widely used transgenic strain expressing functional, fluorescently tagged, CEP-1, CEP-1::GFP, under its own promoter (61, 65, 66). In the *cep-1*(*lg12501*) strain, CEP-1::GFP is the only source of CEP-1/p53, rescues *cep-1* (*lg12501*) apoptosis defects and is expressed at half the levels of wild type (Supplementary Fig. S9a; *cep-1* mRNA levels). CEP-1::GFP did not induce dauer arrest in the *cep-1, lg12501* deletion background (Fig. 3b), indicating that *cep-1* expression levels, and not the GFP tag, was responsible for the dauer arrest of CEP-1::GFP overexpressing larvae. The overexpression of CEP-1/p53 phenocopied the *ilc-17.1* deletion, not only causing all larvae to arrest as dauers at 25°C, but also prompting transient dauer entry at 20°C, as assessed by SDS-resistance (Supplementary Fig. S9b). RNA-seq analysis showed that genes differentially expressed in CEP-1/p53 overexpressing larvae *en route* to their dauer arrest, significantly overlapped with those expressed in larvae lacking ILC-17.1 (Fig. 3e, padj.< 0.05; Datasets S9, S10). CEP-1/p53 overexpressing larvae also upregulated dauer-specific collagens (Supplementary Fig. S9c; Dataset S4), downregulated anabolic pathways (Supplementary Fig. S9d), and upregulated catabolic pathways (Supplementary Fig. S9e). In addition, between 15 hours to 32 hours post-hatching at 25°C, CEP-1/p53 overexpressing larvae and *ilc-17.1* deletion mutants showed a remarkably constant gene expression profile, and only 340 and 34 genes respectively changed expression (p<0.05), compared to the > 10,000 genes that were either upregulated or downregulated in wild-type larvae and *cep-1;ilc-17.1* double mutant larvae that *en route* to becoming reproductive adults (PCA analysis; Supplementary Fig. S9f; Dataset S11). The upregulation of known CEP-1/p53 targets (63, 64, 67, 68), the BH3-only proteins *egl-1* and *ced-13*, in the *ilc-17.1* deletion mutants, in a *cep-1*-dependent manner, at both 20°C and 25°C, further confirmed CEP-1/p53 activation in the absence of ILC-17.1 (Fig 3f and Supplementary Fig. S10a). The upregulation of CEP-1/p53-targets in *ilc-17.1* deletion mutants could be suppressed by re-expressing ILC-17.1 under its endogenous-, or muscle-specific promoters (Fig 3f; Supplementary Fig. S10a). In addition, <u>Ch</u>romatin Immuno<u>p</u>recipitation followed by quantitative PCR (ChIP-qPCR) using FLAG tagged *cep-1* animals showed that the loss of *ilc-17.1* increased CEP-1/p53 binding at promoter regions of *egl-1* and *ced-13*, and the increased occupancy was abrogated upon re-expressing ILC-17.1 (Fig. 3g, h).

**Figure 3:**
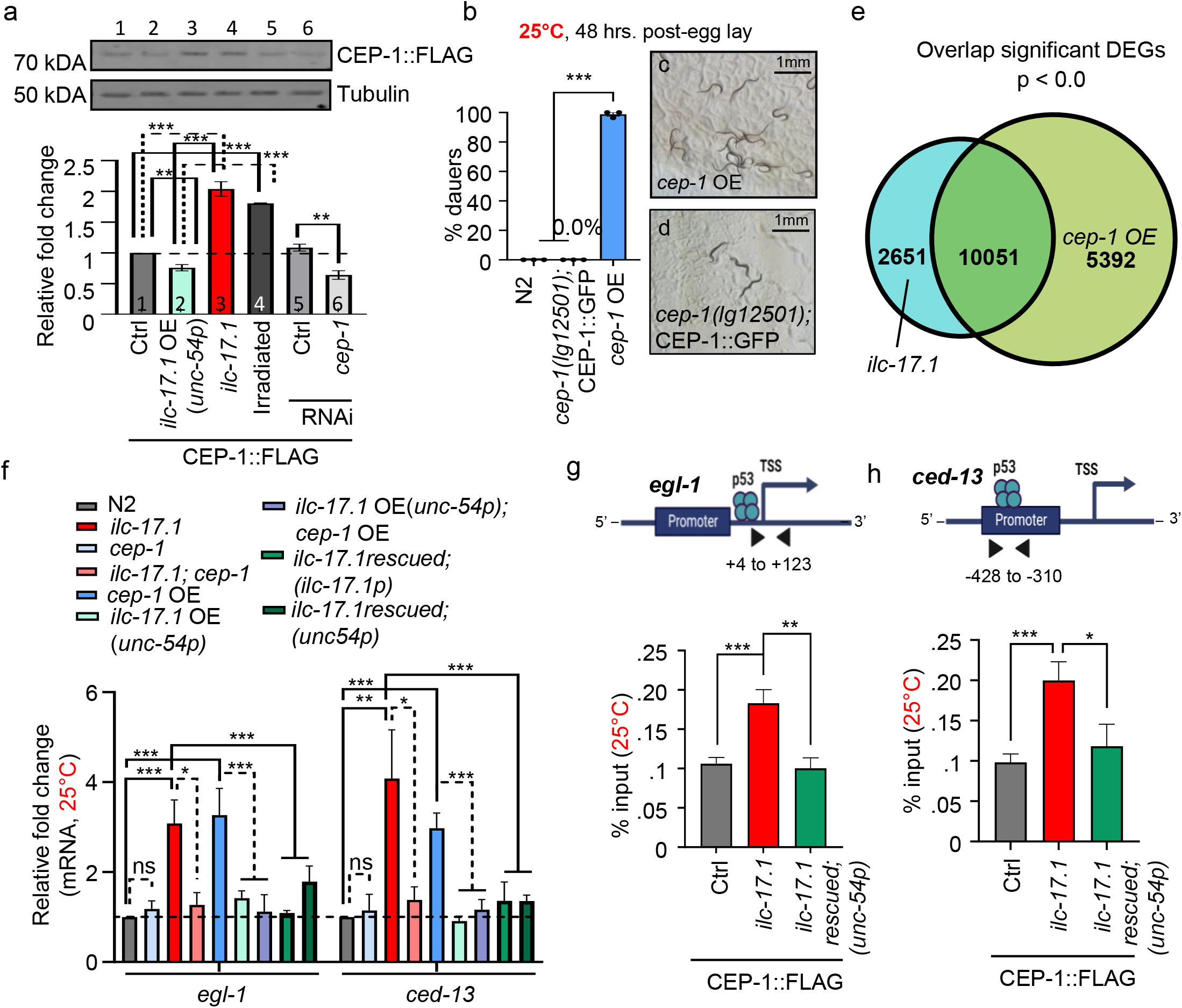
CEP-1/p53 is activated by reduced ILC-17.1 signaling. **(a), Top**: Representative Western blot showing CEP-1::FLAG and tubulin. Bands on western correspond to numbers in data bars. *cep-1* RNAi used for specificity. Samples constitute 32-36 hr. larvae grown at 25°C, except for positive control, irradiated sample (#4; adults). **Bottom**: Average CEP-1 levels quantified relative to tubulin and normalized to control (Ctrl; CEP-1::FLAG in wild-type background). n=4 experiments. (unpaired t-test). (**b),** Percent dauers. X-axis: genotype. (n=3-4 experiments, Chi-squared = 569.35, df = 2, p-value < 2.2e-16). mRNA expression levels in *cep-1* OE and *cep-1 (lg12501)*; CEP-1::GFP were ∼1.5X and ∼0.5X wild-type levels, respectively, see **Supplementary Fig. S9a**. **(c, d),** Representative micrographs 48 hrs. post-hatching at 25°C, of **c,** CEP-1/p53 overexpressing larvae arrested as dauers and **d,** *cep-1 (lg12501)*; CEP-1::GFP L4s. Scale bar, 1 mm. **(e),** Venn diagram depicting overlap between differentially expressed genes (p<0.05) in *ilc-17.1* deletion mutants and CEP-1/p53 overexpressing larvae, 32-36 hr. post-hatching at 25°C. p<0.0; hypergeometric test. **(f)**, Average *egl-1* and *ced-13* mRNA levels in 32-36 hr. larvae grown at 25°C. n=6-7 experiments. mRNA levels relative to *pmp-3* and normalized to wild-type (N2) values. (unpaired t-test). **(g, h),** CEP-1 occupancy (percent input; CEP-1::FLAG was immunoprecipitated with anti-FLAG antibody) at the promoter proximal regions of **g,** *egl-1* and **h,** *ced-13* in 32-36 hr. larvae grown at 25°C, (**Top,** schematic). X-axis: genotype. n=4 experiments. (unpaired t-test). (**a, g, h**), endogenous *cep-1* was FLAG tagged at its C-terminus using CRISPR/Cas9, crossed into backgrounds noted and probed with anti-FLAG antibody. Bars show the mean ± S.E.M. Individual points in bar graphs in **b** represent the % dauers/experiment. ***p < 0.001, **p < 0.01, *p < 0.05 and ns, not significant.

Like in the ILC-17.1 deficient animals, the dauer arrest of larvae overexpressing CEP-1 was dependent on DAF-16, DAF-5 and the steroid hormone pathway: downregulating *daf-16* expression by RNAi (Fig. 4a), crossing CEP-1/p53 overexpressing larvae into *daf-5*, *e1386*, or treatment with 50nM exogenous Δ7-dafachronic acid, all suppressed their dauer arrest (Fig. 4b). In accordance, DAF-16 target genes were also upregulated in larvae overexpressing CEP-1/p53 at both 25°C (Fig 4c) and 20°C (Supplementary Fig. S10b). Differentially expressed genes in CEP-1/53 overexpressing larvae overlapped with published gene expression profiles of *daf-2 (e1370)* and active DAF-16 (Supplementary Fig. S10 c,d; Datasets S12, S13). In addition, and most convincing, the upregulation of *daf-16* target genes in *ilc-71.1* deletion mutant larvae was *cep-1* dependent at both 25°C and 20°C (Fig 4c; Supplementary Fig. S10b). CEP-1/p53 activation-induced dauer entry appeared to be somewhat specific to reduced ILC-17.1 signaling, as the dauer arrest of *daf-2*(*e1370*) larvae that occurred due to reduced insulin signaling did not depend on *cep-1* (Supplementary Fig. S10e). Moreover, in agreement with previous reports(69, 70), *cep-1* was not required for dauer induction at high temperatures as *cep-1* (*gk138*) larvae could enter dauer at 27°C just like wild-type animals (Supplementary Fig. S10f). Taken together, these results showed that ILC-17.1 normally acted to represses CEP-1/p53 transcriptional activity. Upon loss of ILC-17.1 signaling, CEP-1/p53 was activated, in turn activated DAF-16/FOXO (and perhaps DAF-3/SMAD-DAF-5/Ski and DAF-12) to trigger the dauer arrest of *ilc-17.1* deletion mutants. However, the ILC-17.1/CEP-1 dauer axis appeared independent of the insulin-like receptor, *daf-2*, *per se*.

**Figure 4.**
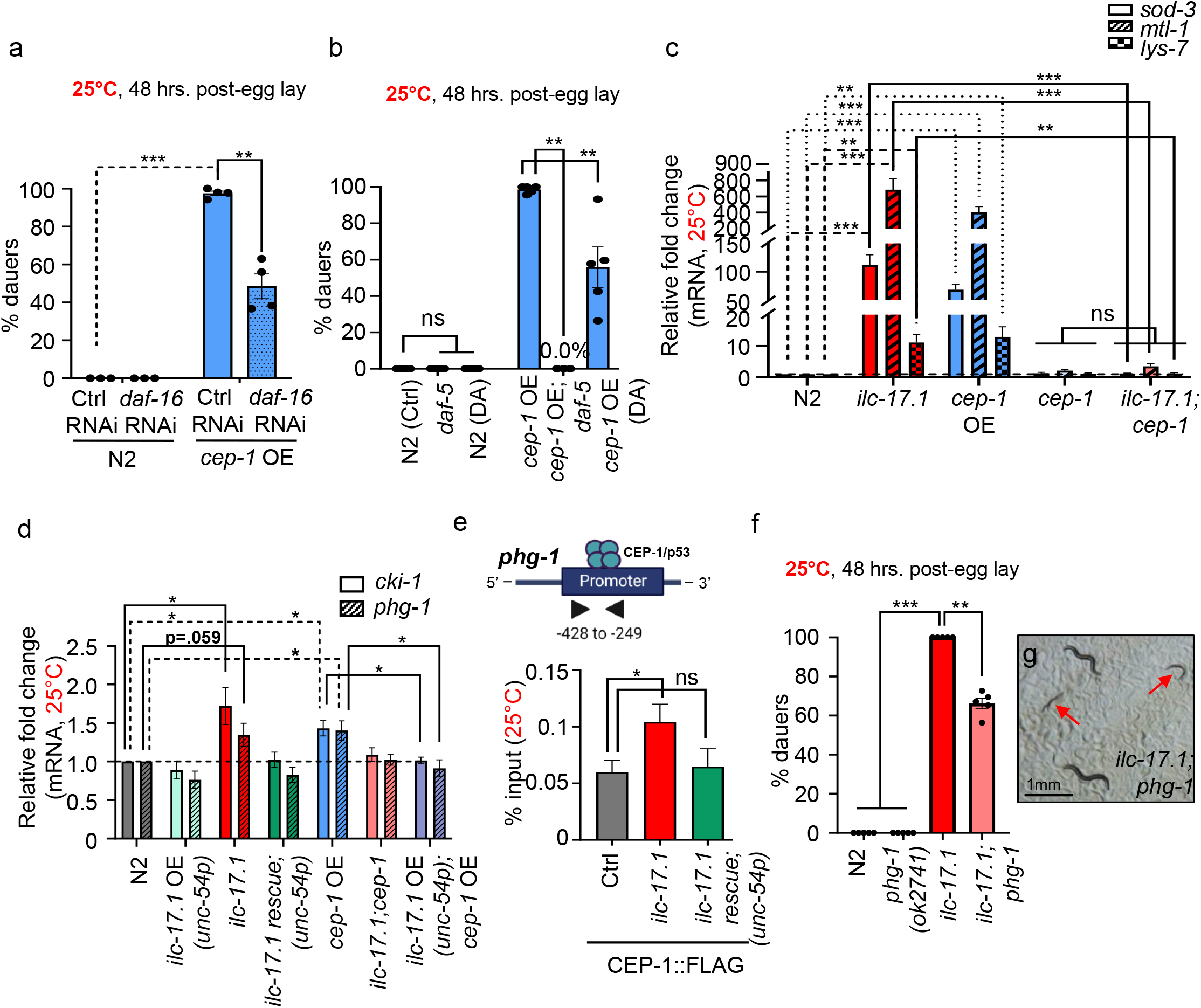
CEP-1/p53 acts through DAF-16/FOXO, DAF-3/SMAD-DAF-5/Ski complex and cell cycle inhibitors to control dauer entry. **(a)**, Percent dauers. X-axis: genotype and RNAi treatment: Ctrl, L4440 or *daf-16*. (n=3-4 experiments, Chi-squared = 494.22, df = 3, p-value < 2.2e-16). **(b),** Percent dauers. X-axis: genotype and treatment: water (control) and 50nM Δ7-dafachronic acid (DA). (n=3-4 experiments, Chi-squared = 982.66, df = 5, p-value < 2.2e-16). **(c),** Average *sod-3, mtl-1, and lys-7* mRNA levels in larvae 32-36 hr. post-hatching at 25°C. n=4-10 experiments. (unpaired t-test). **(d)**, Average *cki-1* and *phg-1* mRNA levels in 32-36 hr. larvae grown at 25°C. (**c, d**) mRNA levels were determined relative to *pmp-3* and normalized to wild-type (N2) values. n=4-6 experiments. *p < 0.05. (unpaired t-test). **(e),** CEP-1 occupancy (percent input) in 32-36 hr. larvae grown at 25°C, at the promoter proximal region of *phg-1* (**Top,** schematic). X-axis: genotype. n=4 experiments. (unpaired t-test). **(f),** Percent dauers. X-axis: genotype. (n=6 experiments, Chi-squared = 826.95, df = 3, p-value< 2.2e-16). **(g),** Representative micrograph 48 hrs. post-hatching at 25°C of *ilc-17.1;phg-1* double mutants, showing dauers (arrows) and L4s. Scale bar, 1 mm**. (a**, **b**, **f),** Bar graphs: bars show the mean ± S.E.M, and individual points represent the % dauers/experiment. ***p < 0.001, **p < 0.01, *p < 0.05 and ns, not significant.

### CEP-1/p53 directly or indirectly controls cell cycle progression

A conserved function of the p53-family of proteins is to restrain cell cycle progression. In *C. elegans*, most cell divisions are completed during embryogenesis, but a subset of somatic and germ line cells divide and differentiate post-embryonically, and these cells arrest during dauer entry(71–76) (77, 78). (79–86). We therefore tested whether CEP-1/p53 activation upon loss of ILC-17.1 was participating in the arrest of cell cycle progression during postembryonic development. Indeed, *ilc-17.1* deleted larvae upregulated mRNA levels of cell cycle inhibitors *cki-1,* one of the two p21 homologs, and *phg-1,* the *C. elegans* homologue of Growth arrest-specific 1 (Gas1) (Fig 4d). These are known to regulated by CEP-1/p53 (87). *phg-1,* but not *cki-1,* was a direct target of CEP-1/p53, in *ilc-17.1* deletion mutants, which displayed increased CEP-1/p53 occupancy at the *phg-1* promoter (Fig. 4e; Supplementary Fig. S11a). Again, as with the other CEP-1/p53 targets, the increased expression of *cki-1* and *phg-1,* and the increased occupancy at the *phg-1* promoter was restored to wild-type levels upon re-expressing ILC-17.1 (Fig. 4d, e). Surprisingly, downregulation of *phg-1* and *cki-1* by RNAi modestly, but significantly, rescued dauer arrest in *ilc-17.1* deletion mutants and CEP-1/p53 overexpressing larvae and allowed a small number of larvae to develop into adults (Supplementary Fig. S11b). Crossing *ilc-17.1* deletion mutants into *phg-1* deletion mutants, *ok2741* also resulted in a rescue and ∼40% escaped dauer arrest and grew into reproductive adults (Fig. 4f, g). In addition, a proportion of *ilc-17.1* deletion mutant and CEP-1/p53 overexpressing larvae that appeared morphologically dauer, displayed modest but significant non-dauer physiological traits on *phg-1* and *cki-1* RNAi such as increased rates of sporadic pumping (Supplementary Fig. S11c; wild-type L2 were used as controls), suggesting there may also occur a wider, albeit incomplete, dauer rescue in some tissues. The dauer arrest of *daf-2* mutant larvae which did not depend on *cep-1,* was also not rescued upon downregulating *phg-1* or *cki-1*, nor were the pumping rates of *daf-2* dauers altered (Supplementary Fig. S11 b,c). Although we cannot distinguish at this time, whether the downregulation of the cell cycle inhibitors allowed larvae to exit dauer, or suppressed dauer entry and complete arrest, these data showed that loss of ILC-17.1 signaling or overexpressing CEP-1/p53 upregulated CEP-1/p53-dependent cell cycle inhibitors, and this increase appeared to be one mechanism responsible, at least in part, for the dauer arrest.

### ILC-17.1 links glucose metabolism to CEP-1/p53 activity

Mammalian p53 plays a central roles in energy metabolism and, in turn, can be activated by energy deprivation or limited glucose, through signaling by AMP-activated protein kinase (AMPK) or Mammalian target of rapamycin (mTOR) pathways (88, 89). Therefore, the responsiveness of ILC-17.1 to food signals suggested the intriguing possibility that the loss of ILC-17.1 signaling activated CEP-1/p53 because of energy deprivation, and this could link nutrient availability to metabolism and cell cycle progression. AMPK did not appear to be activated in *ilc-17.1* deleted animals (Supplementary Fig. S12a), nor was mTOR activity obviously reduced (Supplementary Fig. S12b). However, ILC-17.1 signaling did appear to link glucose utilization to developmental progression, since downregulating *fgt-1,* the main Glucose transporter (GLUT) responsible for glucose absorption in *C. elegans*(90), decreased the extent of dauer-rescue conferred by re-expressing ILC-17.1 in *ilc-17.1* mutant larvae (Fig. 5a), and increased the numbers of *ilc-17.1* deletion mutant larvae that arrested as dauers at 20°C (Fig. 5b). Conversely, extra glucose(91) suppressed the dauer arrest of approximately 30% *ilc-17.1* deficient larvae (Fig. 5c; Supplementary Fig. S12c), suggesting that the normal amounts of glucose available in the diets of *C. elegans* larvae was insufficient for *ilc-17.1* larvae to develop into adults. Consistently, decreasing glucose import through *fgt-1* RNAi once again, lowered the percentage of larvae that were rescued by the extra glucose (Supplementary Fig. S12d), and no rescue was observed on the non-hydrolysable glucose analog 2-Deoxy-d-glucose (2-DOG; Supplementary Fig. S12e) which cannot undergo glycolysis and does not enter the metabolic pathway. The same concentrations of glucose could not rescue *daf-*2 larvae that arrested as dauers (Fig 5c). We ruled out non-specific effects of glucose rescue by testing the role of increased ROS or other supplements (Supplementary Fig S12e-f; see Supporting Information; Extended Text 3 for details). Downregulating *fgt-1* in wild-type larvae, while not sufficient to trigger dauer arrest, modestly increased the expression of CEP-1/p53 target *egl-1* (but not *ced-13*; Supplementary Fig. S12g). Thus, these data supported the possibility that the activation of CEP-1/p53 upon the loss of ILC-17.1 could, at least in part, occur due an impairment in the *ilc-17.1* deleted larvae’s ability to utilize glucose.

**Figure 5.**
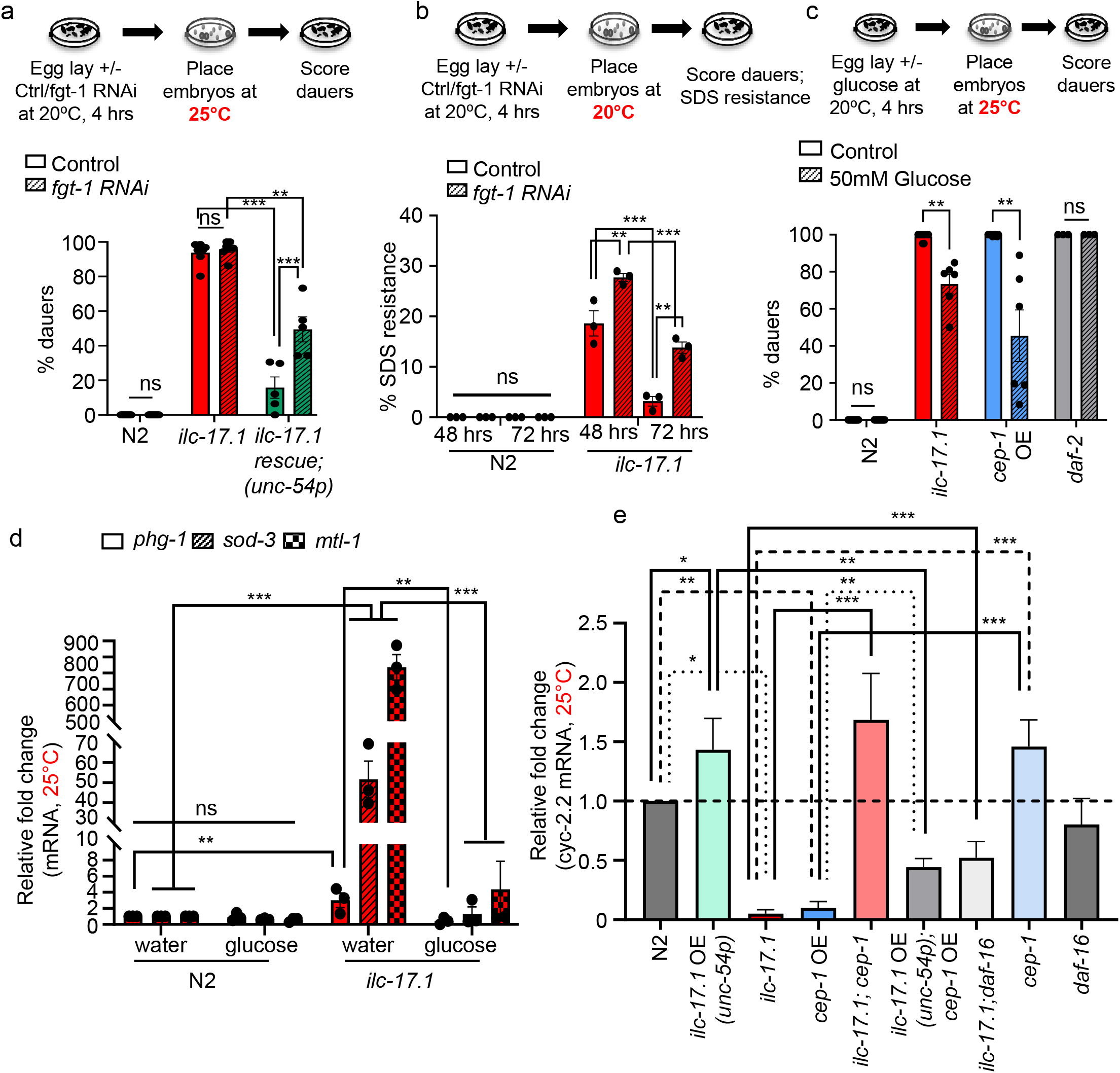
Impaired glucose utilization in ILC-17.1 deficient larvae activates CEP-1/p53 and alters the expression of OXPHOS genes. **(a), Top**: Schematic of experimental design. **Right**: Percent dauers. X-axis: genotype. Larve treated with control (L4440) and *fgt-1* RNAi. (n*=* 5 experiments, Chi-squared = 1817.1, df = 5, p-value < 2.2e-16). **(b), Bottom**: Schematic of experimental design. **Right**: Percent SDS (1%)-resistant dauers at 48 and 72 hrs. post-hatching and grown at optimal conditions of 20°C. X-axis: genotype. Larve treated with control (L4440) and *fgt-1* RNAi. (n*=* 3 experiments, Chi-squared = 206.82, df = 7, p-value < 2.2e-16). **(c), Top**: Schematic of experimental design. **Bottom**: Percent dauers. X-axis: genotype and treatment: Control (H_2_O) and 50mM glucose. (n*=* 3-6 experiments, Chi-squared = 2882.3, df = 5, p-value < 2.2e-16). **(d),** Average *phg-1, sod-3 and mtl-1* mRNA levels in 32-36 hr. larvae that arrest as dauers (on H_2_O, control) and larvae that bypass dauer (50mM glucose) at 25°C. mRNA levels are relative to *pmp-3* and normalized to wild-type (N2) values. n=3 experiments. (unpaired t-test). **(e),** Average *cyc-2.2* mRNA levels in 32-36 hr. larvae. mRNA levels are relative to *pmp-3* and normalized to the wild-type (N2) values. n=4-6 experiments.. (unpaired t-test). Bars show the mean ± S.E.M. Individual points in the bar graphs in **a, b, c** represent the % dauers/experiment, ***p < 0.001, **p < 0.01, *p < 0.05, ns, non-significant.

Glucose supplementation also rescued dauer arrest of CEP-1/p53 overexpressing larvae (Fig 5c; Supplementary Fig. S10c), arguing that decreased glucose intake in the absence of ILC-17.1 was not a consequence of CEP-1/p53 activation, but instead acted logically upstream of CEP-1/p53. In agreement, extra glucose suppressed the increased expression of *phg-1* the direct target of CEP-1/p53, and decreased DAF-16/FOXO activation (Fig.5d). Surprisingly, not only did CEP-1/p53 appear to be activated by the inability of *ilc-17.1* to utilize glucose normally, but CEP-1/p53 activity, in turn decreased the expression levels of key enzymes involved in glucose metabolism(92): phosphofructokinase-1.2 (*pfk-1.2*), the rate limiting enzyme in glycolysis, and cytochrome *c* (*cyc-2.2*), the complex IV subunit of the mitochondrial electron transport chain responsible. This was visible in the RNA-seq data (Supplementary Figure S13 a-f; Dataset S14) and confirmed by qRT-PCR (Supplementary Figure S13g, Fig. 5e). Moreover, although the downregulation of *pfk-1.2* or *cyc-2.2* alone was not sufficient to induce dauer in wild-type animals, RNAi knockdown of *cyc-2.2* in *ilc-17.1* deleted larvae increased the percentage of dauers under optimal conditions of 20°C, suggesting that CEP-1/p53-dependent decrease of *cyc-2.2* could feedback onto *ilc-17.1* loss to increase dauer propensity (Supplementary Figure S13h). These data, together, suggested that the loss of ILC-17.1 impaired glucose utilization and this activated CEP-1/p53, which, then further limited glucose utilization by downregulating genes important for glycolysis and OXPHOS. Thus, ILC-17 secretion from amphid neurons that occurred in response to food availability appeared to link cell fate decisions with energy metabolism through CEP-1/p53.

### ILC-17.1 suppresses CEP-1/p53 in *C. elegans* and human epithelial cells

Our data pointed towards a model whereby ILC-17.1 signaling suppressed CEP-1/p53. Therefore, we tested this directly. Indeed, overexpressing ILC-17.1 in animals also overexpressing CEP-1/p53 inhibited their dauer arrest (Fig. 6a). In addition, since the most-accepted and canonical function of *C.elegans* CEP-1/p53 is in triggering apoptosis in the adult germ line in response to DNA-damage, we also examined whether ILC-17.1 repressed CEP-1/p53’s function in apoptosis. As has been previously shown(67), the loss of CEP-1/p53 in *cep-1 (gk138)* adults has negligible effects on physiological apoptosis, but suppresses the increased apoptosis that occurs in response to gamma irradiation (Fig. 6b). CEP-1/p53 overexpression, surprisingly, caused a significant increase in physiological apoptosis, and a marked increase in irradiation-induced apoptosis, both of which could be suppressed by also overexpressing ILC-17.1 (Fig 6b). Overexpressing *ilc-17.1* in a wild-type background also decreased physiological and irradiation-induced apoptosis, but this effect was variable and did not reach significance (average nos. of apoptotic cells in non-irradiated N2 v ILC-17.1 overexpressing animals were 3.56 ± 1.3 and 2.53 ± 1.1 cells, and upon irradiation, increased to 7.1 ± 1.8 in N2, and 5.72 ± 2.2 in IL-17 overexpressing animals). For reasons that remain to be understood, ilc*-17.1* deletion did not alter apoptosis in the adult germ line, perhaps reflecting a different function post-development. IL-17 could also modestly repress p53, autonomously, in mammalian cells (Supplementary Fig S14a-b; see Extended Text 4 for details). Together, these data suggest that the inhibition of CEP-1/p53’s activity could be a conserved function of IL-17s.

**Figure 6:**
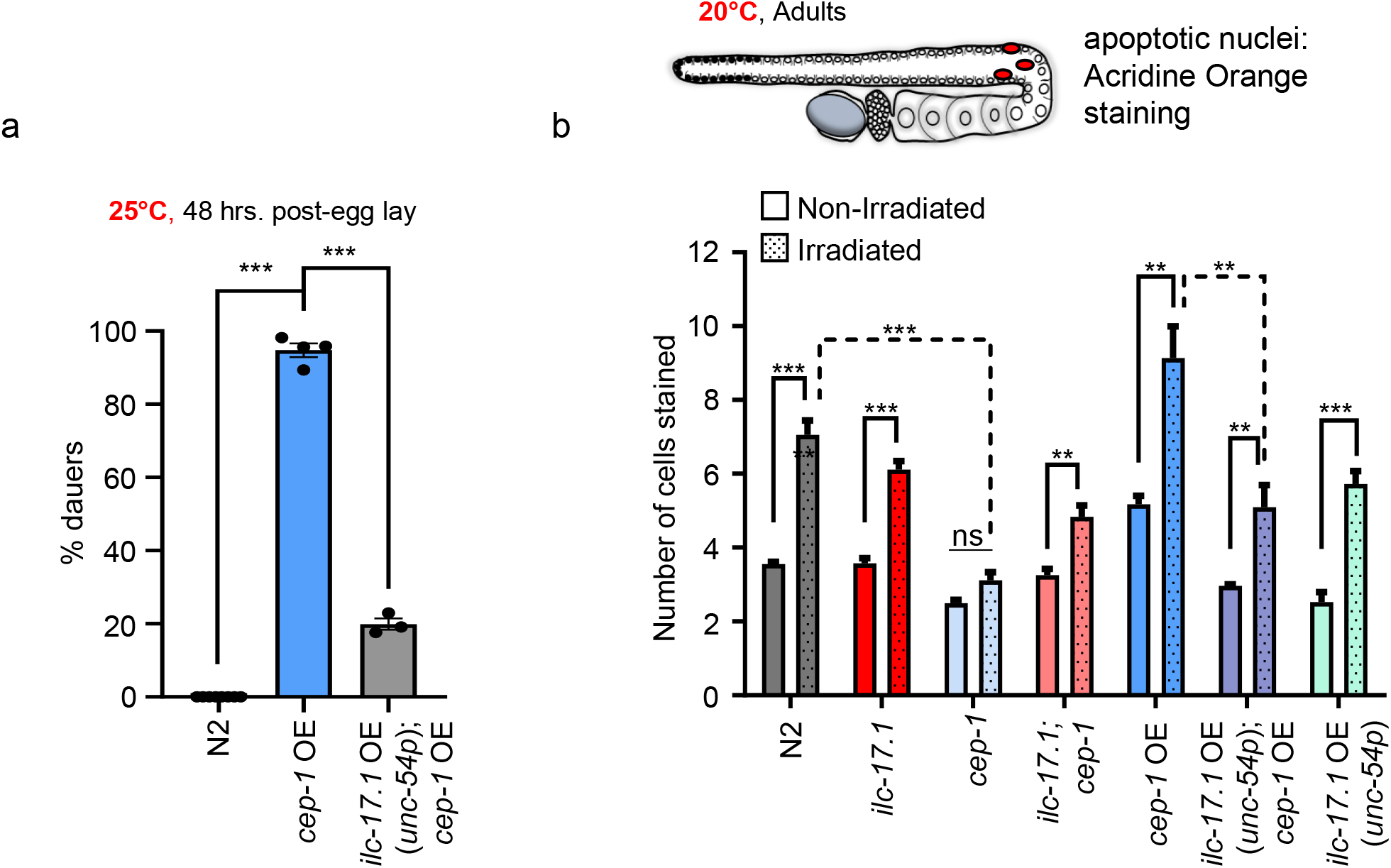
ILC-17.1 suppresses CEP-1/p53. **(a),** Percent dauers. X-axis: genotype. (n=4 experiments, Chi-squared = 1245.4, df = 2, p-value < 2.2e-16). **(b), Top:** Schematic of germline-specific apoptosis (apoptotic cells in red) scored with Acridine Orange. **Bottom:** Average numbers of apoptotic cells in day-two adult animals under control, non-irradiated conditions, and upon irradiation with 75 Gy (day-one adults were irradiated). n*=* 3 experiments, and 12 gonad arms/experiment. (analysis of variance (ANOVA) with Tukey’s correction, df=13). Note: p values between N2 vs *ilc-17.1* OE (*unc-54p*), non-irradiated and irradiated are p=0.0192 and p=0.0623, if compared by themselves (unpaired t-test), but do not rise to significance when corrected for the multiple comparisons. Data in all graphs show mean ± S.E.M ***p < 0.001, **p < 0.01.

## Discussion

Here we show roles for the interleukin 17 (IL-17) family of cytokines and p53 in the development of an invertebrate species, *C. elegans*. Our data support a model whereby ILC-17.1 signaling of food availability by amphid neurons ensures the continuous growth of larvae into reproductive adults under favorable conditions through suppression of CEP-1/p53. IL-17s play broad-ranging immunometabolic roles in mammals and are one of the most conserved proinflammatory cytokines among animal phyla. Thus, their ancestral role in signaling nutrient availability as in *C. elegans* may be broadly conserved. Our data also reveal that reduced ILC-17.1 signaling, expected to occur when food is scarce, activates CEP-1/p53, to trigger dauer arrest through the well characterized dauer activating pathways, DAF-16/FOXO, DAF-3/SMAD-DAF-5/Ski and also, perhaps steroid hormone signaling. In its role in promoting dauer arrest in *C. elegans*, CEP-1/p53 inhibits cell cycle progression, and modulates metabolic flux by directly or indirectly controlling the levels of phosphofructokinase (*pfk-*1) and cytochrome C (*cyc-2.2*). Notably, these are also mechanisms by which p53 restrains tumorigenesis. In mammals too, uncontrolled p53 activity is associated with a developmental syndrome through its effects on cell cycle progression, apoptosis and the migration of neural crest cells(25). We therefore cautiously propose that the ancestral roles of the p53 gene-family in modulating development could have the driving force in their evolution.

While, there remain several puzzling aspects associated with this study, our data indicate that there may exist a previously undiscovered *C. elegans* dauer pathway. The dauers formed by *ilc-17.*1 deletion, or the modest 1.5X increase in CEP-1/p53 expression, display all the cardinal morphological and physiological features of dauers, including SDS-resistance, decreased pumping rates, presence of alae, changes in collagens, upregulation of catabolic and downregulation of anabolic genes, and the ability to remain arrested for days to weeks as dauers. Nevertheless, they also differ in some respects from the well-studied dauers formed through decreased insulin signaling (*daf-2*), and other pathways, in that, unlike previously studied dauers, a modest, but significant percentage of *ilc-17.1* dauers larvae bypass or are rescued from dauer upon downregulating cell cycle inhibitors such as *phg-1*, or when fed extra glucose. This suggests that there may exist different ‘types’ of dauer larvae. The discovery of new *C. elegans* dauer pathway genes is surprising given the extensive genetic screens that have been conducted(3, 5, 30–32). However, genes that require a complete deletion to confer a dauer phenotype, or genes that control development through complex and antagonistic tissue specific effects, as possible for *ilcr-2*, could have been missed. In this regard, two other deletions of the *ilc-17.1* exist in the community (*tm5218 and tm5124*). These do not arrest as dauers, but are also not null alleles and express exons 1 and 2 of *ilc-17.1* transcript (Supplementary Figure S14c). However, the *tm5218* deletion removes the predicted beta strands required for the canonical cysteine knot fold of IL-17s and might have arguably been expected to be result in the loss of ILC-17.1 signaling capability, mimic *ilc-17.1* deletion mutants, and arrest as dauers. Although this is not the case, and all *tm5218* homozygotes escape dauer arrest, a variable, but significant percentage of trans-heterozygotes for *ilc-17.1(tm5218)* and *ilc-17.1*(*syb5296*) larvae arrest as dauers, as confirmed by PCR genotyping, indicating non-complementation (Supplementary Figure S14d; results from one experiment; dauer arrest varied between 0-75% of F1 trans-heterozygotes; n>10 experiments). This observation adds confidence to the role of *ilc-17.1* in dauer, but also reinforces our lack of understanding of the nature and stoichiometry of the IL-17 homo/heterodimers and IL-17-ILCR-2 signaling assemblies in *C. elegans*.

We hypothesize that ILC-17.1 is secreted upon sensory neuronal activity to control CEP-1/p53 activity, but this remains to be shown, and the neuronal circuits identified(93). Nutrient scarcity has shaped much of evolution, and the ability of cells and organisms to sense nutrient availability and prepare in anticipation to pause cell cycle progression and growth and maintain a quiescent state until resources are optimal, has unquestionable selective advantages. For *C. elegans*, like for most organisms, the availability of food is not guaranteed. Neuronal ILC-17.1 signaling appears to be one mechanism which allows the animal to match the cell fate programs and metabolic requirements of the developing larvae with resource availability. Neurons also control dauer entry though TGFβ and insulin signaling. In addition, other stress-responsive transcriptional programs that can be activated autonomously by cells, are also controlled through neuronal signaling in *C. elegans*, allowing the nervous system to coordinate tissue-specific transcriptional and epigenetic responses with organismal physiology and behavior(94–97). The extent to which IL-17s function similarly in mammals remains to be understood (98, 99).

## Materials and Methods

Descriptions of strains and methods are included in Supporting Information (SI), Extended Methods. Strains will be available upon request, after publication. The data discussed in this publication have been deposited in NCBI’s Gene Expression Omnibus (Edgar et al., 2002) and are accessible through GEO Series accession number GSE218596, https://www.ncbi.nlm.nih.gov/geo/query/acc.cgi?acc=GSE218596 and GSE229132, https://www.ncbi.nlm.nih.gov/geo/query/acc.cgi?acc=GSE229132

## Acknowledgements

We thank the V.P. laboratory, Drs. Sarit Smolikove, Josep Comeron, Anna Malkova and Michael Petrascheck for comments, Dr. Peter Ratcliffe, Oxford, for his generous gift of *CeHIF-1* antibody, and Dr. Mario de Bono, Institute of Science and Technology Austria (ISTA) for *C. elegans* strains. Nematode strains were provided by the Caenorhabditis Genetics Center (CGC) (funded by the NIH Infrastructure Programs P40 OD010440). This work was supported by NIH R01 AG060616 (V.P.).

