## Supplementary material for "Neuronal IL-17 controls *C. elegans* developmental diapause through CEP-1/p53": Methods supplementary figures and supplementary figure legends

**Extended Methods**

**Materials and Methods**

***C. elegans* strains**

*C. elegans* strains used in this study are listed in Table 1. Strains were procured from Caenorhabditis Genetics Center (CGC, Twin Cities, MN), generated in the laboratory or generated by Suny Biotech (Suzhou, Jiangsu, China 215028). All strains will be available upon request, after publication.

| **Strain Name** | **Gene name/genotype** | **Source** | **Additional information** |
|---|---|---|---|
| N2, *C. elegans* var *Bristol* | N2, Wild type | Caenorhabditis Genetics Center | |
| TJ1 | *cep-1 (gk138)* I | Caenorhabditis Genetics Center | |
| CF1038 | *daf-16 (mu86)* I | Caenorhabditis Genetics Center | |
| TG12 | *cep-1*(lg12501) I*; unc-119* (ed4) III; *gtIs1* [CEP-1::GFP + *unc-119* (+)] | Caenorhabditis Genetics Center | *cep-1p*::CEP-1(let-585 3'UTR)::GFP; from construct pRH53 |
| CB1370 | *daf-2 (e1370)* III | Caenorhabditis Genetics Center | |
| VEP024 | praEx021 (*ilc-17.1*p::*ilc-17.1* cDNA::3XFLAG::*ilc-17.1* 3'UTR; p*myo-2*::mCherry::*unc-54* 3'UTR) | Prahlad Lab | ILC-17.1 overexpression under its own |

| | | | |
|---|---|---|---|
| | | | promoter and 3’UTR |
| VEP031 | praEx022 (*unc-54*p::*ilc-17.1* cDNA::3XFLAG::*tbb-2* 3’UTR; p*dat-1*::GFP::*unc-54* 3’UTR) | Prahlad Lab | ILC-17.1 overexpression under the muscle promoter and 3’UTR |
| VEP032 | *ilc-17.1 (syb5296)* X | Prahlad Lab/ SunyBiotech | *ilc-17.1* deletion, 2173bp deletion, and the 15bp and 127bp sequences were left in the 5' and 3' deletion end, respectively of the 2315 bp *ilc-17.1* gene |
| VEP033 | *ilc-17.1 (syb5296)* X; praEx022 (*unc-54*p::*ilc-17.1* cDNA::3XFLAG ::*tbb-2* 3’UTR; p*dat-1*::GFP::*unc-54* 3’UTR) | Prahlad Lab | ILC-17.1 paracrine rescue of *ilc-17.1* deletion |
| VEP034 | *ilc-17.1 (syb5296)*X; *cep-1 (gk138)* I | Prahlad Lab | |
| VEP035 | *ilc-17.1 (syb5296); daf-16 (mu86)* I | Prahlad Lab | |
| VEP036 | *unc-119 (ed4)*; *gtIs1* [CEP-1::GFP + *unc-119* (+)] | Prahlad Lab | CEP-1 overexpression (ref: (61)) |

| | | | |
|---|---|---|---|
| VEP037 | *ilc-17.1 (syb5296)*; *praEx022* (*unc-54*p::*ilc-17.1* cDNA:: 3XFLAG::*tbb-2* 3'UTR; p*dat-1*::GFP::*unc-54* 3'UTR); *unc-119* (ed4); gtIs1 [CEP-1::GFP + *unc-119* (+)] | Prahlad Lab | IL-17.1 overexpression; CEP-1/p53 overexpression |
| VEP038 | *ilc-17.1 (syb5296); praEx022* (*unc-54*p::*ilc-17.1* cDNA:: 3XFLAG::*tbb-2* 3'UTR; p*dat-1*::GFP::*unc-54* 3'UTR); *daf-2* (e1370) III | Prahlad Lab | |
| VEP039 | *daf-2 (e1370)* III; *cep-1 (gk138)* I | Prahlad Lab | |
| VEP040 | *cep-1 ((syb6099)* [*cep-1*::3XFLAG]) I | Prahlad Lab/ SunyBiotech | Endogenous *cep-1* 3X FLAG tagged at C-terminus |
| VEP041 | *ilc-17.1 (syb5296)* X ; *cep-1 (pra02* [*cep-1*::3XFLAG]) I | Prahlad Lab | |
| VEP042 | *praEx022 (unc-54p::ilc-17.1* cDNA::3XFLAG::*tbb-2* 3'UTR; p*dat-1*::GFP::*unc-54* 3'UTR); *ilc-17.1* (syb5296) X ; *cep-1* (pra02 [cep-1::3XFLAG]) I | Prahlad Lab | |
| VEP043 | *ilc-17.1 (pra03 [ilc-17.1::*SL2::mCherry]) X | Prahlad Lab/ SunyBiotech | |
| VEP044 | *ilc-17.1* (*pra04* [*ilc-17.1*::3xHA]) X | Prahlad Lab/ SunyBiotech | |
| VEP045 | *ilcr-2* (*pra05* [*ilcr-2*::SL2::GFP]) II | Prahlad Lab/ SunyBiotech | |
| CB1386 | *daf-5 (e1386)* II | Caenorhabditis Genetics Center | |

| | | | |
|---|---|---|---|
| VEP061 | *ilc-17.1 (syb5297)* | Prahlad Lab/ SunyBiotech | *ilc-17.1* deletion, 2188bp deletion, and the 9bp and 118bp sequences were left in the 5' and 3' deletion end, respectively of the 2315 bp *ilc-17.1* gene |
| DV3805 | *reSi7[rgef-1p::TIR1::F2A::mTagBFP2::AID*::NLS::tbb-2 3'UTR] I:-5.32* | Caenorhabditis Genetics Center | |
| LP870 | *cpSi172 [myo-2p::TIR1::F2A::mTagBFP2::AID*::NLS::tbb-2 3'UTR] I:-5.32* | Caenorhabditis Genetics Center | |
| DV3799 | *reSi1[col-10p::TIR1::F2A::mTagBFP2::AID*::NLS::tbb-2 3'UTR] I:-5.32* | Caenorhabditis Genetics Center | |
| DV3801 | *reSi3[unc-54P::TIR1::F2A::mTagBFP2::AID*::NLS::tbb-2 3'UTR] I:-5.32* | Caenorhabditis Genetics Center | |
| JDW225 | *wrdSi23[eft-3p::TIR1::F2A::mTagBFP2::AID*::NLS::tbb-2 3'UTR] I:-5.32* | Caenorhabditis Genetics Center | |

| | | | |
|---|---|---|---|
| JDW221 | *wrdSi18[mex-5p::TIR1::F2A::mTagBFP2::AID*::NLS::tbb-2 3'UTR] I:-5.32* | Caenorhabditis Genetics Center | |
| PHX7467 | *ilcr-2::AID (syb7467)* II | Prahlad Lab/ SunyBiotech | |
| | *ilc-17.1 (tm5218)* X | C. elegans National Bioresource Project, Japan | |
| | *ilc-17.1 (tm5124)* X | C. elegans National Bioresource Project, Japan | |
| | *daf-11(ks67)* | Dr. Gidalevitz (Drexel University) | The original strain was daf-11(ks67); pdaf-7::GFP/plin44::GFP. Non-GFP mothers were selected. |

**Generation of CRISPR strains**

CRISPR/Cas9 was used to create following *C. elegans* strains:

| **Background Strain** | **Type of editing** | **Description/position of editing** | **Gene** | **Resulting strain** |
|---|---|---|---|---|
| N2, *C. elegans* var Bristol | Deletion (*syb5296*) | 2173bp deletion, and the 15bp and 127bp sequences were left in | (*ilc-17.1*)X | VEP032 |

| | | | | |
|---|---|---|---|---|
| | | the 5' and 3' deletion end, respectively of the 2135 bp *ilc-17.1* gene. **sgRNA used - <u>CCA</u>AAATCACCACACAGGACAAA** | | |
| N2, *C. elegans* var Bristol | Deletion (*syb5297*) | 2188bp deletion, and the 9bp and 118bp sequences were left in the 5' and 3' deletion end, respectively of the 2135 bp *ilc-17.1* gene. **sgRNA used- tagGCAAATGCGAATGCGGA<u>TGG</u>** | (*ilc-17.1*)X | VEP032 |
| N2, *C. elegans* var Bristol | Insertion; *SL2::mCherry* | C'-terminus | (*ilc-17.1*)X | VEP043 |
| N2, *C. elegans* var Bristol | Insertion; 3X HA | C'-terminus | (*ilc-17.1*)X | VEP044 |
| N2, *C. elegans* var Bristol | Insertion; SL2::GFP | C'-terminus | (*ilcr-2* )II | VEP045 |
| N2, *C. elegans* var Bristol | Insertion; 3X FLAG | C'-terminus | (*cep-1*)I | VEP040 |
| N2, *C. elegans* var Bristol | Deletion (1475bp of 1692 bp) | Prahlad Lab/ SunyBiotech | F25D1.3 (*syb7367)* V | PHX7367 |

| | | | | |
|---|---|---|---|---|
| N2, *C. elegans* var Bristol | Insertion; AID tag | C'-terminus | (*ilcr-2* )II | PHX7467 |

**Generation of transgenic strains**

**i) Backcrossing of *ilc-17.1 (syb5296)* X**

The *ilc-17.1 (syb5296)* X strain was crossed to wild-type (N2) males, and the male F1 progeny which harbored the *ilc-17.1 (syb5296)* allele on their X chromosome were backcrossed to the *ilc-17.1 (syb5296)* X strain and homozygous F2s were selected by PCR. This procedure was conducted 2X times to generate the 2X backcrossed *ilc-17.1 (syb5296)* X line.

**ii) Generation of *cep-1* overexpression strain**

TG12 (*cep-1*(lg12501) I; *unc-119* (ed4) III; [CEP-1::GFP + *unc-119* (+)]) is a strain that expresses a functional CEP-1 tagged with GFP integrated into its genome, as determined by the rescue of *cep-1*(lg12501) phenotype(63). To overexpress CEP-1 we backcrossed the TG12 strain with wild-type (N2) worms and selected F2 progeny that were homozygous for the wild-type *cep-1* gene, lacked the *cep-1*(lg12501) I mutation (confirmed by PCR), and were homozygous for the CEP-1::GFP transgene (confirmed by 100% GFP expression amongst the F3 and F4 progeny). Overexpression of cep-1 mRNA was verified by qPCR (Supplementary Figure S6d). The CEP-1::GFP construct was PCR amplified from the final transgenic VEP037 *C. elegans* strain, and sequence verified.

**iii) Generation of ILC-17.1 overexpression strain under its own promoter and 3'UTR**

To generate the strain VEP024 – praEx021 [(*ilc-17.1*p::*ilc-17.1* (cDNA)::3xFLAG::*ilc-17.1* 3'UTR); p*myo-2*::mCherry::*unc-54* 3'UTR], which overexpressed *ilc-17.1* under the endogenous *ilc-17.1* promoter and *ilc-17.1* 3'UTR, we first amplified 3 kb of genomic sequence upstream of the translational start site (*ilc-17.1* promotor) and 1 kb of genomic sequence downstream from the translational stop codon (*ilc-17.1* 3' UTR). These regions were cloned using Gateway technology into pDONR221. *ilc-17.1* cDNA fused to C-terminal 3xFLAG sequence was synthesized as a gBlock (Integrated DNA technologies). These three fragments were then submitted for gene synthesis service through GenScript to generate the expression vector pUC57(*ilc-17.1*p:: *ilc-17.1* (cDNA)::3xFLAG:: *ilc-17.1* 3'UTR). All plasmids were sequence verified. The *ilc-17.1* expression vector was then injected at 97.5 ng/ul along with the co-injection marker pCFJ90 (p*myo-2*::mCherry::*unc-54* 3'UTR) at 2.5 ng/ul by InVivo Biosystems injection express service. Animals expressing mCherry were singled, lines transmitting the extrachromosomal array were established, and mCherry positive progeny were PCR verified to ensure they were transmitting *ilc-17.1*p:: *ilc-17.1* (cDNA)::3xFLAG:: *ilc-17.1* 3'UTR. These lines were then harvested for Western blot to verify expression of protein.

**iv) Generation of ILC-17.1 overexpression strain under the *unc-54* muscle promoter and *tbb-2* 3'UTR**

To generate VEP031, we overexpressed ILC-17.1 under muscle promoter by fusing - [(*unc-54*p:: *ilc-17.1* (cDNA)::3xFLAG] which was synthesized as a gBlock (Integrated DNA technologies) and *tbb-2* 3' UTR which was amplified from genomic DNA. The two fragments were cloned into pUC19 plasmid as the backbone using Gibson assembly(113) to create the overexpression plasmid [(*unc-54*p:: *ilc-17.1* (cDNA)::3xFLAG::*tbb-2* 3'UTR); p*dat-1*::ssmito] and sequence verified. The construct was injected at 100 ng/ul along with the co-injection marker that expressed GFP in the dopaminergic neurons under a *dat-1* promoter [ p*dat-1*], at 10 ng/ul into wild-type (N2) worms. Animals expressing GFP in their dopaminergic neurons were singled, lines were established, and GFP-positive progeny were PCR verified to ensure that they were also transmitting harbored the *ilc-17.1* transgene. These lines were then harvested for Western blot to verify expression of protein.

**v) Generation of crosses.**

**Generating trans-heterozygotes with *ilc-17.1 (syb5296)***

To obtain trans-heterozygotes to test for complementation, *ilc-17.1 (sy5296); dat-1p::GFP* males or hermaphrodites were mated with of *ilc-17.1 (tm5218).* Because both self- and cross-progeny of *ilc-17.1 (sy5296); dat-1p::GFP* hermaphrodites mated with of *ilc-17.1 (tm5218)* males would be GFP positive, all GFP positive progeny were also genotyped. The strain *ilc-17.1 (sy5296); dat-1p::GFP* was used, instead of the mutant *ilc-17.1 (sy5296)* alone, to ensure mating and identify cross progeny. Animals were allowed to mate at 20°C for 2 days, mated hermaphrodites were allowed to lay eggs, the eggs of the mated hermaphrodites were placed at 25°C. The F1 progeny were scored for the presence of GFP and whether they were dauers or L4s/adults. GFP positive F1's were then singled and lysed for genotyping by PCR. Similar experiments were conducted without *dat-1p::GFP* and we obtained similar results. The following primers were used:

VP14-ko-s: AAAGTTGAGCTCAAACTGGG

VP14-ko-a: TACAGATCACGTGAACGAAG

The remaining crosses between strains were generated according to standard procedures. The primers used to genotype all strains are included in the table below.

| | Forward primer | Reverse primer |
|---|---|---|
| VEP032 | TGAATGAGCAAATCGACCAG | AGTTTTGAGATTGAGGGCAAA |
| | CAGGTCAAGGGGCTAAAACA | CCCACATTAATACTCGCACCT |
| CF1038 | CGGATCAGGAAGAACCAGAA | CATTCCCCGAGTACGAGCTA |
| | AAGCTGCTGCCTTCACTCTC | ATGGGACTCGAGTTGAGTGG |
| TJ1 | CGGTCTTTTTGGACGATGAA | GTAAACATTGCCGCGTTTTT |
| | GCGATTCGAAATCCAAGAAG | AGGAACGCATCCATGAAGAC |
| TJ12 | GCCTTCCTCTATTCACTTGAAAC | ACCGAGAGCTCCATGAAGAA |
| | TCACTCTTTGAATTGGTTCTTTTG | GCTTACGTTCGATTGGCTTC |
| | AAAAACGCGGCAATGTTTAC | GAGCTTTGTGGGAAGCAATC |
| RB2075 | GACAGTTACGCTGTGCTTGG | CTCCCGGGGTATTCTGAAAT |

| | | |
|---|---|---|
| | GGTGATTCTCCCGTTGGTAA | AAACGGAAATGGCAGAGAGA |
| CB1370 | TAA ATC TCG GAA TTC TGG TGA A | GGA TGA GAC TGT CAA GAT TGG A |
| | AAG ATG AAC GAG CTG GAG GA | AGA TTG GAG ATT TCG GAA TGG |
| VEP040 | AGA ATT CGA GAA ACA TAA TGA CAT CTT | AAA AAT GCG ATT ATA AAA AGG AAC A |
| CB1386 | CGG CTT GAT CGG ATA GTG AT | AAG AGC AAG GAG TGC GAA AA |
| | Wild-type PCR product was digested by BstNI | |
| PHX7467 | GAAACGAGATGGAAACCGAA | TTGCTTCTCTCACTGTTGCC |

**Growth conditions for *C. elegans* strains**

All strains were grown and maintained at 20°C unless otherwise mentioned. Animals were grown in 20°C incubators (humidity controlled) on 60mm nematode growth media (NGM) plates by passaging 8-15 L4s (depending on the strain) onto a fresh plate. Extra care was taken to ensure equal worm densities across all strains. Animals were fed *Escherichia coli* OP50 obtained from Caenorhabditis Genetics Center (CGC) that were seeded ($OD_{600}$=1.5 and this was strictly maintained throughout the experiments) onto culture plates 2 days before use. The NGM plate thickness was controlled by pouring 8.9ml of autoclaved liquid NGM per 60mm plate. Laboratory temperature was maintained at 20°C and monitored throughout. For all experiments, age-matched day-one hermaphrodites, or larvae timed to reach specific developmental stages as mentioned in the figure legend, were used.

For all experiments with larvae, the timings were adjusted to account for the differences in growth rates at different temperatures. Thus, at 25°C, larvae were harvested 30-36 hrs. post bleach hatch or egg lay, and at 20°C, larvae were harvested 40 hrs. post bleach hatch or egg lay.

**Obtaining synchronized embryos by 'bleach-hatching'**

Bleach-hatching' was performed as previously described(107). Populations of 250-300 gravid adults were generated by passaging L4s, as described above. These plates were used for obtaining synchronized embryos by bleach-induced solubilization of the adults. Specifically, animals were washed off the plates with 1X PBS and pelleted by centrifuging at 2665Xg for 30s. The PBS was removed carefully, and worms were gently vortexed in the presence of bleaching solution [250µl 1N NaOH, 200µl standard (regular) bleach and 550µl sterile water] until all the worm bodies had dissolved (approximately 5-6 minutes), and only eggs were viable. The eggs were pelleted by centrifugation (2665Xg for 45s), bleaching solution was carefully removed and then embryos were washed with sterile water 3-4 times and counted under the microscope. The desired number of embryos were seeded on fresh OP50 plates and allowed to grow at 20°C or 25°C for specific time periods depending on the experimental need. If >5% eggs remained unhatched, these plates were discarded.

**Dauer Assay**

Embryos were allowed to hatch and grow on OP50 plates at 20°C or 25°C for this assay. Embryos were generated according to one of the following two methods: (i) Day-one gravid adults that had grown under normal culture conditions ( on OP50 at 20°C ) were bleach dissolved as described above, and ~50-100 embryos obtained from these gravid adults were seeded on fresh OP50 plates or (ii) day-one gravid adults were allowed to lay eggs on fresh OP50 (or RNAi) plates for 2-4 hours at 20°C, the adults were removed, and then embryos were allowed to develop at 20°C or 25°C under humified condition for 48-72 hrs. The former, bleach-synchronization method was used for RNA-seq method and experiments requiring large numbers of age-matched larvae. The latter, timed-egg lay method was used for all other experiments. For each of the strains and experiments, when we had to use bleach synchronization, we first ensured that bleaching alone did not alter the percentage of dauers or gene expression profiles by conducting a pilot experiment using both methods side-by-side. This was the case for all experiments, except experiments where glucose rescued the dauer phenotype. Here, although the trends were the same, and glucose resulted in a significant rescue of ilc-17.1 deletion mutants and cep-1 OE larvae that were generated either by bleach-synchronization or timed-egg lay, bleach synchronization caused a lower percentage to be rescued from dauer arrest. To assess dauer entry, the number of larvae that arrested as dauers (determined by phenotype and/or resistance to 1%SDS) and those that developed into L4s or adults were counted and percentage of dauers was calculated. The total number of embryos in each plate was ~50-100. Each experiment was performed in triplicate or more.

For the *ilc-17.1* deletion mutants, in order to ascertain that the dauer phenotype was not simply due to the presence of a background mutation, we also conducted the dauer assay after backcrossing the *ilc-17.1* (syb5296) X animals with wild-type Bristol N2 animals, 2X times. The backcrossed strains also arrested as dauers under those same conditions and 100 ± 0% larvae (n=3 independent repeats, 212 larvae scored), arrested as dauers at 25°C as determined by SDS resistance compared to 0% of wild-type larvae.

**SDS treatment**

Larvae were washed off the dauer assay plates, washed 1X with PBS to remove bacteria, treated with 1% sodium dodecyl sulfate (SDS) for 30 minutes, washed again with M9, and then transferred onto a fresh plate. Larvae could be scored easily as dead/dissolved or as live dauer larvae (based on their phenotype, and movement).

**Auxin treatment**

Auxin treatment for auxin-induced degradation experiments were performed by transferring 10 synchronized day 1 adult worms to OP50 bacteria-seeded plates, or RNAi plates seeded with

HT115 (L4440), containing auxin. There were allowed to egg lay for 4 hrs. and then removed from the auxin plates. These plates were place at 25°C and dauers were scored at 48-72 hrs.

Auxin plates were made at 4mM final concentration using auxin indole-3-acetic acid (IAA) (Alfa Aesar #A10556). A 400 mM stock solution in ethanol was prepared and added to NGM agar, cooled to about 55°C, before pouring plates. Plates were dried for two days and 1.5 OD op50 bacteria was added on top. Plates were left at room temperature for 2 days to allow bacterial lawn growth before adding the worms for egg lay.

Efficacy of AID was evaluated by measuring Blue Fluorescence Protein (BFP) fluorescence in a population of larvae that were exposed to auxin. Only those larvae where BFP was no longer detectable in all tissues where BFP was expressed, under an inverted confocal microscope under the 63X objective, were scored as positive.

**RNAi mediated downregulation of genes**

All RNAi clones used were verified by sequencing, and plates were seeded with RNAi bacteria for a maximum of two days before being used. Day-one gravid adults grown on OP50 were allowed to lay eggs on RNAi-bacteria seeded plates for 2-4 hours at 20°C. A second generation of RNAi knockdown was performed where necessary by transferring L4's on existing RNAi plates to fresh plates for knockdown in their progeny. The adults were allowed lay eggs and then removed, and the embryos were transferred to 25°C (or kept at 20°C) under humified condition for 48-72 hrs. The number of larvae that arrested as dauers (determined by phenotype and/or resistance to 1%SDS) and those that developed into L4s or adults were counted and percentage of dauers was calculated.

**Exposure to food [OP50] availability**

Gravid day-one *ilc-17.1*(pra03 [*ilc-17.1*::SL2::mCherry]) X adults or *ilc-17.1*(pra04 [*ilc-17.1*::3xHA]) X adults were bleach dissolved and embryos were placed either on empty plates with no food or NGM plates seeded with OP50 and allowed to grow for 24-36 hours at 25°C. To observe mRNA expression of *ilc-17.1, ilc-17.1*(pra03 [*ilc-17. 1:*SL2::mCherry]) X larvae were anesthetized with 1mM levamisole, mounted on 1% agarose pads, and imaged using a Confocal SPE microscope using optimal settings. mCherry expression served to mark cells that expressed *ilc-17.1* mRNA. To identify where ILC-17.1 protein was expressed prior to food exposure, and after food exposure, 24–36-hour *ilc-17.1*(pra04 [*ilc-17.1*::3xHA]) X larvae were fixed and immunostained using Bouin's Tube Fixation method (see below).

**Immunofluorescence**

Nematodes were fixed and immunostained using a modification of the Bouin's Tube Fixation method (114). Worms were fixed for 30 min. at room temperature (RT) in 400 µl Bouin's fix (Sigma Aldrich) + 600 µl methanol and 10 µl β-mercaptoethanol by tumbling, freeze cracked three times in liquid nitrogen and again tumbled for 30 mins at RT. For permeabilization, the fixative was removed and exchanged for borate-Triton-βmercaptoethanol (BTB: 1xBorate Buffer, 0.5% Triton and 2% β-mercaptoethanol) solution. Worms were tumbled 3 times for 1 hour each in fresh BTB solution at RT. Worms were washed with PBS-0.05% Tween and incubated in block-solution (5% BSA). Staining with primary antibody was performed overnight at 4 °C, incubation with secondary antibody (Donkey anti-Mouse Alexa Fluor 488, 1:2000) for 2-4 hours at RT. All antibody dilutions were performed using antibody buffer containing 5% BSA. Samples were mounted onto glass slides using VECTASHIELD antifade mounting medium (Vector Laboratories, Burlingame, CA, USA). Primary antibodies used: (i) Monoclonal ANTI-FLAG M2, Sigma Aldrich, 1:500, (ii) Mouse anti-HA antibody, Thermofisher; 2-2.2.14, at 1:1000), (iii) anti β-Actin, Cell Signaling Technology, #4967, 1:1000.

**Chemotaxis**

Chemotaxis assay was performed on 9 cm petri dishes containing NGM. Two marks were made on the back of the plate at opposite sides of the plate about 0.5 cm from the edge of the agar. About 5 μl of attractant diluted in water was placed on the agar over one mark, and 5 μl of water was placed as the control over the opposite mark. Attractants and concentrations used are listed in drugs and metabolites section. 5 ul sodium azide with the concentration of 1M was also placed at both the attractant source and the control source. This drug could anesthetize animals within about a 0.5-cm radius of the attractant. Age synchronized day-one adult worms were transferred to the middle of the NGM plates at a point equidistant from the middle of each odorant. After 1 hr., the assay was quantified by counting the number of worms that had left the center origin a chemotaxis index was calculated [#Odor - #Control] / [#Odor + #Control]. Each repeat consisted of 50 worms, and experiments were repeated a minimum of three times.

**Measuring Pharyngeal pumping**

Synchronized day-one adults were singled onto NGM plates seeded with OP50, and onto plates without OP50. Pharyngeal pumping rates were determined by recording the pharyngeal region of animals by video using a Leica S9i digital stereo microscope at 5X magnification and slowing down the video to manually count the number of 'pumps' in 10 seconds, three times per animal. The mean of these pumps was determined and the number of pumps/minute calculated. One complete cycle of synchronous contraction and relaxation of the corpus and the terminal bulb was counted as a pump.

**Feeding of fluorescent latex beads**

Overnight 5ml OP50 culture LB was pelleted and resuspended in fresh 0.5ml LB to concentrate bacteria. 1 μl of fluorescent beads of 0.5μm mean particle size that mimic size of *E. coli*. (Sigma L3280, red fluorescence) was added to 1 ml of concentrated OP50 [1:1000 ratio (v/v)], and 100ul of the mixture was seeded onto NGM plates (115). Synchronized day-one adults were allowed to lay eggs on the bead containing plates for 2 hours at 20°C and then removed. After 40 hours, larvae were picked onto 1% agarose pads on glass slides and anesthetized using 10mM levamisole. Z-stack images of the larvae were taken on Leica TCS SPE confocal microscope and number of beads within a set area near the tail region per larvae was quantified.

**Drug and metabolite treatment**

The following drugs or metabolites were used:

| Drug/metabolite | Source | Final concentration |
|---|---|---|
| Glucose | Research products International | 50/100mM |
| 2-Deoxy-d-glucose | R&D systems | 50mM |
| Mitotempo | Millipore sigma | 0.1mM |
| L-Glutamine | Sigma-Aldrich | 0.26mM |
| Skim Milk powder | HyVee - instant nonfat dry milk | 20mM |
| Diacetyl | Sigma-Aldrich | 0.01% v/v |
| Lysine | Sigma-Aldrich | 1M |
| (25S)-Δ7-Dafachronic Acid | Cayman Chemical | 50nM |
| 8-Bromo-cGMP | Cayman Chemical | 5mM |

The drug solution (0.5 ml) was spread onto NGM agar plates containing an OP50 lawn and left for 1 hr. to dry or used for chemotaxis assays as described. Age synchronized day-one adult worms were transferred to the middle of the NGM plates for 4 hrs. and dauer assay were performed as described. Sterile water was used as control.

8-Bromo-cGMP solution was first added to standard NGM plates and allowed to dry for 1 hour. 1.5 OD op50 was added on top and allowed to grow for 1 day before starting the dauer assay as described.

Plates containing D-glucose were prepared as described in published methods (100), except previously except that we seeded live OP50 onto Glucose plates. Stock solutions of 0.1 and 1M Sterile D-glucose was prepared and added into NGM agar to obtain final concentration of 5mM and 50mM, respectively. For control plate, equivalent amount of sterile water was added to NGM agar. The plates were left to dry for 2 days at RT before seeding 1.5 OD OP50 onto them. After drying at RT for another 1day, dauer assays were performed as described.

**Fluorescence Image analysis**

ImageJ (Fiji) v1.53i was used for measuring fluorescence intensity as follows: single planes [, DAF-16::GFP, CEP-1::GFP, TG12] or projections of z-planes [DAF-28, INS-4, fluorescent latex beads,pax-3p::mCherry] were used for measuring fluorescence intensity. The region of interest was circled using the circle selection tool. The mean fluorescence intensity of each circled area was recorded using the measurement tool. When appropriate, the background fluorescence intensity was measured and subtracted. Quantification of measurements was done in Microsoft excel/GraphPad.

**RNA-sequencing and Data analysis**

a) **RNA isolation, library preparation and sequencing**

For the Total RNA-seq samples: Day-one adult worms were bleach-hatched and ~3200 eggs/genotype (~800 eggs/plate and 4 plates/genotype) were seeded on fresh OP50 plates and allowed to grow for 30-34 hrs. at 25°C. Worms were washed with sterile water and total RNA was extracted from biological triplicates using the Direct-zol RNA Miniprep Kits (catalog no. R2050, Zymo Research). Libraries were prepared using the Illumina Stranded Total RNA Prep with Ribo-Zero Plus, for rRNA depletion (catalog no. 20040525, Illumina). Samples were sequenced in one lane of the Illumina NovaSeq 6000, generating 2x150bp paired-end reads.

For the mRNA-seq samples: Day-one adult worms were bleach-hatched and ~3200 eggs/genotype (~800 eggs/plate and 4 plates/genotype) were seeded on fresh OP50 plates and allowed to grow for either 15 hours or 32 hours at 25°C. Worms were washed with sterile water and total RNA was extracted from biological triplicates using the Direct-zol RNA Miniprep. Libraries were prepared using Illumina TruSeq Stranded mRNA kit. Samples were sequenced in one lane of the Illumina NovaSeq 6000, generating 2x100bp paired-end reads.

**b) RNA-seq analysis**

The quality of the RNA sequences was assessed with FastQC. Adapters and sequence reads with a quality lower than Q25 were trimmed by using Trimgalore (116)(v0.67). Ribosomal RNA (rRNA) contamination was filtered with sortmeRNA(117) (version 4.3.4). Sequencing reads were aligned to the *C. elegans* genome (WBcel235(118)) using the Star aligner(119) (v2.7.9a) and then the aligned reads were quantified with FeatureCounts v2.0.1 from the R package Rsubread(120) . Differential expression analysis was performed using DESeq2(121) and genes with FDR corrected p-value of < 0.05 were considered significant. Pairwise distance analysis (sample-to-sample) was performed by using normalized counts coupled with the variance stabilization transformation (VST) on the complete set of genes and calculating the Euclidean distance between the replicates. Principal Component Analysis (PCA) was done using the plotPCA function in DESeq2(121) with the VST transformed data. The mRNA-seq was analyzed following a similar protocol as described above, excluding the SortmeRNA step.

The microarray data for the *C. elegans* dauer stage was obtained from the Supplementary Table 1 of the study by Wang, J. & Kim (2003)(38). The expression values of the dauer stage were defined as the dauers expression (fold-change) at 0hr relative to the mixed staged reference RNA. Sequences IDs from the array were converted to standard gene nomenclature using Simplemine in Wormbase(122). The correlation between the expression at the dauer stage and the fold change in expression in the *ilc-17.1* deletion mutant relative to the wild type, was determined by performing a Spearman Rank Test.

The expression of C. elegans larvae (L1) treated with ascaroside cocktail was obtained from Cohen, Sun, Schroeder and Sternberg (123). The correlation between the Fold-Change in expression in the larvae treated with ascaroside relative to N2 larvae and the fold change in expression in the *ilc-17.1* deletion mutant relative to the wild type, was determined by performing a Spearman Rank Test. The differentially expressed genes (DEGs) between N2 and daf-1(m40) was obtained from. Hu, Crossman, Prasain, Miller and Serra (124) and were presented as $Log_2$(Fold-Change) values. The overlap between differentially expressed genes (DEGs) in the cep-1 overexpression mutant and the *ilc-17.1* deletion mutant was tested using Fisher exact test from the Gene Overlap package(125). Venn diagram was done using the R package VennDiagram(126).

**c) Functional analysis**

Gene ontology analysis (GO) and KEGG enrichment analysis was performed on the differentially expressed genes by using ClusterProfiler 4.0(127). Terms with an FDR corrected p-value <0.05 were considered significant. The annotations were obtained from R package org.Ce.eg.db(128) (version 3.14) and the KEGG database(129). The annotations of genes as part of the metabolic pathways were obtained from the database Wormpaths(102) and heatmaps were done using the R packages Complexheatmap and pheatmap(130, 131).

**DiO dye filling assay to identify amphid neurons**

A stock dye solution containing 2mg/ml DiO (Molecular Probes, catalog # D-275) in dimethyl formamide is made and maintained under dark conditions. Day-one adult worms were transferred into an Eppendorf tube with 1 ml of M9, spun down at 2000-3000 rpm and supernatant removed. Worms were resuspended in 1 ml of M9 and 5 microliter DiO stock sol (1:200 dilution) was added and incubated on a slow shaker for 2 hours at room temperature. Worms were spun down and washed with M9 twice before transferring them onto agar pads with 1mM levamisole to visualize by fluorescence using a Leica TCS SPE confocal microscope.

**Dauer pharyngeal pumping**

Synchronized day-one adults were allowed to lay eggs on OP50 plates for 2 hours at 20°C and then removed. The plates were then placed at 25°C, and larvae were allowed to arrest as dauers or grow into L4 larvae (RNAi treatment) . On day 3, larvae that remained arrested as dauers were used to measure pharyngeal pumping rates using Zeiss Axio Observer.A1 at 40X magnification. WT (N2) L2 stage larvae were used to measure pumping rates as controls. Pumping rates were measured by counting the number of grinder movements in the terminal bulb per minute. The average of three biological replicates consisting of 10 dauer larvae each was quantified.

**Scoring apoptotic nuclei with Acridine Orange**

Acridine orange (AO) staining was performed as previously described (Hefel et al. 2021). AO staining to detect apoptotic germline cells was performed by picking 50 L4 animals to a fresh plate and allowing them to grow into day-one adults for 24 hours at 20°C. Day 1 adults were then either exposed to 75 Gy from a Cesium source or left unirradiated. 24 hours later, AO staining was performed. For each trial, 10 mg/ml AO stain was freshly prepared and then diluted 1:400 with M9 buffer. Worms were then picked to a tube of the diluted AO stain, wrapped tightly with foil, and rotated at room temperature for 1 hour. After mixing, worms were transferred to a fresh OP50 plate and allowed to crawl away from residual AO stain for 1 hour before being picked to a droplet of 10 mM levamisole on a 1% agarose pad. A cover slip was added, and worms were imaged immediately using the Leica fluorescence microscope on 60x magnification. It was important to visualize AO within 10 mins after mounting animals on the pad. The number of AO stain positive cells found in 12 individual gonad arms per experiment were recorded for each sample.

**Gamma irradiating *C. elegans* to activate CEP-1/p53**

100 L4 larvae were picked onto an OP50 seeded plate, allowed to mature into day-one adults and exposed to 75 Gy gamma irradiation, and subsequently harvested for Western blot analysis. To determine the number of AO positive apoptotic nuclei, at least 50 L4 larvae were exposed to 75 Gy gamma irradiation. The number of AO positive cells were scored 24 hrs. later, when animals had matured into day-one adults.

**Western blot analysis**

**i. Western blot analysis of *C. elegans***

Western blot analysis was performed with 30–100 adult day-one animals, or approximately 400 30-36 hr. larvae as indicated. Animals were harvested in 15 µl of 1X PBS (pH 7.4), and 4X Laemmli sample buffer (catalog no. 1610737, Bio-Rad) supplemented with 10% β-mercaptoethanol was added and samples were boiled for 30 min. Whole-worm lysates were resolved on 12% SDS-PAGE gels and transferred onto nitrocellulose membrane (catalog no. 1620115, Bio-Rad). Membranes were blocked with Odyssey Blocking Buffer (part no. 927–50000, LI-COR). Immunoblots were imaged using LI-COR Odyssey Infrared Imaging System (LI-COR

Biotechnology, Lincoln, NE). Mouse anti-FLAG M2 antibody (catalog no. F1804, RRID:AB_262044,_Sigma Aldrich) was used at 1:500 to detect CEP-1::FLAG. Mouse anti-HA antibody (Thermofisher; 2-2.2.14), was used at 1:1000 to detect ILC-17.1::HA. Mouse anti-α-tubulin primary antibody (AA4.3, RRID:AB_579793), developed by C. Walsh, was obtained from the Developmental Studies Hybridoma Bank (DSHB), created by the National Institute of Child Health and Human Development (NICHD) of the National Institute of Health (NIH), and maintained at the Department of Biology, University of Iowa. Phospho-AMPKα (Thr172) (40H9) (CST 2535) at 1:1000 dilution was used to detect AMPK. Phospho-p70 S6 Kinase (Thr389) Antibody (CST 9205S) at 1:1000 was used to detect phospho-S6 kinase. The following secondary antibodies were used: Donkey anti-mouse IgG (H and L) Antibody IRDye 800CW Conjugated (Licor) and Alexa Fluor 680 goat anti-rabbit IgG (H+L) (Invitrogen). LI-COR Image Studio software (RRID:SCR_015795) was used to quantify protein levels in different samples, relative to α-tubulin levels. Fold change of protein levels was calculated relative to wild type (N2)/ controls.

**ii. Western blot analysis of epithelial cell lines**

A549 epithelial cells were lysed in a modified M2 buffer containing 20mM tris[hydroxymethyl]aminomethane, 150mM NaCl at pH 7.4, 0.5% NP40, 3mM EDTA, 3mM EGTA, 4mM PMSF, and a cOmplete Mini, EDTA-free protease inhibitor tablet according to the manufacturer's instructions (Roche, #11836170001). Epithelial cells were collected by mechanical dislodgment in lysis buffer at individual time points, incubated for 20 minutes on ice, and spun at 14,000 x g for 5 minutes to remove insoluble material. 25ug protein was resolved by NuPAGE™ 4-12% Bis-Tris gel (Invitrogen) and transferred to PVDF for immunoblotting. Anti-p53 (Cell Signaling Technology, #9282) and β-actin (DSHB, 224-236-1) primary antibodies were used with HRP-conjugated secondary antibodies to anti-rabbit IgG (Jackson ImmunoResearch, #111-035-003) and anti-mouse IgG (Invitrogen, #62-6520). Densitometry was analyzed with ImageJ software.

**RNA extraction and real-time quantitative reverse-transcriptase PCR (RT-PCR)**

Worms were bleach-hatched as described above and ~400-800 eggs/plate (2 plates/strain) were seeded on fresh OP50 plates and were allowed to grow (i) for 30-34 hrs. at 25°C or (ii) for 36 hrs. at 20°C and then harvested for RNA extraction. RNA was extracted as described earlier (132). Briefly, plates were washed with sterile water and centrifuged. Water was carefully removed and 300 µl of Trizol (catalog no. 400753, Life Technologies) was added and snap-frozen immediately in liquid nitrogen. Samples were thawed on ice and then lysed using a Precellys 24 homogenizer (Bertin Corp.). RNA was then purified as detailed with appropriate volumes of reagents modified to 300 µl of Trizol. The RNA pellet was dissolved in 17 µl of RNase-free water. The purified RNA was then treated with deoxyribonuclease using the TURBO DNA-free kit (catalog no. AM1907, Life Technologies) as per the manufacturer's protocol. cDNA was generated by using the iScript cDNA Synthesis Kit (catalog no. 170–8891, Bio-Rad). qRT-PCR was performed using PowerUp SYBR Green Master Mix (catalog no. A25742, Thermo Fisher Scientific) in QuantStudio 3 Real-Time PCR System (Thermo Fisher Scientific) at a 10 µl sample volume, in a 96-well plate (catalog no. 4346907, Thermo Fisher Scientific). The relative amounts of mRNA were determined using the $\Delta\Delta C_t$ method for quantitation. We selected *pmp-3* as an appropriate internal control for gene expression analysis in *C. elegans*.

All relative changes of mRNA were normalized to either that of the wild-type control or the control for each genotype (specified in figure legends). Each experiment was repeated a minimum of three times. For qPCR reactions, the amplification of a single product with no primer dimers was confirmed by melt-curve analysis performed at the end of the reaction. Primers were designed using Primer3 software and generated by Integrated DNA Technologies. The primers used for the qRT-PCR analysis are listed below:

| **Gene** | **Forward primer (5'-3')** | **Reverse primer (5'-3')** |
|---|---|---|

| | | |
|---|---|---|
| *pfk-1.2* | TTGCATCGAATCTGTGAAGC | TCGCTGCAGTCAAAGCTAGA |
| *cyc-2.2* | CGGTGGAGCTATTCCAGAAG | GCAACTTGTCCGGATTGTCT |
| *egl-1* | TCCAAGCTAGCAGCAATGTG | GCGAAAAAGTCCAGAAGACG |
| *ced-13* | GCAACTCAAACACCGTTGAA | CAATGCTGGCATACGTCTTG |
| *phg-1* | CAGAGGAGCTTGTACCGACA | TCGTCTCTAGCAGTGCATGT |
| *sod-3* | CACTGCTTCAAAGCTTGTTCA | ATGGGAGATCTGGGAGAGTG |
| *mtl-1* | TGGATGTAAGGGAGACTGCAA | CATTTTAATGAGCCGCAGCA |
| *lys-7* | GCCGTCAAACTTGGCATCTT | GGGTTGTATGCACGAACGAA |
| *pmp-3* | TAGAGTCAAGGGTCGCAGTG | ATCGGCACCAAGGAAACTGG |
| *cep-1(a+b isoform)* | GCTCACTCTGTCGACTGCTGAGT | AACCCAAGTGTATCTGGGAACTTT |
| *cep-1(a isoform)* | GTTGTGCTCGACTCCCAAAAG | GGCACGCTTCTCAATTACAAGTT |
| *ilc-17.1* exon 1 | CACAGGACAAATTTACAGTC | GAATCCAGGGCGAGATGTG |
| *ilc-17.1* exon 2 | GTTGGAGGATACGAAGGGAAG | CTGCAACCTTAAGAGCGCTTG |

**Chromatin immunoprecipitation (ChIP)**

Chromatin immunoprecipitation (ChIP) was performed as described earlier (Das et al., 2020). Day-one adult worms were bleach-hatched as described above and ~1600 eggs were seeded (~800 eggs/plate and 2 plates/genotype) on fresh OP5O plates. Plates were kept at 25°C for 30-34 hrs. and larvae were washed with 1X PBS (pH 7.4) and cross-linked with freshly prepared 2% formaldehyde (catalog no. 252549, Sigma Aldrich) at room temperature for 10 min followed by addition of 250 mM Tris (pH 7.4) at room temperature for 10 min. Samples were then washed three times in ice-cold 1X PBS supplemented with protease inhibitor cocktail and snap-frozen in liquid nitrogen. The worm pellet was resuspended in FA buffer [50 mM HEPES (pH 7.4), 150 mM NaCl, 50 mM EDTA, 1% Triton-X-100, 0.5% SDS and 0.1% sodium deoxycholate], supplemented with 1 mM DTT and protease inhibitor cocktail. The suspended worm pellet was lysed using a Precellys 24 homogenizer (Bertin Corp.), and then sonicated in a Bioruptor Pico Sonication System (catalog no. B0106001, Diagenode) (15 cycles of 30 s on/off).

Endogenous CEP-1 was immunoprecipitated with anti-FLAG M2 magnetic bead (catalog no. M-8823, Sigma-Aldrich). Beads were first pre-cleared with salmon sperm DNA (catalog no. 15632–011, Invitrogen). Worm lysate was incubated at 4°C overnight with the pre-cleared FLAG beads. Beads were washed with low salt, high salt and LiCl wash buffers and then eluted in buffer containing EDTA, SDS and sodium bicarbonate. The elute was then de-crosslinked overnight in presence of Proteinase K. The DNA was purified by ChIP DNA purification kit (catalog no. D5205, Zymo Research). qPCR analysis of DNA was performed as described above using primer sets specific for different target genes. For all ChIP experiments, 10% of total lysate was used as 'input' and chromatin immunoprecipitated by different antibodies were expressed as % input values. The primers used for ChIP experiments, and the expected amplicon sizes are as follows:

| **Gene** | **Position** | **Forward primer (5'-3')** | **Reverse primer (5'-3')** | **Amplicon size** |
|---|---|---|---|---|
| *egl-1* | TSS +4 to +123) | CTCACCTTTGCCTCAACCTC | CGAGGAGAAGTCCTGAGACG | 120 bp |
| *ced-13* | Promoter (-428 to -310) | CTATTCTTGGCCGTGCTCAT | AGGCAATCTAGCATGCACCT | 119 bp |

| | | | | |
|---|---|---|---|---|
| *phg-1* | Promoter -428 to -294) | GCCAAACCTTCCAGA TTTACA | TTCCTAGATAAGGGT TAGATGATGAGA | 135 bp |
| *phg-1* | Intron 1 (+223 to +326) | AAGCTGAGCTCCGAA AACAA | TTTCCCGCTAAACGA GACAT | 104 bp |
| *cki-1* | Promoter (-698 to -722) | TTT TCC ATA CTT CAC TAG TCA AAA CCT | GAC AGT GAG AAG CTT TCG TAT TGA | 103 bp |
| *cki-1* | Promoter (-542 to -428) | TTC CTC ATA ATC ACG GAG CA | GGA ACC GAA GTG GTC AGA TG | 114 bp |
| *cki-1* | Promoter (-323 to -217) | TCT CCG ACT GCT GAC CT | CGA GAA GGG GTG GAG TCA TA | 106 bp |

**Treatment of mouse epithelial cell lines with human IL-17 and Nutlin3A**

A549 epithelial cells were seeded in a 6-well plate overnight and stimulated with 50-100ng/mL recombinant human-IL-17 (Peprotech, SKU 200-17) or 10uM Nutlin3A (Tocris, 675576-98-4), and supernatants and cell lysates were harvested 18 hours later.

**QUANTIFICATION AND STATISTICAL ANALYSIS**

Each ‘experiment’ refers to a set of independent biological samples that undergo treatment together. No statistical methods were used to predetermine sample size, and experiments were not randomized, but some were blinded. For the assays where we scored dauers, for each experiment, independent biological samples of the different strains were allowed to lay eggs and the eggs were subjected to 'treatment' or 'control' conditions. The results of an experiment generated counts categorized as either (a) dauers, or (b) non-dauers. The counts of dauers/non-dauers for the different types of experiments were used to generate a contingency table, and a chi-squared test was performed to test the independence (whether the results are related or not) of the experiment. This Chi-square statistic, degrees of freedom (df) and p value for each experiment is reported. Yates' continuity correction was used to prevent overestimation of statistical significance for small data. For qPCR or other experiments, the data were analyzed by using Student's t-test and/or one-way ANOVA with Tukey's correction (GraphPad Prism software) as described in respective figure legends. P values are indicated as follows: *$p<0.05$; **$p< 0.01$; ***$p<0.001$, ns, not significant.

#### DATA AVAILABILITY

The following data sets were generated

Genes differentially expressed in the following *C. elegans* larvae bleach-hatched and allowed to grow for 30-36 hr. at 25°C: wild type (N2), *ilc-17.1*(syb5296) X and praEx022 (*unc-54*p::*ilc-17.1* cDNA::3XFLAG::*tbb-2* 3'UTR; p*dat-1*::GFP::*unc-54* 3'UTR).

Genes deferentially expressed in the following *C. elegans* larvae bleach-hatched and allowed to grow for 15 hr. and 32 hr. at 25°C: wild type (N2), *ilc-17.1*(syb5296) X, *cep-1* (gk138) I, *ilc-17.1*(syb5296) X; *cep-1* (gk138) I, and VEP036 (unc-119 (ed4); gtIs1 [CEP-1::GFP + unc-119 (+)].

The data discussed in this publication have been deposited in NCBI's Gene Expression Omnibus (Edgar et al., 2002) and are accessible through GEO Series accession number GSE218596, , https://www.ncbi.nlm.nih.gov/geo/query/acc.cgi?acc=GSE218596 and GSE229132, https://www.ncbi.nlm.nih.gov/geo/query/acc.cgi?acc=GSE229132

### Figure S1

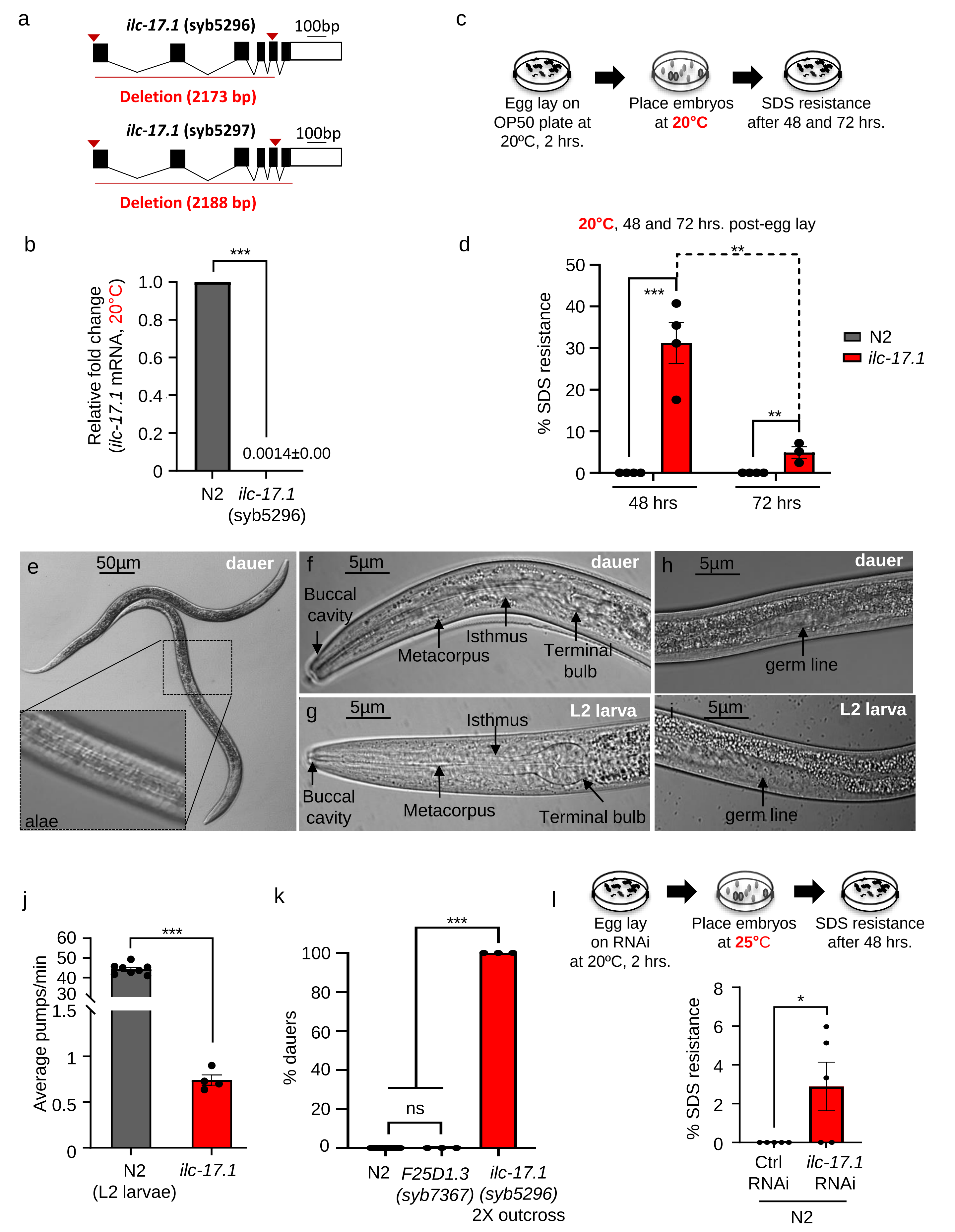

a
ilc-17.1 (syb5296)
100bp
Deletion (2173 bp)
ilc-17.1 (syb5297)
100bp
Deletion (2188 bp)
b
Relative fold change (ilc-17.1 mRNA, 20°C)
***
1.0
0.8
0.6
0.4
0.2
0
0.0014±0.00
N2
ilc-17.1 (syb5296)
c
Egg lay on OP50 plate at 20ºC, 2 hrs.
Place embryos at 20°C
SDS resistance after 48 and 72 hrs.
d
20°C, 48 and 72 hrs. post-egg lay
% SDS resistance
50
40
30
20
10
0
***
**
**
48 hrs
72 hrs
N2
ilc-17.1
e
50µm
dauer
alae
f
5µm
dauer
Buccal cavity
Metacorpus
Isthmus
Terminal bulb
g
5µm
L2 larva
Isthmus
Buccal cavity
Metacorpus
Terminal bulb
h
5µm
dauer
germ line
i
5µm
L2 larva
germ line
j
Average pumps/min
60
50
40
30
1.5
1
0.5
0
***
N2 (L2 larvae)
ilc-17.1
k
% dauers
100
80
60
40
20
0
***
ns
N2
F25D1.3 (syb7367)
ilc-17.1 (syb5296) 2X outcross
l
Egg lay on RNAi at 20°C, 2 hrs.
Place embryos at 25°C
SDS resistance after 48 hrs.
% SDS resistance
8
6
4
2
0
*
Ctrl RNAi
ilc-17.1 RNAi
N2

**Supplementary Figure S1: Loss of ILC-17.1 triggers *C. elegans* developmental diapause (dauer).**

**a.** Schematic of *ilc-17.1* gene depicting the *syb5296* and *syb5297* deletions made by CRISPR/Cas9 editing using two distinct guide RNAs. *ilc-17.1* (*syb5296*) X deletion mutants were used in all future experiments.

**b.** Average mRNA expression in *ilc-17.1* deletion mutants relative to wild type (N2). n=4-8 experiments. ***p < 0.001 (unpaired t-test,).

**c.** Schematic of experiment leveraging the resistance of dauers to 1% SDS used to identify dauers developing under optimal growth conditions at 20°C.

**d.** Percent *ilc-17.1* larvae that transiently enter the dauer state as seen by SDS-resistance 48 hrs and 72 hrs. post-hatching under optimal conditions at 20°C. ***p < 0.001, **p < 0.01 (n=4 experiments, Chi-squared = 166.53, df = 3, p-value < 2.2e-16).

**e-i.** Representative micrographs showing DIC images of **e**, whole *ilc-17.1* deletion mutant dauer larvae (inset shows alae), **f,** pharyngeal region and **h,** arrested germline 72 hr. post-hatching at 25°C. Wild-type (N2) larval stage 2 (L2) animals used as comparisons: **g**, pharyngeal region and **i**, germline. Scale bar, a=50µm, f-i, 5 µm.

**j**. Average pharyngeal pumping rates/minute in *ilc-17.1* deletion mutants relative to wild type (N2) larval stage 2 (L2) animals. n=4-8 experiments. ***p < 0.001, (unpaired t-test).

**k**. Percent F25D1.3 (syb7367) V, and 2X backcrossed *ilc-17.1* larvae that arrest as dauers at 25°C. ***p < 0.001, (n=3 experiments; 50-100 larvae each).

**l. Top:** Schematic of experiment. **Bottom:** Percent wild-type (N2) larvae that arrest as dauers as determined by SDS-resistance, 48 hrs post-hatching on control (Ctrl; L4440) and *ilc-17.1* RNAi at 25°C. *$p < 0.05$. (*n*=5 experiments, Chi-squared = 5.1712, df = 1, p-value = 0.02296).

Bars show the mean ± S.E.M**.** Individual points in the bar graphs in **d, k, l** represent the % dauers/experiment, Pearson's Chi-squared test with Yates' continuity correction.

Bars show mean ± S.E.M. Individual data points in the bar graphs showing % dauers represent the % dauers/experiment, and the bar graph depicts mean of these percentages.

Figure S2

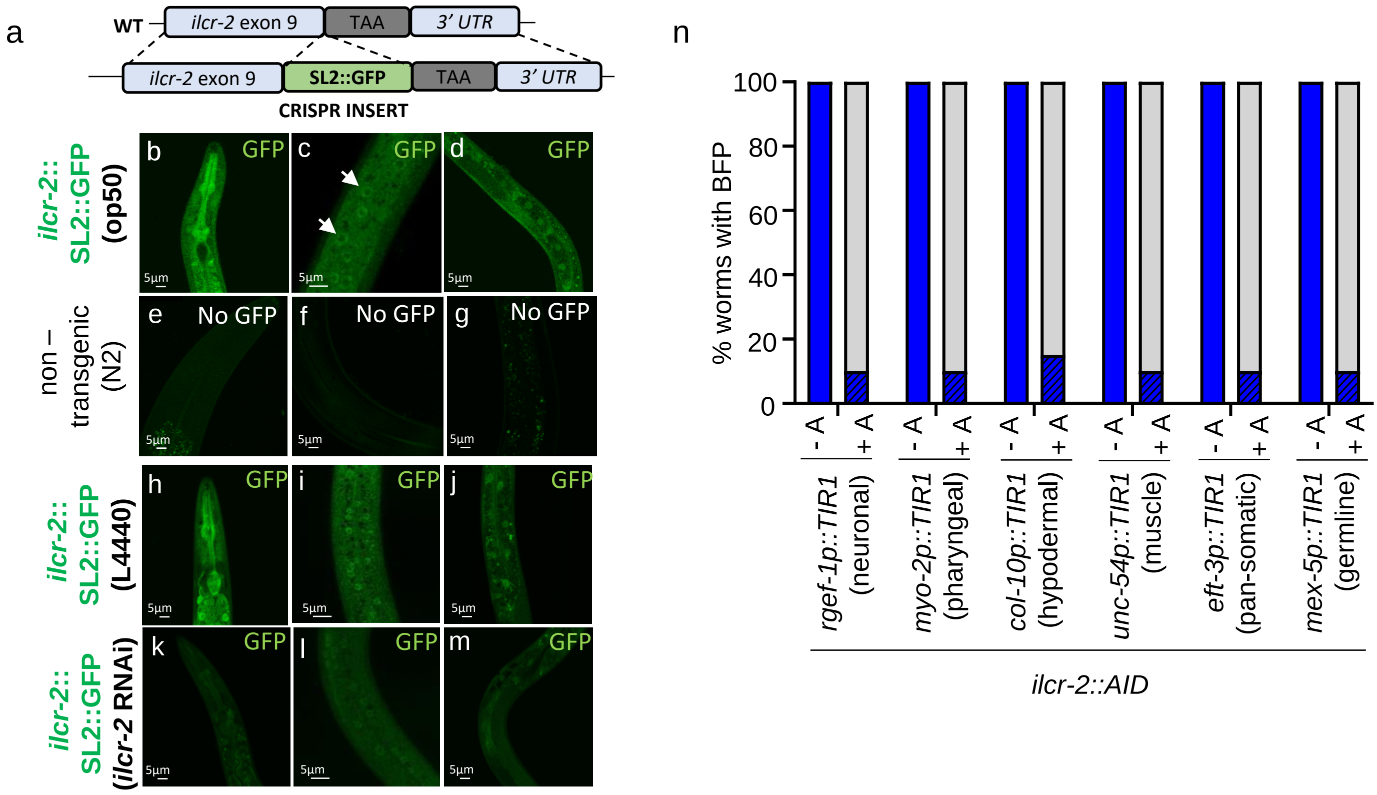
a
WT
ilcr-2 exon 9
TAA
3′ UTR
ilcr-2 exon 9
SL2::GFP
TAA
3′ UTR
CRISPR INSERT
ilcr-2::
SL2::GFP
(op50)
b
c
d
GFP
5µm
non –
transgenic
(N2)
e
f
g
No GFP
ilcr-2::
SL2::GFP
(L4440)
h
i
j
ilcr-2::
SL2::GFP
(ilcr-2 RNAi)
k
l
m
n
% worms with BFP
100
80
60
40
20
0
- A
+ A
rgef-1p::TIR1
(neuronal)
myo-2p::TIR1
(pharyngeal)
col-10p::TIR1
(hypodermal)
unc-54p::TIR1
(muscle)
eft-3p::TIR1
(pan-somatic)
mex-5p::TIR1
(germline)
ilcr-2::AID

**Supplementary Figure S2: Tissue-dependence of ILCR-2 knockdown in promoting dauer arrest.**

**a:** Schematic of CRISPR insertion of SL2::GFP into *ilcr-2* locus to identify sites of *ilcr-2* expression.

**b-d:** Representative micrographs showing GFP expression in cells of the pharynx (**b**), epidermis (**c**; arrows) and gonad (**d**).

**e-g:** Control, wild-type (N2) non-transgenic animals not expressing GFP confirm specificity of GFP expression. Scale bar=5µm.

**h-j:** Representative micrographs showing GFP expression in cells of the pharynx (**h**), epidermis i; arrows) and gonad (**j**) on control, L4440 RNAi.

**k-m**: Representative micrographs showing GFP expression in cells of the pharynx (**k**), epidermis (**l**; arrows) and gonad (**m**) on control, L4440 RNAi and *ilcr-2* RNAi.

**n.** Efficiency of degradation using the Auxin-induced degradation (AID) system to degrade ILCR-2 in different tissues of *C. elegans.* Animals expressing endogenous *ilcr-2* tagged with an AID degron using CRISPR/Cas9, were crossed into strains expressing an integrated bicistronic cassette of TIR1 and Blue Fluorescent Protein (BFP) (TIR1::F2A::mTagBFP2::AID*::NLS::tbb-2 3'UTR; Chr I). BFP was also tagged with an AID tag and served as an internal control for degradation upon auxin exposure. BFP fluorescence in the presence and absence of auxin was scored. Only larvae that showed <u>undetectable</u> BFP expression in their respective tissues of expression, were scored as positive for degradation. Y-axis: % larvae that expressed BFP. X-axis: genotype and treatment. TIR1 was expressed in: *rgef-1p* promoter: nervous system, *myo-*

*2p*: pharynx, *col-10p*: hypodermis, *unc-54p*: muscle, *eft-3p*: all somatic cells, and *mex-5p*: germline. -A: without auxin. +A: larvae on auxin. n=20 larvae/genotype/treatment.

Figure S3

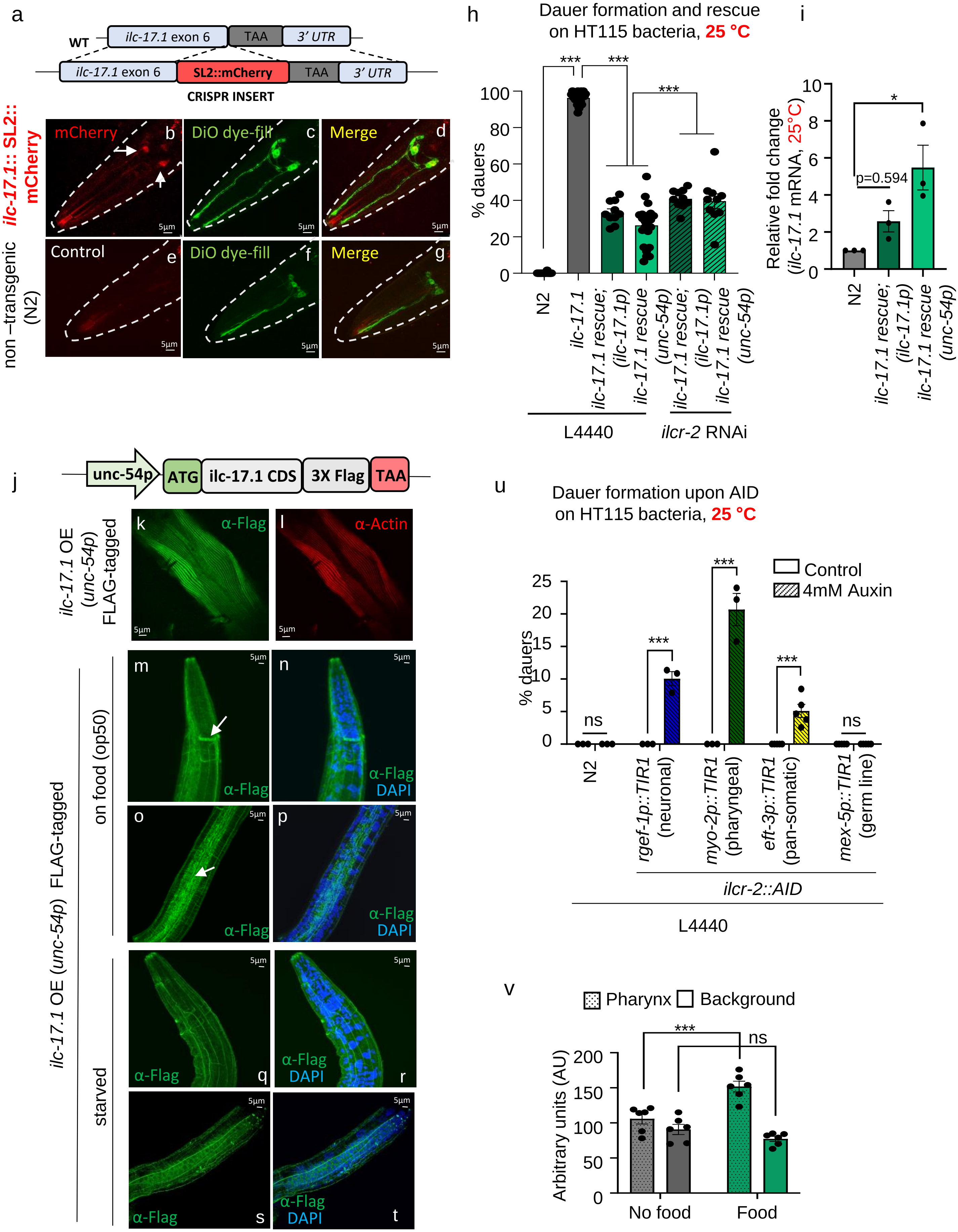
a
WT
ilc-17.1 exon 6
TAA
3' UTR
SL2::mCherry
CRISPR INSERT
ilc-17.1:: SL2:: mCherry
mCherry
DiO dye-fill
Merge
non –transgenic (N2)
Control
h
Dauer formation and rescue on HT115 bacteria, 25 °C
% dauers
N2
ilc-17.1
ilc-17.1 rescue; (ilc-17.1p)
ilc-17.1 rescue (unc-54p)
L4440
ilcr-2 RNAi
i
Relative fold change (ilc-17.1 mRNA, 25°C)
p=0.594
j
unc-54p
ATG
ilc-17.1 CDS
3X Flag
TAA
ilc-17.1 OE (unc-54p) FLAG-tagged
α-Flag
α-Actin
on food (op50)
starved
DAPI
u
Dauer formation upon AID on HT115 bacteria, 25 °C
Control
4mM Auxin
ns
rgef-1p::TIR1 (neuronal)
myo-2p::TIR1 (pharyngeal)
eft-3p::TIR1 (pan-somatic)
mex-5p::TIR1 (germ line)
ilcr-2::AID
v
Pharynx
Background
Arbitrary units (AU)
No food
Food

**Supplementary Figure S3: Characterization of ILC-17.1 signaling pathway components in *C. elegans*.**

**a.** Site of *ilc-17.1* mRNA expression**. a:** Schematic of CRISPR insertion of SL2::mCherry into *ilc-17.1* locus. **b-d**: Representative z-section micrographs showing mCherry expression in amphid neurons of *C. elegans* expressing mCherry as a bicistronic SL2 cassette along with the endogenous *ilc-17.1* gene (**b**), to report on sites of *ilc-17.1* mRNA expression. Amphid neurons were identified by DiO dye filling (**c**; green); mCherry overlapped with a subset of DiO filled neurons (**d**). **e-g**: Control, wild-type (N2) animals not expressing mCherry (**e**) confirm specificity of mCherry expression. Scale bar=5μm.

**h.** Percentage of wild type (N2) and *ilc-17.1* larvae that arrest as dauers on control HT115 (L4440) bacteria and *ilcr-2* RNAi. ***p < 0.001 (n=10-20 experiments, Chi-squared = 3206, df = 3, p-value < 2.2e-16). Representative average numbers of embryos/plates of wild type (N2), *ilc-17.1*, *ilc-17.1* rescued with *ilc-17.1p::ilc-17.1*, and *ilc-17.1* rescued with *unc-54p::ilc-17.1* on HT115 (L4440) bacteria are 59, 66, 66, 46 respectively. Individual points in the bar graphs represent the % dauers/experiment, Pearson's Chi-squared test with Yates' continuity correction.

**i.** Average expression in *ilc-17.1* mRNA in 32-26 hr. larvae of wild type (N2), *ilc-17.1* deletion mutants rescued with *ilc-17.1p::ilc-17.1::ilc-17.1 CDS* [*ilc-17.1* rescue; (*ilc-17.1p*)], and *unc-54p::ilc-17.1::ilc-17.1 CDS* [*ilc-17.1* rescue; (*unc-54p*)]. Values normalized to *pmp-3* and shown relative to wild type (N2) values. *p<0.05 (unpaired t-test).

**j.** Schematic of *ilc-17.1::*flag driven by the *unc-54* promoter [*ilc-17.1* OE *(unc-54p)*] that was used to rescue *ilc-17.1* deletion mutants. **k, l:** Representative micrographs showing a z-section of these animals immunostained with α-FLAG antibody, and α-actin (to show expression in body wall muscle cells). Scale bar=5μm

**m-p.** Representative micrographs showing z-sections through animals overexpressing ILC-17.1::FLAG in their body wall muscle cells as in **k**. ILC-17.1 protein was detected by immunolocalizing the associated FLAG tag. **m, n:** head regions (arrow: nerve cord). **o, p:** mid-body region (arrow: hypodermal cells).Note: ILC-17.1::FLAG is detected at the amphid commissures near the pharynx (**top panel**; arrow) and in the epidermis (**bottom panel**; arrows) in addition to the body wall muscle cells (**k, l**), Scale bar=5μm

**q-t:** Representative micrographs showing z-sections through animals overexpressing ILC-17.1::FLAG in their body wall muscle cells as in **m-p, but in the absence of food.** ILC-17.1 protein continued to be detected outside the body wall muscle cells where the mRNA was expressed (**r, t**), Scale bar=5μm

**u.** Percentage dauers following tissue specific degradation of ILCR-2. X-axis: tissue specific TIR1 expression, (n=3-4 experiments; Chi-squared = 215.63, df =7 , p-value <2.2e-16). **Note:** the experiment was conducted on RNAi plates seeded with **HT115 (L4440)** experiments and supplemented with auxin (Compare with **Figure 1f, on OP50**).

**v.** Quantification of fluorescent intensity in pharynx versus background (arbitrary units; AU), in larvae harboring CRISPR tagged endogenous ILC-17.1::HA**,** immunostained with anti-HA (**Figure 1 j, l, n)**. n=3 experiments. 5-6 larvae/experiment. ***p < 0.001, (unpaired t-test).

Bars show the mean ± S.E.M.

Figure S4

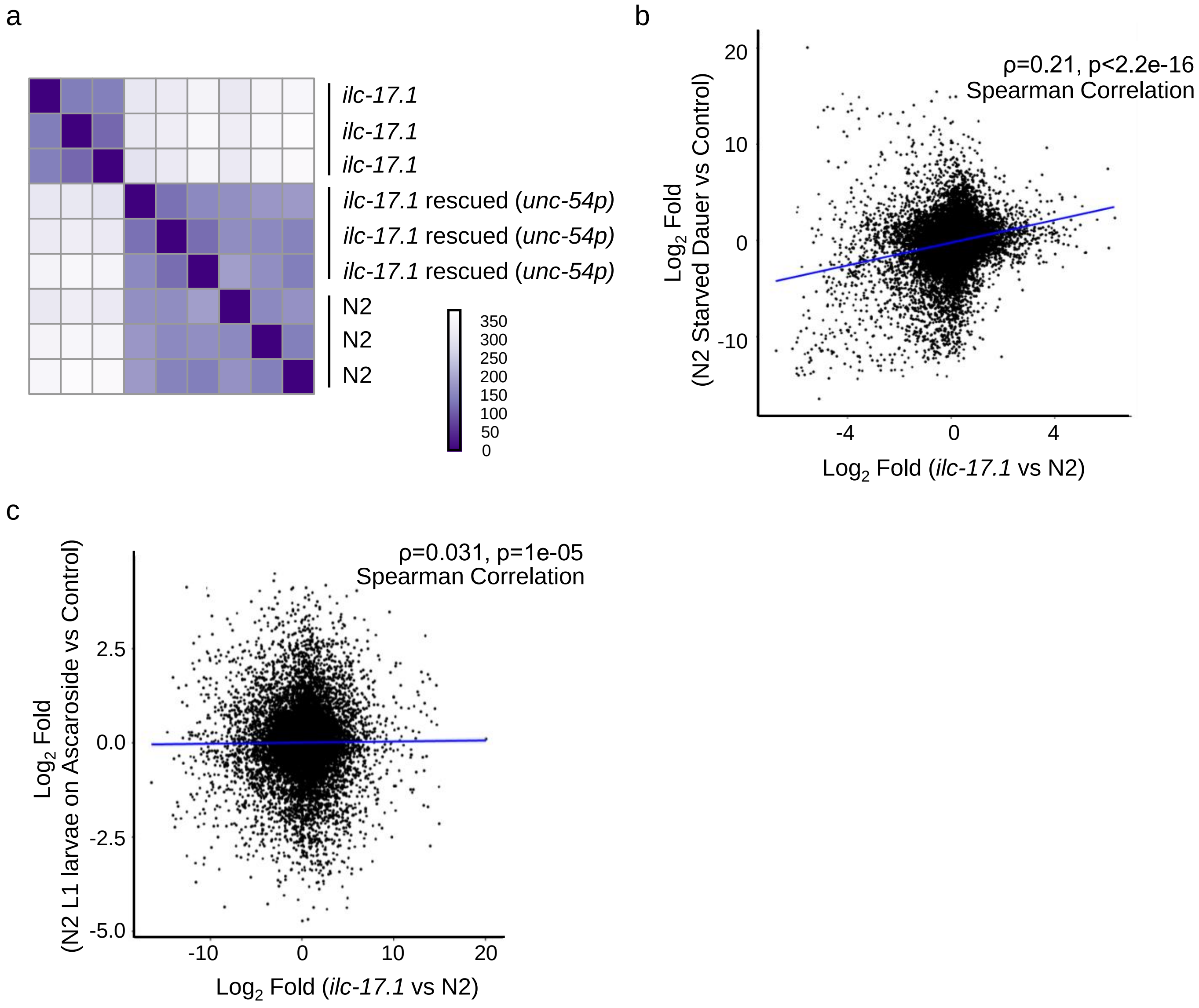
a
ilc-17.1
ilc-17.1
ilc-17.1
ilc-17.1 rescued (unc-54p)
ilc-17.1 rescued (unc-54p)
ilc-17.1 rescued (unc-54p)
N2
N2
N2
350
300
250
200
150
100
50
0
b
ρ=0.21, p<2.2e-16
Spearman Correlation
Log2 Fold
(N2 Starved Dauer vs Control)
20
10
0
-10
-4
0
4
Log2 Fold (ilc-17.1 vs N2)
c
ρ=0.031, p=1e-05
Spearman Correlation
Log2 Fold
(N2 L1 larvae on Ascaroside vs Control)
2.5
0.0
-2.5
-5.0
-10
0
10
20
Log2 Fold (ilc-17.1 vs N2)

**Supplementary Figure S4: RNA-seq analysis of ILC-17.1 deficient larvae.**

**a.** Pair-wise distance matrix of RNA-seq samples shows the expected clustering of total RNA of the biological triplicates of each strain [Strains used: wild type (N2), *ilc-17.1*, and *ilc-17.1* rescued with *unc-54p::ilc-17.1* CDS]

**b.** Spearman correlation between all genes in a microarray analysis of dauer larvae that were induced by starvation and *ilc-17.1* deleted larvae *en route* to dauer arrest grown for 32-36 hrs. at 25°C.

**c**. Spearman correlation between all genes in L1 larvae exposed to ascarosides and *ilc-17.1* deleted larvae *en route* to dauer arrest grown for 32-36 hrs. at 25°C.

Figure S5

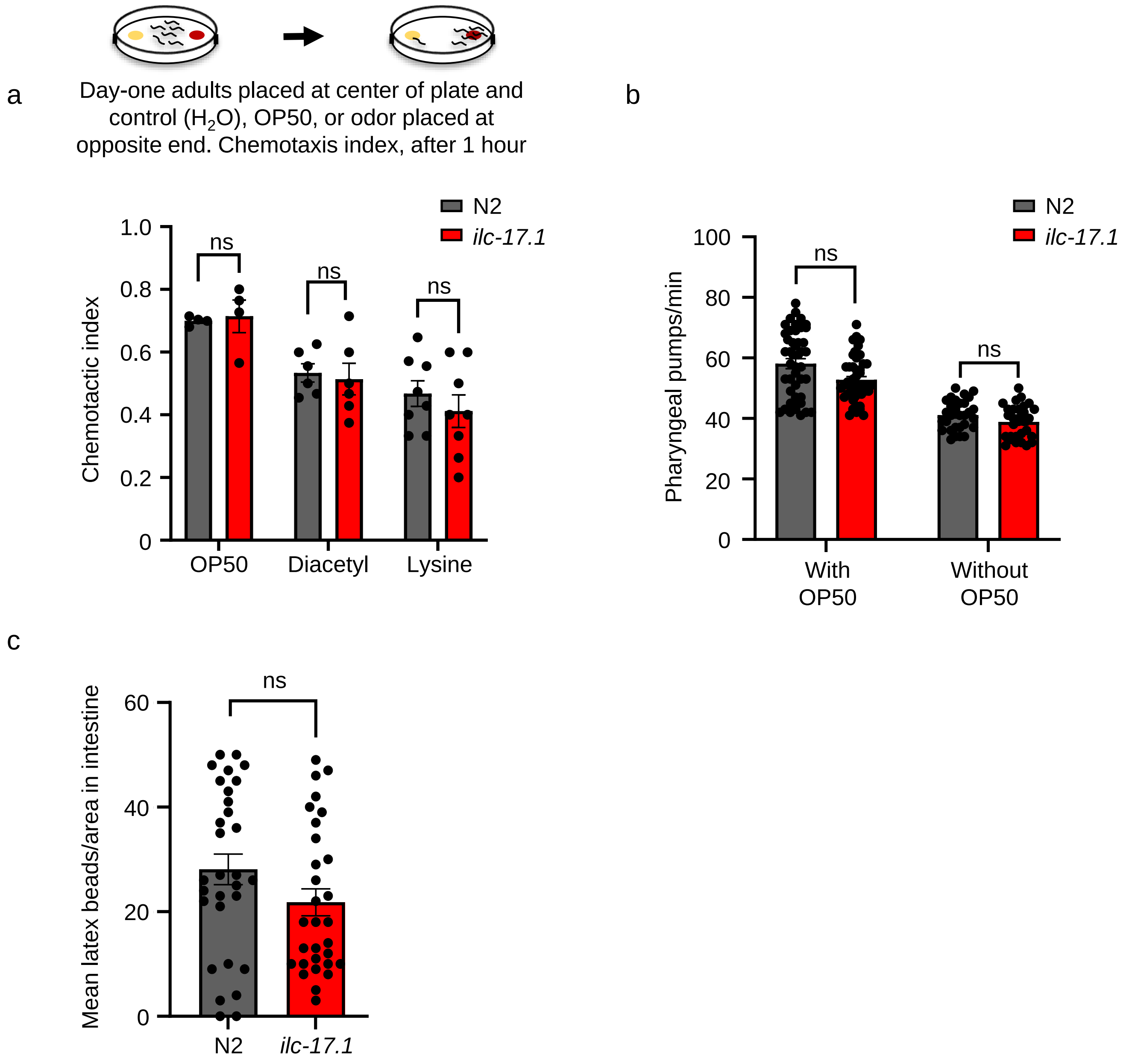
a
Day-one adults placed at center of plate and control ($H_2O$), OP50, or odor placed at opposite end. Chemotaxis index, after 1 hour
N2
ilc-17.1
Chemotactic index
1.0
0.8
0.6
0.4
0.2
0
ns
ns
ns
OP50
Diacetyl
Lysine
b
N2
ilc-17.1
Pharyngeal pumps/min
100
80
60
40
20
0
ns
ns
With OP50
Without OP50
c
Mean latex beads/area in intestine
60
40
20
0
ns
N2
ilc-17.1

**Supplementary Figure S5 (associated with Extended text 1): ILC-17.1 loss does not affect the animal's chemotaxis or pumping.**

**Extended text 1.** *ilc-17.1* deleted animals showed normal chemotaxis towards lawns of OP50, or organic molecules like lysine or diacetyl thought to serve as bacterial signals [(42) Supplementary Fig. S5a]. They were able to feed like wild-type animals as determined pharyngeal pumping rates on or off food (Supplementary Fig. S5b), and the accumulation of latex beads in their intestinal lumen used to track ingestion rates (Supplementary Fig. S5c). These data suggested that *ilc-17.1* deficient larvae arrested development and entered a dauer diapause state despite being able to find and ingest nutrients.

**a. Top:** Schematic of experimental design. **Bottom:** Average chemotaxis index towards OP50, 0.01% v/v diacetyl, and 1M lysine for *ilc-17.1* deletion mutants and wild-type (N2) day-one adults at 20°C. n= 5-8 experiments. ns, not significant (unpaired t-test). Data points: average chemotaxis/experiment.

**b.** Number of pharyngeal pumps/min in *ilc-17.1* deletion mutants and wild-type (N2) day-one adults at 20°C, on OP50 lawns, and on plates without food. n=3 experiments, with at least 5 animals/experiment scored. Data points: pumping rates of individual animals. ns, not significant (unpaired t-test).

**c.** Average numbers of fluorescent latex beads in similar areas of distal gut lumen of 40-hour *ilc-17.1* deletion mutants and wild-type (N2) larvae at 20°C. n=3 experiments, with at least 10 animals/experiment. Data points: the numbers of beads in individual animals. ns, not significant (unpaired t-test).

Data in all graphs show mean ± S.E.M.

Figure S6

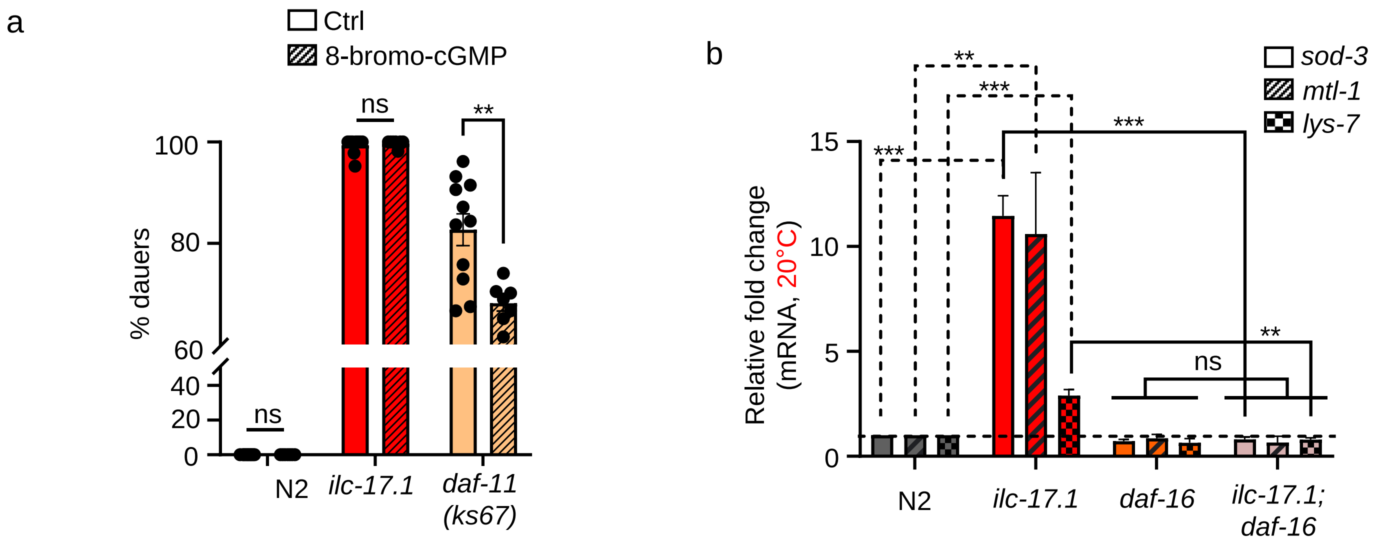
a
Ctrl
8-bromo-cGMP
% dauers
100
80
60
40
20
0
ns
ns
**
N2
ilc-17.1
daf-11 (ks67)
b
sod-3
mtl-1
lys-7
Relative fold change (mRNA, 20°C)
15
10
5
0
**
***
***
***
**
ns
N2
ilc-17.1
daf-16
ilc-17.1; daf-16

**Supplementary Figure S6: ILC-17.1 loss-induced dauer arrest cannot be rescued by 8-bromo-cGMP, and activated DAF-16 even at 20°C.**

**a.** Percent dauers amongst wild type (N2), *ilc-17.1* deletion mutants and *daf-11(ks67)* raised on Control ($H_2O$) and 5mM 8-bromo-cGMP at 25°C. Note discontinuous Y-axis. ***$p < 0.001$, Pearson's Chi-squared test with Yates correction, (n=3 experiments. Chi squared = 798, df = 3, p-value <2e-16.). Individual points in the bar graphs represent the % dauers/experiment. n=7-11 experiments, Chi squared = 1949, df = 5, p-value < 2.2e-16.

**b.** Average *sod-3, mtl-1, and lys-7* mRNA levels in 40 hr. old larvae grown at 20°C. mRNA levels were determined relative to *pmp-3* and normalized to wild type (N2) values. n=3 experiments. ***$p < 0.001$, **$p < 0.01$, ns, not significant (unpaired t-test).

Data in all graphs show mean ± S.E.M.

Figure S7

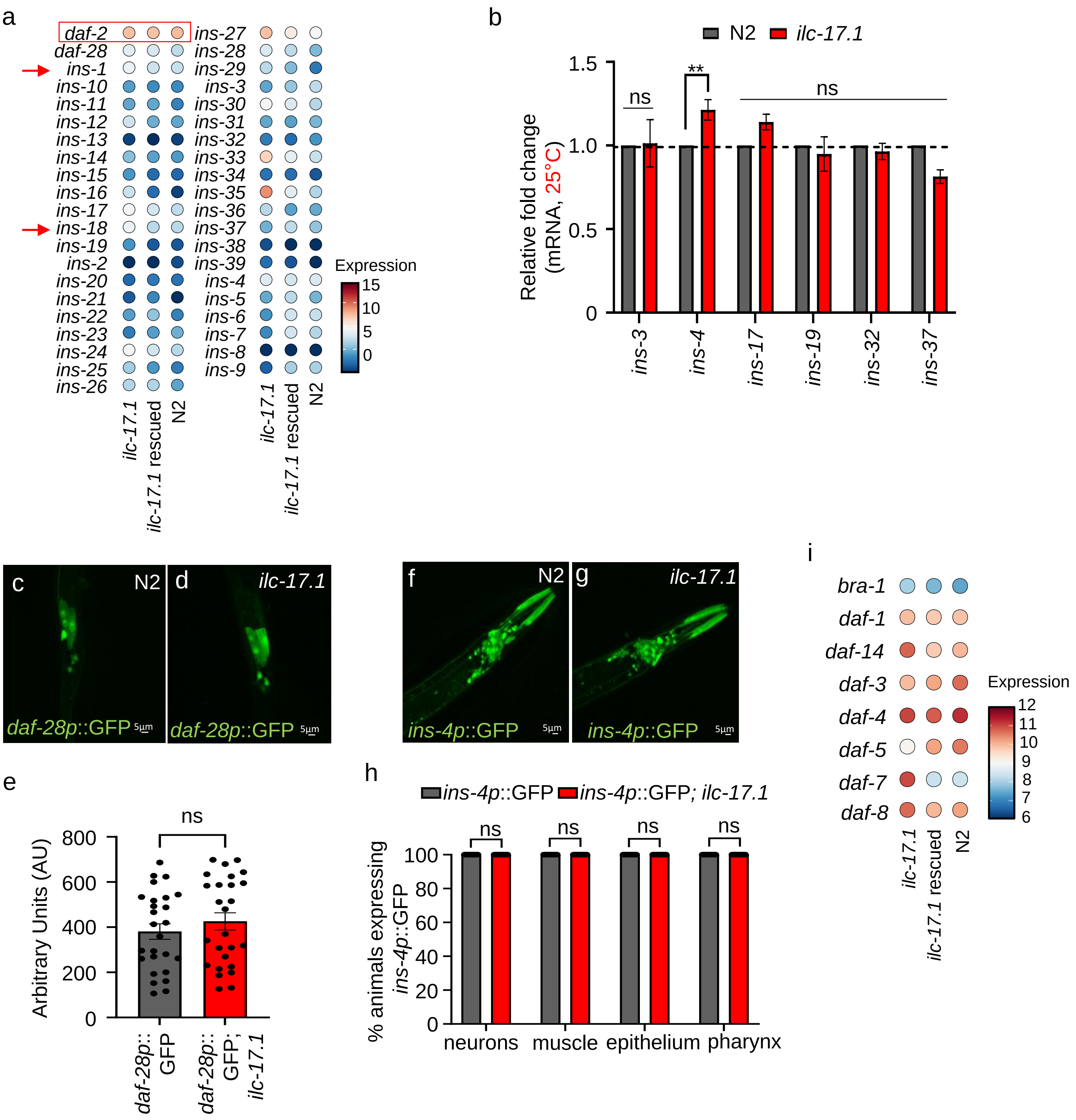
a
daf-2
daf-28
ins-1
ins-10
ins-11
ins-12
ins-13
ins-14
ins-15
ins-16
ins-17
ins-18
ins-19
ins-2
ins-20
ins-21
ins-22
ins-23
ins-24
ins-25
ins-26
ins-27
ins-28
ins-29
ins-3
ins-30
ins-31
ins-32
ins-33
ins-34
ins-35
ins-36
ins-37
ins-38
ins-39
ins-4
ins-5
ins-6
ins-7
ins-8
ins-9
ilc-17.1
ilc-17.1 rescued
N2
Expression
15
10
5
0
b
N2
ilc-17.1
1.5
1.0
0.5
0
Relative fold change
(mRNA, 25°C)
ns
**
ns
ins-3
ins-4
ins-17
ins-19
ins-32
ins-37
c
N2
daf-28p::GFP
5µm
d
ilc-17.1
daf-28p::GFP
5µm
f
N2
ins-4p::GFP
5µm
g
ilc-17.1
ins-4p::GFP
5µm
i
bra-1
daf-1
daf-14
daf-3
daf-4
daf-5
daf-7
daf-8
Expression
12
11
10
9
8
7
6
ilc-17.1
ilc-17.1 rescued
N2
e
ns
800
600
400
200
0
Arbitrary Units (AU)
daf-28p:: GFP
daf-28p:: GFP; ilc-17.1
h
ins-4p::GFP
ins-4p::GFP; ilc-17.1
ns
ns
ns
ns
100
80
60
40
20
0
% animals expressing ins-4p::GFP
neurons
muscle
epithelium
pharynx

**Supplementary Figure S7 (associated with Extended text 2): ILC-17.1-deficiency does not cause obvious changes in the expression of Insulin-signaling pathway (ILS), and TGFβ signaling pathway genes.**

**Extended text 2**: The expression levels of *daf-2* mRNA and the 28 insulin ligands(50, 51) were not decreased in *ilc-17.1* deletion mutants, (Supplementary Fig. S7a, *daf-2* boxed; Supplementary Fig. S7b; Supplementary Table 7). The localization of insulin expression (51) also did not appear to be changed (Supplementary Fig. S7c-h). The expression levels of *daf-7* mRNA and other genes in the TGFβ signaling pathway (50, 51) were also not decreased in *ilc-17.1* deletion mutants, (Supplementary Fig. S7i).

**a.** Heatmap depicting expression levels ($log_2$ normalized counts) of *daf-2* (boxed) and insulin ligands in the *ilc-17.1* deletion mutants, *ilc-17.1* deletion mutants rescued by *unc-54p::ilc-17.1* and wild type (N2). Arrows highlight lack of changes in *ins-1* and *ins-18* mRNA. RNA-seq data from larvae grown for 32-36 hrs. at 25°C.

**b.** Average mRNA levels (insulins) in 32-36 hr. larvae grown at 25°C. mRNA levels were determined relative to *pmp-3* and normalized to wild type (N2) values. n=3 experiments. **$p < 0.001$, ns, not significant (unpaired t-test).

**c-d**: Representative micrographs showing *daf-28p*::GFP expression in the last intestinal cell of 32-36hr. larvae at 25°C of *ilc-17.1* deletion mutants and wild type (N2). Images are projections of confocal z-sections. Scale bar: 5μm. **e** Quantification of fluorescent intensity (arbitrary units, AU) of GFP. n=3 experiments. Data points: the fluorescence intensity of *daf-28p*::GFP in individual animals. ns, not significant (unpaired t-test).

**f-g** Representative micrographs showing *ins-4p*::GFP expression in the pharynx and neurons of 32-36 hr. larvae at 25°C in *ilc-17.1* deletion mutants and wild type (N2). Images are projections of confocal z-sections. Scale bar: 5μm. **h:** Bars represent percent larvae that express GFP in the different tissues where INS-4 is known to be expressed. n=3 experiments of 5-10 larvae of each strain. ns, not significant (unpaired t-test).

**i.** Heatmap depicting expression levels ($log_2$ normalized counts) of TGFβ signaling pathway components in the *ilc-17.1* deletion mutants, *ilc-17.1* deletion mutants rescued by *unc-54p::ilc-17.1*, and wild type (N2).

Data in all graphs show mean ± S.E.M.

Figure S8

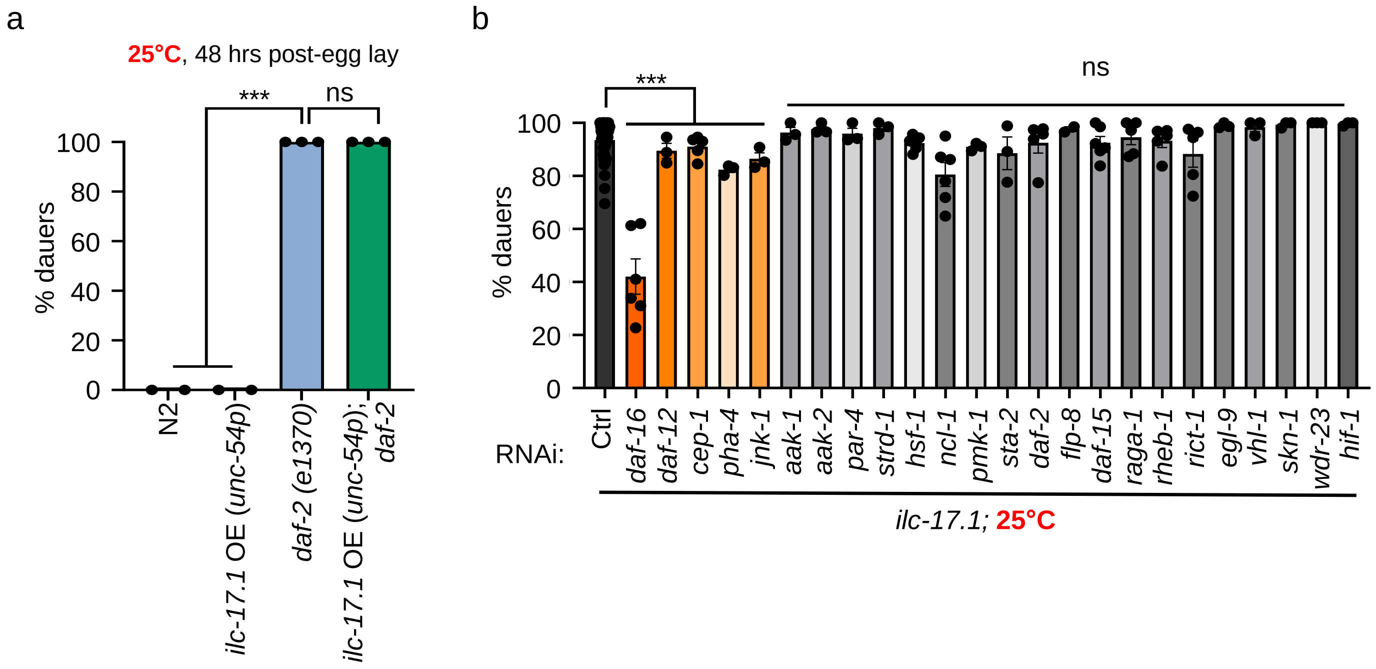

a
25°C, 48 hrs post-egg lay
***
ns
% dauers
0
20
40
60
80
100
N2
ilc-17.1 OE (unc-54p)
daf-2 (e1370)
ilc-17.1 OE (unc-54p); daf-2
b
***
ns
% dauers
RNAi:
Ctrl
daf-16
daf-12
cep-1
pha-4
jnk-1
aak-1
aak-2
par-4
strd-1
hsf-1
ncl-1
pmk-1
sta-2
daf-2
flp-8
daf-15
raga-1
rheb-1
rict-1
egl-9
vhl-1
skn-1
wdr-23
hif-1
ilc-17.1; 25°C

**Supplementary Figure S8: Investigating the genetic dependence of dauer arrest by loss of ILC-17.1.**

**a.** Overexpressing ILC-17.1 under *unc-54p* [*ilc-17.1* OE (*unc-54p)*] in *daf-2* (e1370) III strains does not rescue their dauer arrest. ***p < 0.001, ns, not significant (n=3 experiments, Chi-squared = 521, df = 3, p-value < 2.2e-16).

**b.** Percent dauers in *ilc-17.1* deletion mutant larvae following RNAi-mediated downregulation of several genes known to interact with DAF-16 and DAF3/DAF-5. *daf-16* RNAi was used as a positive control. ***p < 0.001, ns, not significant. (*n=3*-5 experiments, Chi-squared = 156.66, df = 3, p-value < 2.2e-16).

Data in all graphs show mean ± S.E.M. Individual points in the bar graphs represent the % dauers/experiment, and the bar graph depicts mean of these percentages.

Pearson's Chi-squared test with Yates' continuity correction.

Figure S9

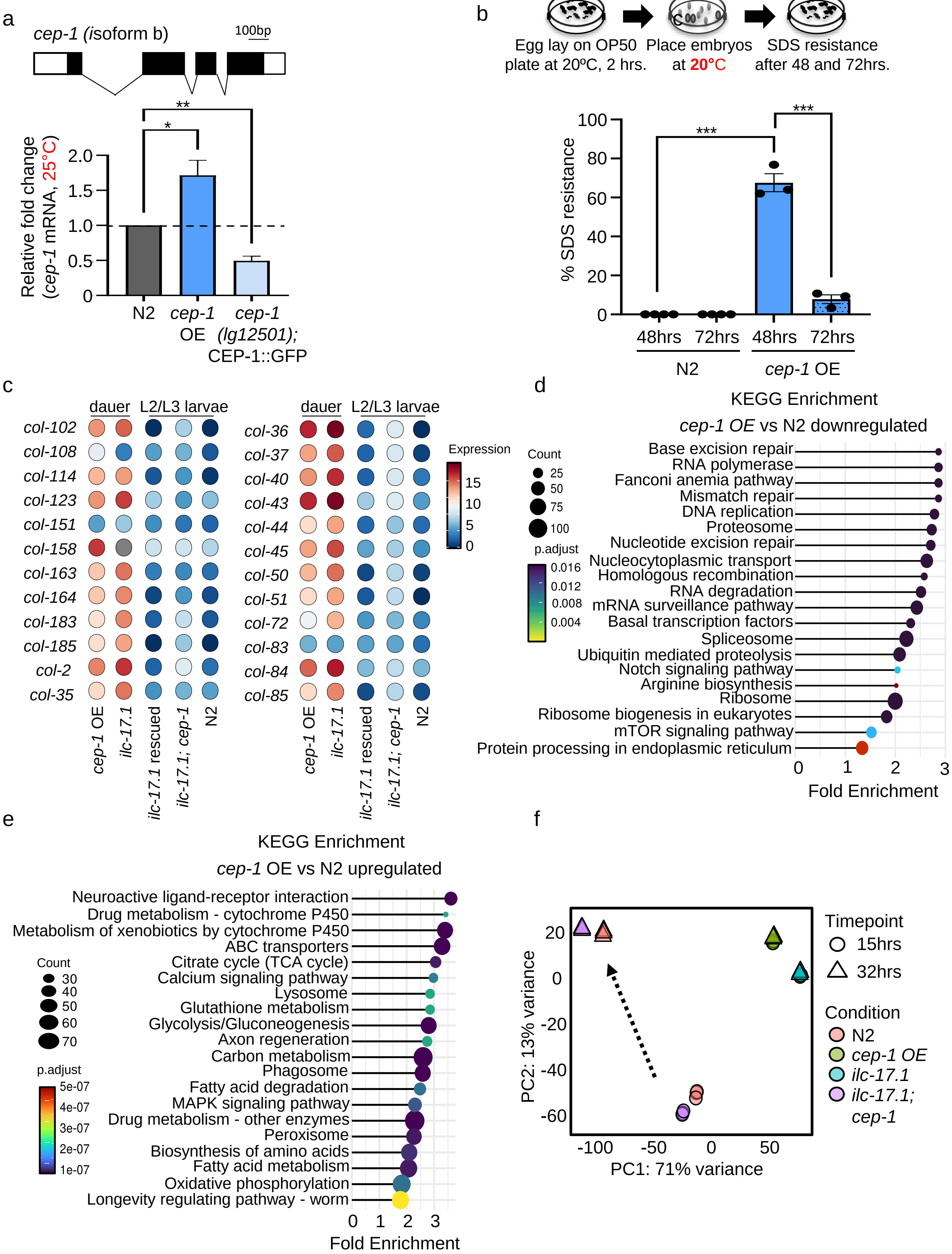
a
cep-1 (isoform b)
100bp
Relative fold change (cep-1 mRNA, 25°C)
N2
cep-1 OE
cep-1 (lg12501); CEP-1::GFP
b
Egg lay on OP50 plate at 20ºC, 2 hrs.
Place embryos at 20°C
SDS resistance after 48 and 72hrs.
% SDS resistance
48hrs
72hrs
N2
cep-1 OE
c
dauer
L2/L3 larvae
col-102
col-108
col-114
col-123
col-151
col-158
col-163
col-164
col-183
col-185
col-2
col-35
col-36
col-37
col-40
col-43
col-44
col-45
col-50
col-51
col-72
col-83
col-84
col-85
cep-1 OE
ilc-17.1
ilc-17.1 rescued
ilc-17.1; cep-1
N2
Expression
d
KEGG Enrichment
cep-1 OE vs N2 downregulated
Count
p.adjust
Base excision repair
RNA polymerase
Fanconi anemia pathway
Mismatch repair
DNA replication
Proteosome
Nucleotide excision repair
Nucleocytoplasmic transport
Homologous recombination
RNA degradation
mRNA surveillance pathway
Basal transcription factors
Spliceosome
Ubiquitin mediated proteolysis
Notch signaling pathway
Arginine biosynthesis
Ribosome
Ribosome biogenesis in eukaryotes
mTOR signaling pathway
Protein processing in endoplasmic reticulum
Fold Enrichment
e
KEGG Enrichment
cep-1 OE vs N2 upregulated
Neuroactive ligand-receptor interaction
Drug metabolism - cytochrome P450
Metabolism of xenobiotics by cytochrome P450
ABC transporters
Citrate cycle (TCA cycle)
Calcium signaling pathway
Lysosome
Glutathione metabolism
Glycolysis/Gluconeogenesis
Axon regeneration
Carbon metabolism
Phagosome
Fatty acid degradation
MAPK signaling pathway
Drug metabolism - other enzymes
Peroxisome
Biosynthesis of amino acids
Fatty acid metabolism
Oxidative phosphorylation
Longevity regulating pathway - worm
Count
p.adjust
Fold Enrichment
f
Timepoint
15hrs
32hrs
Condition
N2
cep-1 OE
ilc-17.1
ilc-17.1; cep-1
PC1: 71% variance
PC2: 13% variance

**Supplementary Figure S9: CEP-1/p53 overexpression induces dauer arrest.**

**a. Top:** Schematic of *cep-1* mRNA, isoform b. **Bottom:** Average *cep-1* isoform b mRNA levels in 32-36 hr. larvae grown at 25°C. mRNA levels were determined relative to *pmp-3* and normalized to wild type (N2) values. *n*=3 experiments. **p < 0.001, *p < 0.05 (unpaired t-test).

**b. Top:** Schematic of SDS treatment to assess dauer formation at 20°C. **Bottom:** Percent of wild type (N2) and *cep-1* overexpressing larvae (*cep-1* OE) that enter a dauer state during development, 48 hrs. and 72 hrs. post-hatching at 20°C. ***p < 0.001, (n=3 experiments, Chi-squared = 414.65, df = 3, p-value < 2.2e-16). Individual points in the bar graphs represent the % dauers/experiment, Pearson's Chi-squared test with Yates' continuity correction.

**c.** Heatmap comparing expression levels ($\log_2$ normalized counts) of the major dauer-specific collagens in the *ilc-17.1* deletion mutants, CEP-1/p53 overexpressing larvae, *ilc-17.1; cep-1* double mutants, *ilc-17.1* deletion mutants rescued by *unc-54p::ilc-17.1,* and wild type (N2).

**d.** KEGG enrichments associated with differentially downregulated genes (p.adjust <0.05) in CEP-1/p53 overexpressing larvae [RNA-seq: 32-36 hr. larvae grown at 25°C; **Supplementary Tables 9**].

**e**. KEGG enrichments associated with differentially upregulated genes (padjust <0.05) in CEP-1/p53 overexpressing larvae [RNA-seq: 32-36 hr. larvae grown at 25°C; **Supplementary Tables 9**].

**f.** Principle Component Analysis **(PCA)** of the observed variance in the triplicate RNA-seq samples collected during development, 15 hrs. and 32 hrs. post-hatching at 25°C, of *ilc-17.1* deletion mutants, CEP-1/p53 overexpressing larvae, *ilc-17.1; cep-1* double mutants and wild-type (N2) larvae (**Supplementary Tables 9).** Arrow highlights distance between 15 hrs and 32 hrs post-hatched N2 larvae and *ilc-17.1; cep-1* larvae as they continue to develop. Note that there is little change between 15 hrs and 32 hrs in *ilc-17.1* and *cep-1 OE* larvae.

Data in all graphs show mean ± S.E.M. Individual points in the bar graphs in **b**, represent the % dauers/experiment, and the bar graph depicts mean of these percentages.

Figure S10

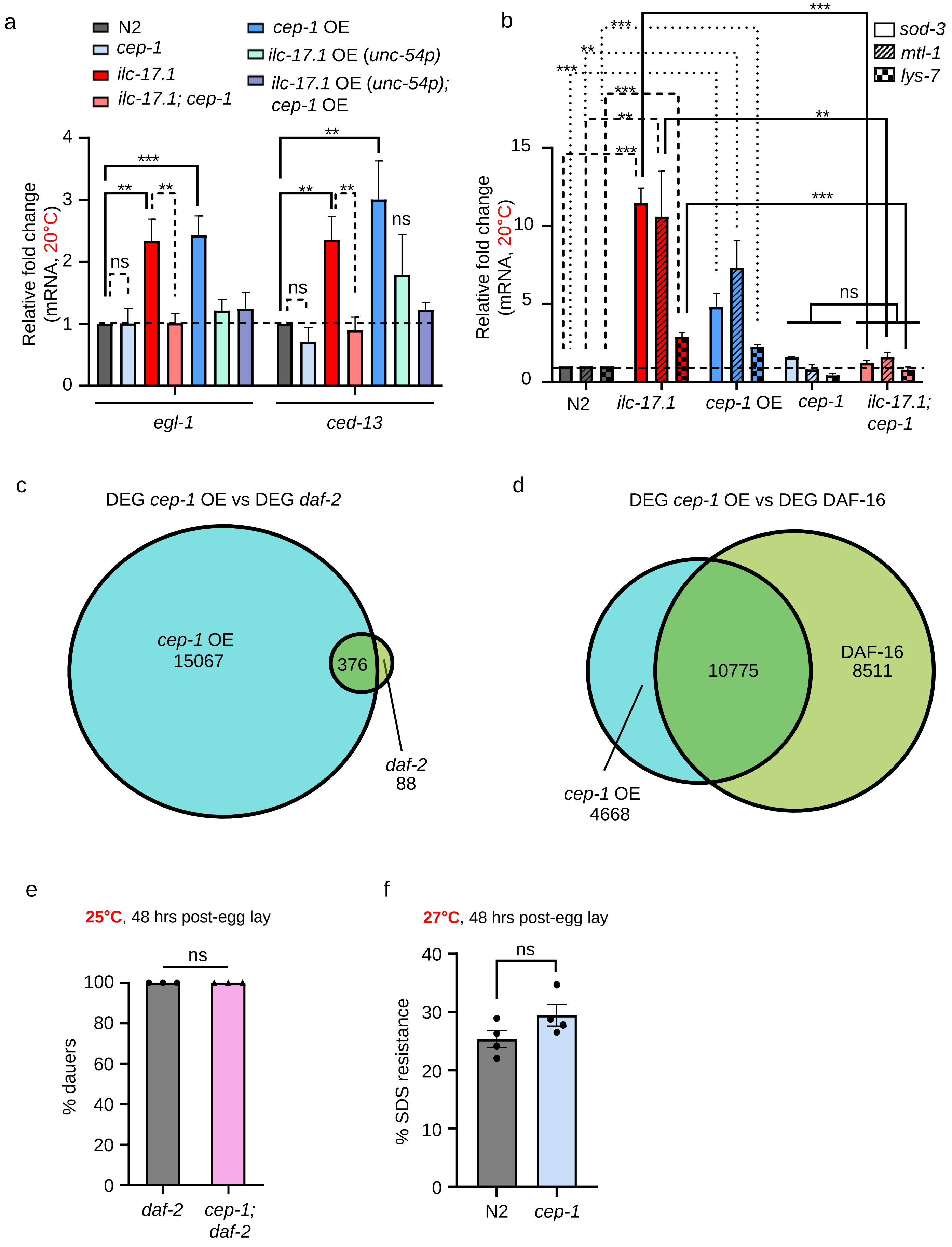

a
N2
cep-1
ilc-17.1
ilc-17.1; cep-1
cep-1 OE
ilc-17.1 OE (unc-54p)
ilc-17.1 OE (unc-54p); cep-1 OE
Relative fold change (mRNA, 20°C)
0
1
2
3
4
***
**
**
ns
**
**
**
ns
ns
egl-1
ced-13
b
sod-3
mtl-1
lys-7
Relative fold change (mRNA, 20°C)
0
5
10
15
***
***
**
***
***
**
***
**
***
ns
N2
ilc-17.1
cep-1 OE
cep-1
ilc-17.1; cep-1
c
DEG cep-1 OE vs DEG daf-2
cep-1 OE
15067
376
daf-2
88
d
DEG cep-1 OE vs DEG DAF-16
10775
DAF-16
8511
cep-1 OE
4668
e
25°C, 48 hrs post-egg lay
% dauers
0
20
40
60
80
100
ns
daf-2
cep-1; daf-2
f
27°C, 48 hrs post-egg lay
% SDS resistance
0
10
20
30
40
ns
N2
cep-1

**Supplementary Figure S10: CEP-1/p53 overexpression upregulates DAF-16/FOXO targets but *cep-1* is not downstream of *daf-2.***

**a.** Average *egl-1* and *ced-13* mRNA levels in 40 hr. larvae grown at 20°C. mRNA levels were determined relative to *pmp-3* and normalized to wild type (N2) values. n=6-7 experiments. ***p < 0.001, **p < 0.01, ns, not significant (unpaired t-test).

**b.** Average *sod-3, mtl-1, and lys-7* mRNA levels in 40 hr. larvae grown at 20°C. mRNA levels were determined relative to *pmp-3* and normalized to wild type (N2) values. Strains used are on X-axis. n=6-10 experiments. ***p < 0.001, **p < 0.01, ns, not significant. (unpaired t-test).

**c.** Venn diagram depicting overlap between differentially expressed genes (p<0.05) in CEP-1/p53 overexpressing larvae, 32-36 hr. post-hatching at 25°C. and published *daf-2*-dependent genes (*Murphy, C.T., et al., 2003, Nature, 424, 277-284*). p=0.0; hypergeometric test.

**d.** Venn diagram depicting overlap between differentially expressed genes (p<0.05) in CEP-1/p53 overexpressing larvae, 32-36 hr. post-hatching at 25°C and published DAF-16 targets (*Tepper, R.G., et al., 2013, Cell 154, 676-690*). p=0.0; hypergeometric test.

**e.** Percent dauers in *daf-2* and *cep-1;daf-2* double mutants. Note: no rescue. n=3 experiments. ns, not significant. (Chi-squared = Not appropriate, df = 1, p-value = no significant)

**f.** Percent high temperature (27.5°C; HID phenotype) dauers in wild type (N2) and *cep-1* deletion mutants. ns, not significant. (n=4 experiments, Chi-squared = 0.67295, df = 1, p-value = 0.412).

Data in all graphs show mean ± S.E.M. Individual points in the bar graphs in **e,f** represent the % dauers/experiment, Pearson's Chi-squared test with Yates' continuity correction.

Figure S11

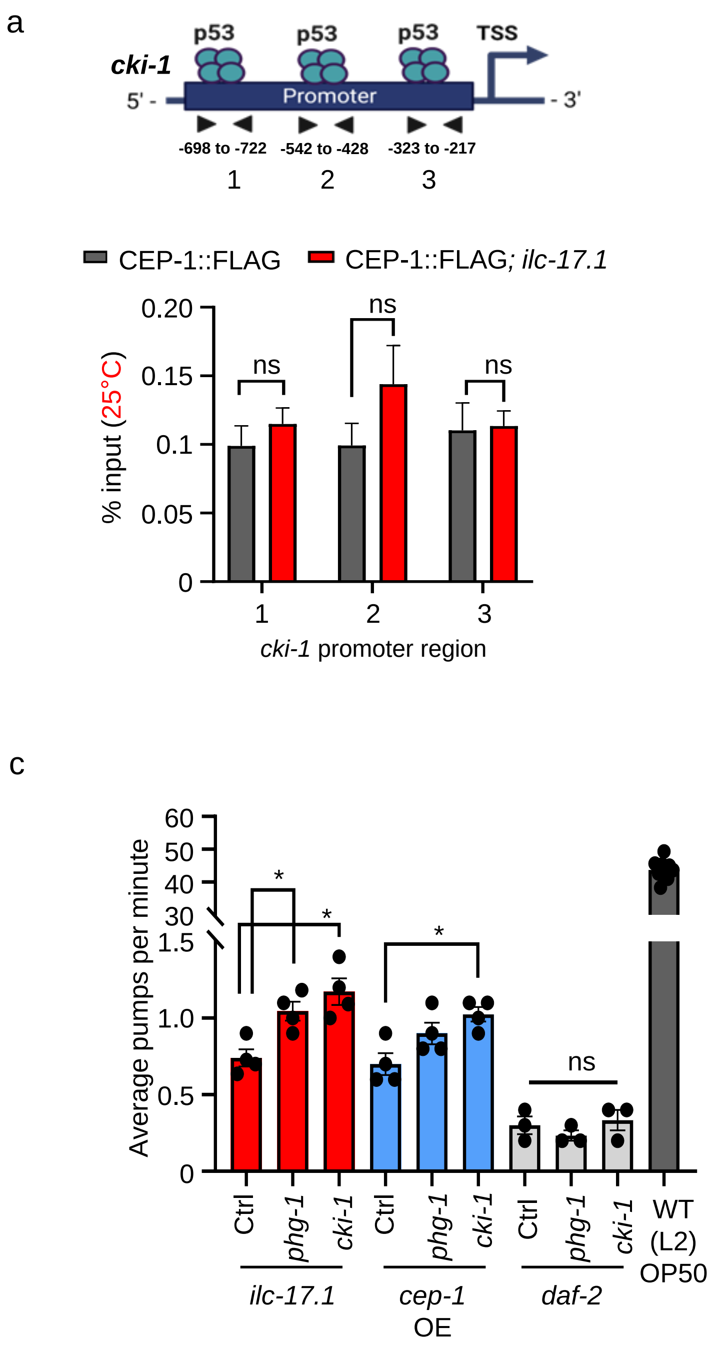
a
p53
p53
p53
TSS
cki-1
5' -
Promoter
- 3'
-698 to -722
-542 to -428
-323 to -217
1
2
3
CEP-1::FLAG
CEP-1::FLAG; ilc-17.1
% input (25°C)
0.20
0.15
0.1
0.05
0
ns
ns
ns
cki-1 promoter region
b
25°C, 48 hrs post-egg lay
% dauers
100
95
90
85
80
20
0
***
**
***
**
ns
Ctrl
phg-1
cki-1
N2
ilc-17.1
cep-1 OE
daf-2
c
Average pumps per minute
60
50
40
30
1.5
1.0
0.5
0
*
*
*
ns
WT (L2) OP50

**Supplementary Figure S11: CEP-1/p53 overexpression upregulates cell cycle inhibitors**

**a.** CEP-1 occupancy (expressed as percent input) in 32-36 hr. larvae grown at 25°C, measured at three promoter proximal regions of *cki-1* (Schematic on **top**). Strains used: CEP-1::FLAG expressing animals in wild-type background and in *ilc-17.1* deletion background. n=4 experiments. ns, not significant. (unpaired t-test).

**b.** Percent dauers in *ilc-17.1* larvae, CEP-1/p53 overexpressing larvae and *daf-2* larvae upon RNAi induced downregulation of *cki-1* and *phg-1.* Note discontinuous Y-axis. ***$p < 0.001$, **$p < 0.01$, ns, not significant (n=6 experiments, Chi-squared = 7443.7, df = 14, p-value < 2.2e-16).

**c.** Average pumps/minute in *ilc-17.1, cep-1 OE and daf-2* dauers when subjected to control (L4440; Ctrl), *cki-1* and *phg-1* RNAi. n= 4 experiments, and 10 dauers were scored per experiment. *$p < 0.05$, ns, not significant (unpaired t-test). Pumping rates in wild type (N2), L2 larvae are shown for comparison.

Data in all graphs show mean ± S.E.M. Individual points in the bar graphs in **b** represent the % dauers/experiment, Pearson's Chi-squared test with Yates' continuity correction.

Figure S12

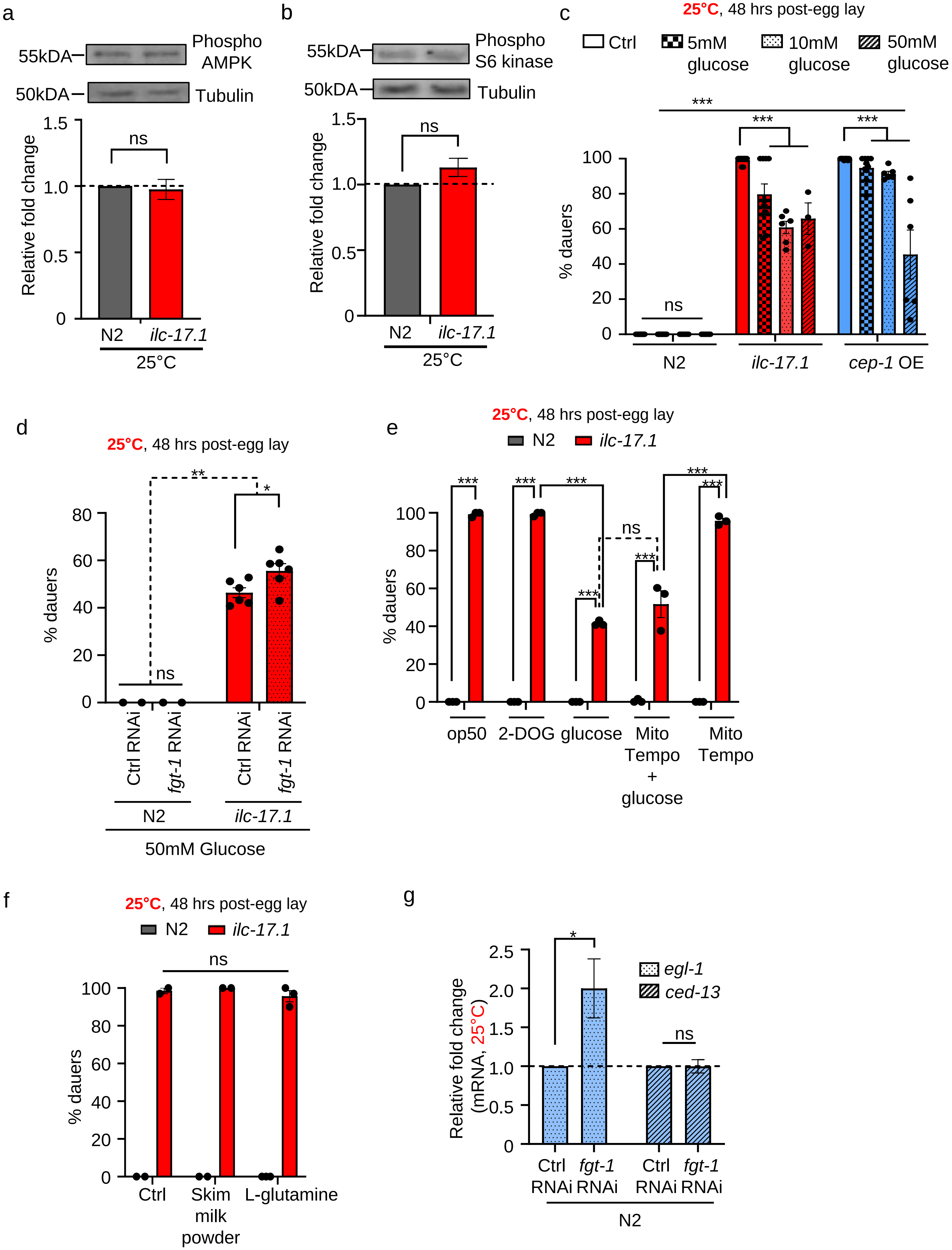
a
55kDA
Phospho AMPK
50kDA
Tubulin
Relative fold change
ns
N2
ilc-17.1
25°C
b
Phospho S6 kinase
Tubulin
c
25°C, 48 hrs post-egg lay
Ctrl
5mM glucose
10mM glucose
50mM glucose
% dauers
cep-1 OE
d
25°C, 48 hrs post-egg lay
Ctrl RNAi
fgt-1 RNAi
50mM Glucose
e
op50
2-DOG
glucose
Mito Tempo + glucose
Mito Tempo
f
Ctrl
Skim milk powder
L-glutamine
g
egl-1
ced-13
Relative fold change (mRNA, 25°C)
Ctrl RNAi
fgt-1 RNAi

**Supplementary Figure S12: CEP-1/p53 activity can be repressed by glucose supplementation.**

**a. Top:** Representative Western blot showing (upper panel) phospho-AMPK levels in *ilc-17.1* deletion mutant larvae and wild type (N2); (lower panel) tubulin, loading control. **Bottom:** Quantification of Phospho-AMPK levels relative to tubulin and normalized to control animals. n=3 experiments. ns, not significant (unpaired t-test).

**b. Top:** Representative Western blot showing (upper panel) phospho-S6 kinase levels in *ilc-17.1* deletion mutant larvae and wild type (N2); (lower panel) tubulin, loading control. **Bottom:** Quantification of Phospho-S6 kinase levels relative to tubulin and normalized to control animals. n=3 experiments. ns, not significant (unpaired t-test).

**c.** Percent dauers amongst *ilc-17.1* deletion mutants, CEP-1/p53 overexpressing larvae and wild type (N2) larvae on 5 mM, 10mM and 50 mM glucose at 25°C. ***p < 0.001, **p < 0.01, ns, not significant. (*n*= 5-9 experiments, Chi-squared = 5263.6, df = 11, p-value < 2.2e-16)

**d.** Percent dauers amongst *ilc-17.1* deletion mutants and wild type (N2) larvae on 50mM glucose on Control (Ctrl; L4440) and *fgt-1* RNAi. **p < 0.01, *p < 0.05, ns, not significant. (*n*=6 experiments, Chi-squared = 299.36, df = 3, p-value < 2.2e-16).

**Extended text 3:** We ruled out non-specific effects of glucose rescue testing the involvement of mitochondrial reactive oxygen species, ROS, known to be increased under glucose-rich diets(97) (Supplementary Fig. S12e): glucose rescue was not diminished upon treatment with mitochondria-targeted antioxidant, Mito-Tempo. Similarly, other supplements such as L-glutamine or skim milk powder (98) did not rescue the dauer arrest (Supplementary Fig. S12f).

**e.** Percent dauers amongst *ilc-17.1* deletion mutants and wild-type (N2) larvae upon exposure to 50mM DOG, 50mM glucose, and 0.1mM MitoTEMPO at 25°C (see X-axis for different conditions). ***$p < 0.001$, ns, not significant, (n=3- experiments, Chi-squared = 1251.7, df = 9, p-value < 2.2e-16). **Associated with Extended text 3.**

**f.** Percent dauers amongst *ilc-17.1* deletion mutants and wild-type (N2) larvae on 20mM Skim Milk powder and 0.26 mM L-Glutamine at 25°C. ns, not significant, (n=3 experiments, Chi-squared = 648.89, df = 5, p-value < 2.2e-16). **Associated with Extended text 3**.

**g.** Average mRNA levels of *egl-1* and *ced-13* in wild-type (N2) larvae subjected to Control (Ctrl; L4440) and *fgt-1* RNAi. mRNA levels were determined relative to *pmp-3* and normalized to wild type (N2) values. *n=3* experiments, *$p < 0.05$, ns, not significant (unpaired t-test).

Data in all graphs show mean ± S.E.M. Individual points in the bar graphs in **c-f** represent the % dauers/experiment, Pearson's Chi-squared test with Yates' continuity correction.

Figure S13

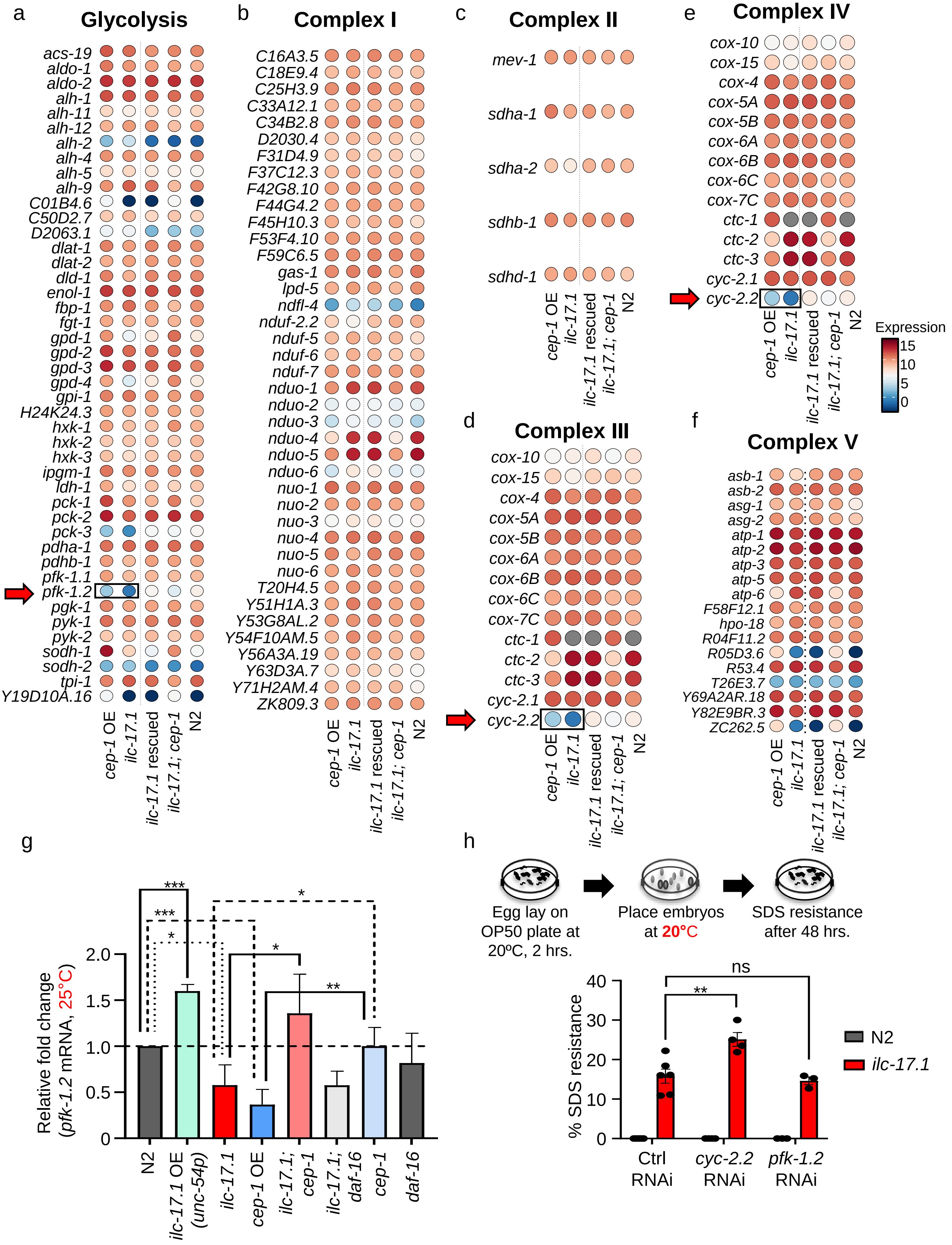

a
Glycolysis
acs-19
aldo-1
aldo-2
alh-1
alh-11
alh-12
alh-2
alh-4
alh-5
alh-9
C01B4.6
C50D2.7
D2063.1
dlat-1
dlat-2
dld-1
enol-1
fbp-1
fgt-1
gpd-1
gpd-2
gpd-3
gpd-4
gpi-1
H24K24.3
hxk-1
hxk-2
hxk-3
ipgm-1
ldh-1
pck-1
pck-2
pck-3
pdha-1
pdhb-1
pfk-1.1
pfk-1.2
pgk-1
pyk-1
pyk-2
sodh-1
sodh-2
tpi-1
Y19D10A.16
cep-1 OE
ilc-17.1
ilc-17.1 rescued
ilc-17.1; cep-1
N2
b
Complex I
C16A3.5
C18E9.4
C25H3.9
C33A12.1
C34B2.8
D2030.4
F31D4.9
F37C12.3
F42G8.10
F44G4.2
F45H10.3
F53F4.10
F59C6.5
gas-1
lpd-5
ndfl-4
nduf-2.2
nduf-5
nduf-6
nduf-7
nduo-1
nduo-2
nduo-3
nduo-4
nduo-5
nduo-6
nuo-1
nuo-2
nuo-3
nuo-4
nuo-5
nuo-6
T20H4.5
Y51H1A.3
Y53G8AL.2
Y54F10AM.5
Y56A3A.19
Y63D3A.7
Y71H2AM.4
ZK809.3
cep-1 OE
ilc-17.1
ilc-17.1 rescued
ilc-17.1; cep-1
N2
c
Complex II
mev-1
sdha-1
sdha-2
sdhb-1
sdhd-1
cep-1 OE
ilc-17.1
ilc-17.1 rescued
ilc-17.1; cep-1
N2
d
Complex III
cox-10
cox-15
cox-4
cox-5A
cox-5B
cox-6A
cox-6B
cox-6C
cox-7C
ctc-1
ctc-2
ctc-3
cyc-2.1
cyc-2.2
cep-1 OE
ilc-17.1
ilc-17.1 rescued
ilc-17.1; cep-1
N2
e
Complex IV
cox-10
cox-15
cox-4
cox-5A
cox-5B
cox-6A
cox-6B
cox-6C
cox-7C
ctc-1
ctc-2
ctc-3
cyc-2.1
cyc-2.2
cep-1 OE
ilc-17.1
ilc-17.1 rescued
ilc-17.1; cep-1
N2
Expression
15
10
5
0
f
Complex V
asb-1
asb-2
asg-1
asg-2
atp-1
atp-2
atp-3
atp-5
atp-6
F58F12.1
hpo-18
R04F11.2
R05D3.6
R53.4
T26E3.7
Y69A2AR.18
Y82E9BR.3
ZC262.5
cep-1 OE
ilc-17.1
ilc-17.1 rescued
ilc-17.1; cep-1
N2
g
Relative fold change (pfk-1.2 mRNA, 25°C)
0
0.5
1.0
1.5
2.0
***
***
*
*
*
**
N2
ilc-17.1 OE (unc-54p)
ilc-17.1
cep-1 OE
ilc-17.1; cep-1
ilc-17.1; daf-16
cep-1
daf-16
h
Egg lay on OP50 plate at 20ºC, 2 hrs.
Place embryos at 20°C
SDS resistance after 48 hrs.
% SDS resistance
0
10
20
30
40
ns
**
Ctrl RNAi
cyc-2.2 RNAi
pfk-1.2 RNAi
N2
ilc-17.1

**Supplementary Figure S13: ILC-17.1 deficiency and CEP-1/p53 overexpression modulate key glucose metabolic enzymes.**

**a-f**. Heatmap depicting expression levels ($log_2$ normalized counts) of glycolysis and mitochondrial OXPHOS genes in the *ilc-17.1* deletion mutants, CEP-1/p53 overexpressing animals, *ilc-17.1; cep-1* double mutants, *ilc-17.1* rescued with *unc-54p::ilc-17.1*, and wild type (N2). Arrows: highlight decreases in *pfk-1.2* and *cyc-2.2* mRNA levels in *ilc-17.1* deleted and CEP-1/p53 overexpressing larvae (Boxed). RNA-seq data was collected as previously described.

**g**. Average *pfk-1.2* mRNA levels in 32-36 hr. old larvae grown at 25°C. mRNA levels were determined relative to *pmp-3* and normalized to wild type (N2) values. Strain names on X-axis. n=4-6 experiments. ***p < 0.001, **p < 0.01, *p < 0.05. (unpaired t-test).

**h. Top:** Schematic of SDS treatment to assess dauer formation at 20°C. **Bottom:** Percent of wild type (N2) and *ilc-17.1* deletion mutant larvae that enter a dauer state at 20°C, 48 hr. post-hatching, when subjected to control (Ctrl; L4440), *cyc-2.2* and *pfk-1.2* RNAi. **p < 0.01, ns, not significant (*n*=3-4 experiments, Chi-squared = 213.38, df = 5, p-value < 2.2e-16 ). Individual points in the bar graphs represent the % dauers/experiment, Pearson's Chi-squared test with Yates' continuity correction.

Data in all graphs show mean ± S.E.M.

### Figure S14

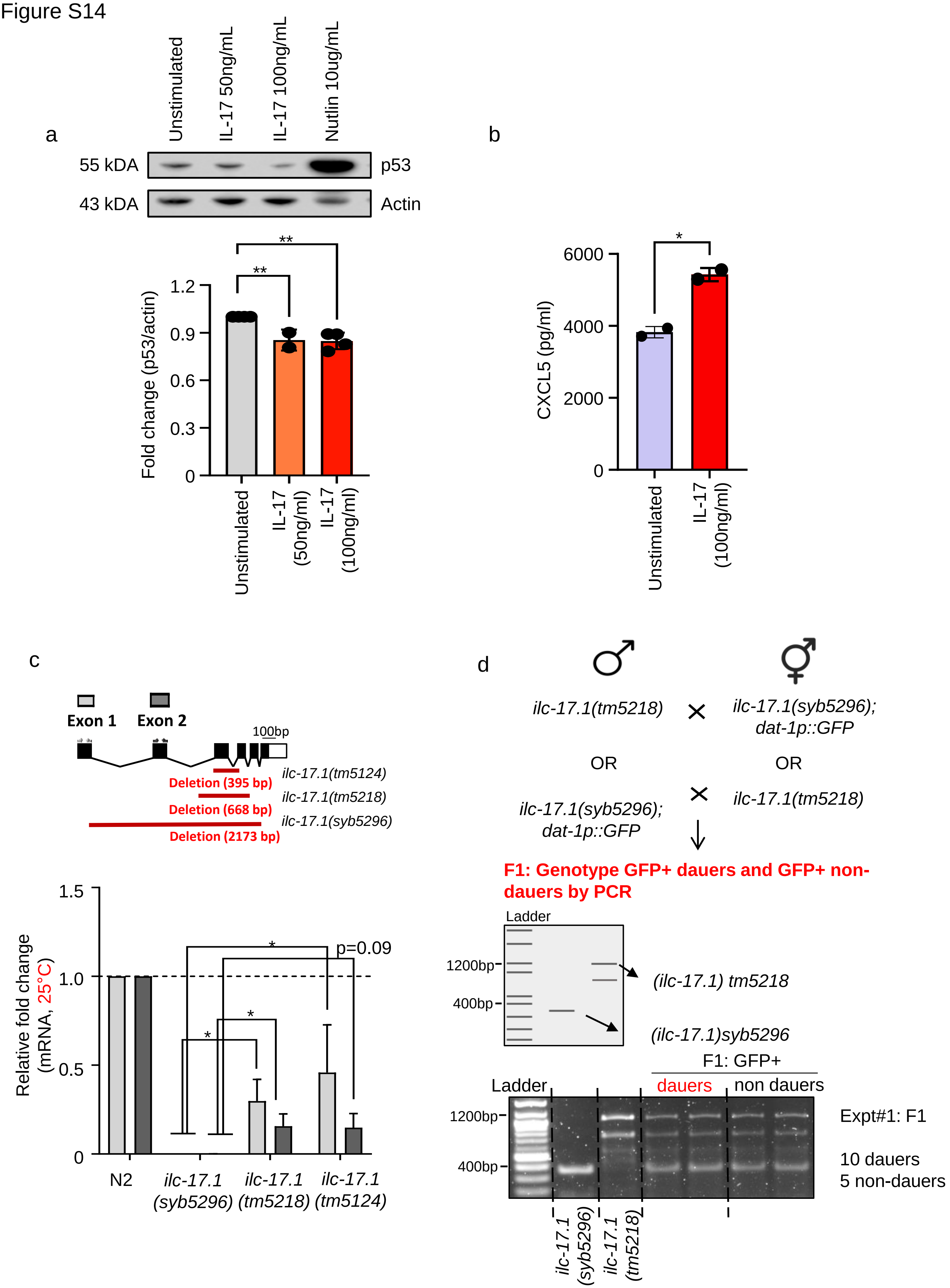
a
Unstimulated
IL-17 50ng/mL
IL-17 100ng/mL
Nutlin 10ug/mL
55 kDA
p53
43 kDA
Actin
**
**
Fold change (p53/actin)
1.2
0.9
0.6
0.3
0
Unstimulated
IL-17 (50ng/ml)
IL-17 (100ng/ml)
b
*
CXCL5 (pg/ml)
6000
4000
2000
0
Unstimulated
IL-17 (100ng/ml)
c
Exon 1
Exon 2
100bp
ilc-17.1(tm5124)
Deletion (395 bp)
ilc-17.1(tm5218)
Deletion (668 bp)
ilc-17.1(syb5296)
Deletion (2173 bp)
Relative fold change (mRNA, 25°C)
1.5
1.0
0.5
0
*
p=0.09
*
*
N2
ilc-17.1 (syb5296)
ilc-17.1 (tm5218)
ilc-17.1 (tm5124)
d
ilc-17.1(tm5218)
ilc-17.1(syb5296); dat-1p::GFP
OR
OR
ilc-17.1(syb5296); dat-1p::GFP
ilc-17.1(tm5218)
F1: Genotype GFP+ dauers and GFP+ non-dauers by PCR
Ladder
1200bp
400bp
(ilc-17.1) tm5218
(ilc-17.1)syb5296
F1: GFP+
Ladder
dauers
non dauers
1200bp
400bp
Expt#1: F1
10 dauers
5 non-dauers
ilc-17.1 (syb5296)
ilc-17.1 (tm5218)

**Supplementary Figure S14: (associated with Extended text 4). rIL-17 induces the expected cytokine response in human epithelial cells.**

**Extended text 4:** Stimulation of untransformed human epithelial cells with human recombinant IL-17A induced a modest but significant downregulation of p53 expression levels (Supplementary Fig. S14a). For these experiments, IL-17A stimulation was confirmed by measurement of secreted CXCL5 (100) (Supplementary Fig. S14b). **Associated with Extended text 4.**

**a. Top:** Representative Western blot showing p53 levels in A459 epithelial cells stimulated for 18hrs. with increasing doses of rIL-17. β-actin was used as a loading control and Nutlin was used as a positive control. **Bottom:** Quantification of p53 levels normalized to actin levels. n= 4 independent experiments. **p<0.01. (One-Way ANOVA with uncorrected Fisher's LSD; df=2). **Associated with Extended text 4.**

**b.** Soluble CXCL5 levels in the supernatant of A459 epithelial cells stimulated for 18hrs. with rIL-17 measured by ELISA. Data is a summary of 2 independent experiments, performed in triplicates. *p<0.05, (unpaired t-test). Data show mean ± S.E.M. **Associated with Extended text 4.**

**c.** Average mRNA levels of exon 1 and 2 in wild-type (N2) larvae and *ilc-17.1(tm5218)X,* and *ilc-17.1(tm5124)X.* mRNA levels were determined relative to *pmp-3* and normalized to wild type (N2) values. *n=3* experiments; 100-200 larvae each. *p<0.05, (unpaired t-test).

**d.** An example experiment, to assess the phenotypes of *ilc-17.1(tm5218)* and *ilc-17.1(syb5296)* trans-heterozygotes. Trans-heterozygotes were generated by mating *ilc-17.1(tm5218)* or *ilc-17.1(syb5296)* males with *ilc-17.1(tm5218)* or *ilc-17.1(syb5296)* hermaphrodites. *ilc-17.1(syb5296)* were crossed into a GFP-expressing background (*dat-*

*1p*::GFP) and progeny were assessed for GFP expression as a means to confirm mating, and cross-progeny were verified by PCR. Approximately ten GFP-positive dauers and GFP-positive non-dauers were harvested for PCR genotyping per experiment (n>10 experiments; 1 experiment consisted of scoring F1 progeny from five mated hermaphrodites from one mating; progeny were grown at 25°C as in **Figure 1a**). An example of PCR results from lysates of four animals (2 dauer; 2 non-dauer adults) is shown. The presence of trans-heterozygous dauers indicates non-complementation. The ratio of **dauer: non-dauer** trans-heterozygotes varied between experiments.
